# Shared functional specialization in transformer-based language models and the human brain

**DOI:** 10.1101/2022.06.08.495348

**Authors:** Sreejan Kumar, Theodore R. Sumers, Takateru Yamakoshi, Ariel Goldstein, Uri Hasson, Kenneth A. Norman, Thomas L. Griffiths, Robert D. Hawkins, Samuel A. Nastase

**Affiliations:** Princeton Neuroscience Institute, Princeton University, Princeton, NJ, 08540, USA; Department of Computer Science, Princeton University, Princeton, NJ 08540, USA; Faculty of Medicine, The University of Tokyo, Bunkyo-ku, Tokyo 113-0033, Japan; Department of Psychology, Princeton University, Princeton, NJ 08540, USA

**Author notes:** Equal contribution. The name listed first in the manuscript was determined by coinflip. Each person reserves the right to put their name first on their CV.

## Abstract

Humans use complex linguistic structures to transmit ideas to one another. The brain is thought to deploy specialized computations to process these structures. Recently, a new class of artificial neural networks based on the Transformer architecture has revolutionized the field of language modeling, attracting attention from neuroscientists seeking to understand the neurobiology of language *in silico*. Transformers integrate information across words via multiple layers of structured circuit computations, forming increasingly contextualized representations of linguistic content. Prior work has focused on the internal representations (the “embeddings”) generated by these circuits. In this paper, we instead analyze the circuit computations directly: we deconstruct these computations into functionally-specialized “transformations” to provide a complementary window onto linguistic computations in the human brain. Using functional MRI data acquired while participants listened to naturalistic spoken stories, we first verify that the transformations account for considerable variance in brain activity across the cortical language network. We then demonstrate that the emergent syntactic computations performed by individual, functionally-specialized “attention heads” differentially predict brain activity in specific cortical regions. These heads fall along gradients corresponding to different layers, contextual distances, and syntactic dependencies in a low-dimensional cortical space. Our findings indicate that large language models and the cortical language network may converge on similar trends of functional specialization for processing natural language.

## Introduction

Language comprehension is a fundamentally constructive process. We resolve local dependencies among words to assemble lower-level linguistic units into higher-level units of meaning (Chomsky, 1965; MacDonald et al., 1994; Partee, 1995; Goldberg, 2006; Berwick et al., 2013; Christiansen & Chater, 2016), ultimately arriving at the kind of narratives we use to understand the world (Bruner, 1985; Graesser et al., 1994). For example, if a speaker refers to “the secret plan,” we implicitly process the relationships between words in this construction to understand that “secret” modifies “plan.” At a higher level, we use the context of the surrounding narrative to understand the meaning of this phrase—*what* does the plan entail, *who* is keeping it secret, and who are they keeping it secret *from*? This context may comprise hundreds of words unfolding over the course of several minutes. The human brain is thought to implement these processes via a series of functionally-specialized computations that transform acoustic speech signals into actionable representations of meaning (Hickok & Poeppel, 2007; Hasson et al., 2015; Ding et al., 2016; Friederici et al., 2017; Martin & Doumas, 2017; Pylkkänen, 2019; Martin, 2020).

Traditionally, neuroimaging research has used targeted experimental manipulations to isolate particular linguistic computations—for example, by manipulating the presence/absence or complexity of a given syntactic structure—and mapped these computations onto brain activity in controlled settings (Bookheimer, 2002; Vigneau et al., 2006; Hickok & Poeppel, 2007; Friederici, 2011; Price, 2012). While these findings laid the groundwork for a neurobiology of language, they have limited generalizability outside the laboratory setting, and it has proven difficult to synthesize them into a holistic model that can cope with the full complexity of natural language. This has prompted the field to move toward more naturalistic comprehension paradigms (Hamilton & Huth, 2020; Nastase et al., 2020; Willems et al., 2020). However, these paradigms introduce their own challenges: principally, how to explicitly quantify the linguistic content and computations supporting the richness and expressivity of natural language (Mitchell et al., 2008; Wehbe et al., 2014; Huth et al., 2016; Pereira et al., 2018).

In recent years, the field of natural language processing (NLP) has been revolutionized by a new generation of deep neural networks capitalizing on the Transformer architecture (Vaswani et al., 2017; Devlin et al., 2019; Radford et al., 2019). Transformers are deep neural networks that forgo recurrent connections (Elman, 1990; Hochreiter & Schmidhuber, 1997) in favor of layered “attention head” circuits, facilitating self-supervised training on massive real-world text corpora. Following pioneering work on word embeddings (Landauer & Dumais, 1997; Mikolov et al., 2013; Pennington et al., 2014), the Transformer architecture represents the meaning of words as numerical vectors in a high-dimensional “embedding” space where closely related words are located nearer to each other. However, while the previous generation of embeddings assign each word a single static (i.e. non-contextual) meaning, Transformers process long sequences of words simultaneously to assign each word a context-sensitive meaning. The core circuit motif of the Transformer—the attention head—incorporates a weighted sum of information exposed by other words, where the relative weighting “attends” more strongly to some words than others. The initial embeddings used as input to the Transformer are non-contextual. Within the Transformer, attention heads in each layer operate in parallel to *update* the contextual embedding, resulting in surprisingly sophisticated representations of linguistic structure (Manning et al., 2020; Linzen & Baroni, 2021).

The success of Transformers has inspired a growing body of neuroscientific work using them to model human brain activity during natural language comprehension (Jain & Huth, 2018; Toneva & Wehbe, 2019; Antonello et al., 2021; Caucheteux et al., 2021a, 2021b, 2021c, 2022; Lyu et al., 2021; Goldstein, Zada, et al., 2022; Heilbron et al., 2022). These efforts have focused exclusively on the “embeddings”—the Transformer’s representation of linguistic *content*—and have largely overlooked the “transformations”—the actual *computations* performed by the attention heads. Although no functionally-specific language modules are built into the architecture at initialization, recent work in NLP has revealed emergent functional specialization in the network after training (Clark et al., 2019; Tenney et al., 2019). That is, particular attention heads are shown to selectively implement interpretable linguistic operations. For example, attention head 10 in the eighth layer of BERT appears to be specialized for resolving the direct object of a verb (e.g. in “the boy in the yellow coat greeted his teacher”, the verb “greeted” attends to “boy”), whereas head 11 in the same layer closely tracks nominal modifiers (e.g. attending to “coat” in the phrase modifying “boy”). Although the individual heads that implement these computations operate independently, in parallel, their transformations are ultimately “fused” together to form the resulting embedding. Thus, unlike the embeddings, the transformations at a given layer can be disassembled into the specialized computations performed by the constituent heads. These transformations are the unique component of the circuit that allows information to flow between words: contextualization of each word’s meaning occurs solely via the transformations. Can we leverage emergent functional specialization within Transformers to better understand linguistic computations in the brain?

In the current work, we argue that the headwise transformations—the functionally-specialized internal computations implemented by individual attention heads—provide a complementary window onto linguistic processing in the brain (Fig. 1A) with respect to the embeddings. Using the widely-studied BERT model (Devlin et al., 2019; Rogers et al., 2020), we extract two internal features of the Transformer architecture (Fig. 1B, C): (1) the layerwise *embeddings* capturing the contextual linguistic content of each word, with contextual information accumulating across successive layers; and (2) the headwise *transformations* capturing the incremental information contributed by the computation at each attention head. We use encoding models to evaluate how well different components of the Transformer predict fMRI data acquired during natural language comprehension. To set the groundwork for our investigation of functional specialization in these headwise transformations, we first compare these Transformer features against two other classes of language features (classical linguistic features and static semantic embeddings), finding that both embeddings and transformations outperform other models. We then show that the transformations recapitulate layer preferences previously observed for embeddings across the language network, although transformations tend to peak at earlier layers than embeddings. Finally, we decompose these transformations into individual attention heads—functionally specialized for particular linguistic operations—and find shared trends of functional specialization in the cortical language network.

**Figure 1.**
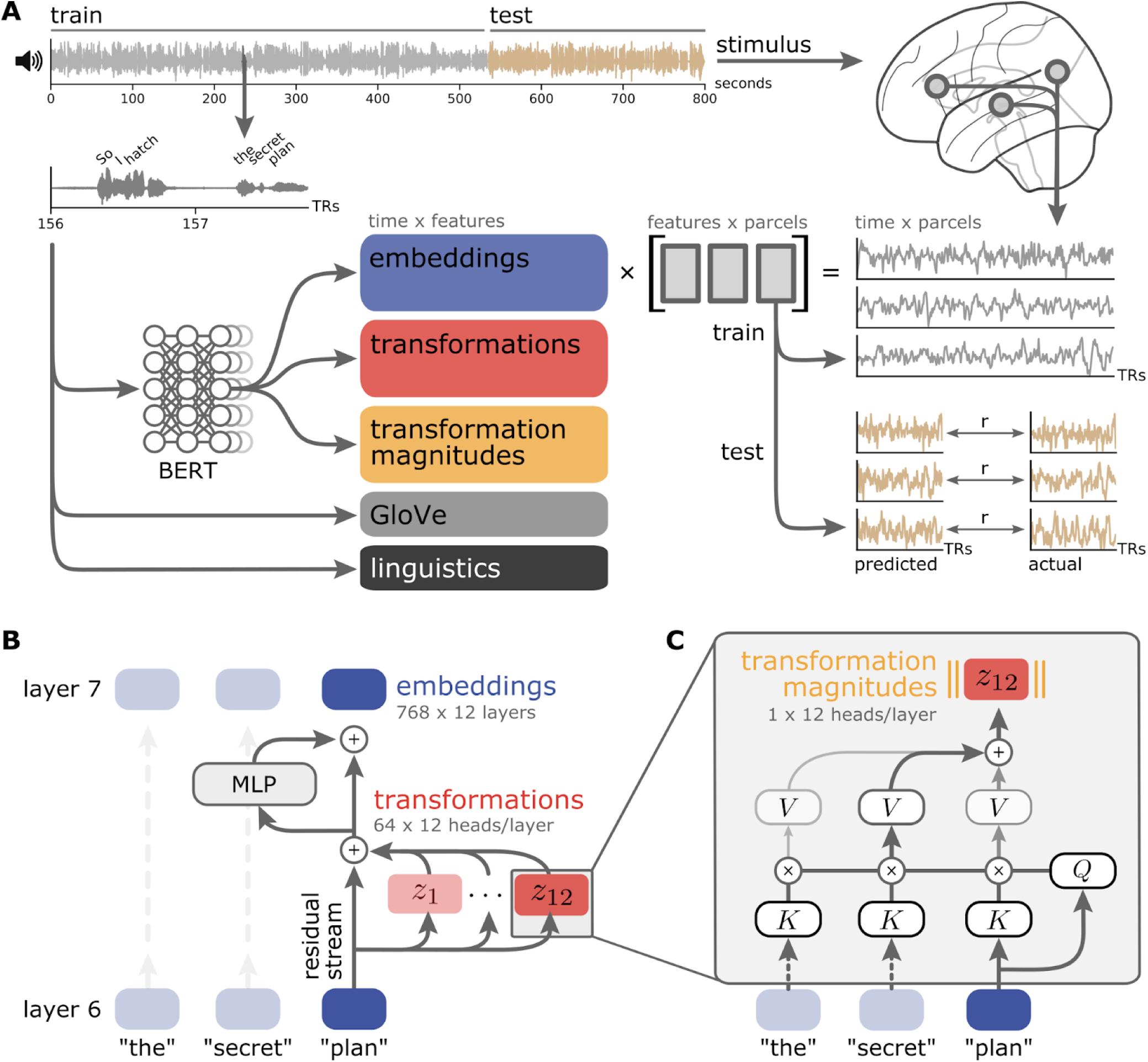
Encoding models for predicting brain activity from the internal components of language models. (**A**) Various features are used to predict parcelwise fMRI time series acquired while subjects listened to naturalistic spoken stories. In addition to classical linguistic features (e.g. parts of speech; black) and static semantic features (e.g. GloVe vectors; gray) extracted from the story transcript, we extracted internal features from a widely-studied Transformer model (BERT-base). The encoding model is estimated from a training subset of each story using banded ridge regression and evaluated on a left-out test segment of each story using three-fold cross-validation. The weights learned from the training set are used to predict the parcelwise time series for the test set; predictions are evaluated by computing the correlation between the predicted and actual parcelwise time series for the test set. (**B**) We consider two core components of the Transformer architecture at each layer (BERT-base and GPT-2 each have 12 layers): the embeddings (blue) and the transformations (red). Embeddings represent the contextualized semantic content of the text. Transformations are the output of the self-attention mechanism for each attention head (BERT-base and GPT-2 have 12 heads per layer, each producing a 64-dimensional vector). Due to the residual connection, transformations represent the incremental information content added to the embedding in that layer. Finally, we also consider the transformation magnitudes (yellow; the L2 norm of each attention head’s 64-dimensional transformation vector), which represent how active each attention head was at that layer. MLP: multilayer perceptron. (**C**) Attention heads use learned matrices to produce content-sensitive updates to each token. Here, we illustrate the process for a single token (“plan”) passing through a single attention head (layer 7, head 12). First, the input token vectors are multiplied by the head’s learned weight matrices (which are invariant across inputs) to produce query (Q), key (K), and value (V) vectors for each input token. The inner product between the query vector for this token (“plan”, Q) and the key vector (K) for each other token yields a set of “attention weights” that describe how relevant the other tokens are to “plan.” These “attention weights” are used to linearly combine the value vectors (V) from the other tokens. The summed output is the transformation for each head (here, Z_12_). The results from each attention head in this layer are concatenated and added back to the token’s original representation.

## Results

We adopted a model-based encoding framework (Naselaris et al., 2011; Yamins & DiCarlo, 2016; Richards et al., 2019) in order to map Transformer features onto brain activity measured using fMRI while subjects listened to naturalistic spoken stories (Fig. 1A). The brain data were spatially downsampled according to a fine-grained functional atlas comprising 1,000 cortical parcels (Schaefer et al., 2018), which were grouped into a variety of regions of interest (ROIs) spanning early auditory cortex to high-level language areas (Fedorenko et al., 2010). Parcelwise encoding models were estimated using banded ridge regression with three-fold cross-validation for each subject and each story (Dupré la Tour et al., 2022). Phonemes, phoneme rate, word rate, and a silence indicator were included as confound variables during model estimation and discarded during model evaluation (Huth et al., 2016). Encoding models were evaluated by computing the correlation between the predicted and actual time series for test partitions; correlations were then converted to the proportion of a noise ceiling estimated via intersubject correlation (Nastase et al., 2019; Fig. S1).

### Functional anatomy of a Transformer model

We take BERT-base (Devlin et al., 2019) as a representative Transformer model due to its success across a broad suite of NLP tasks and the burgeoning literature studying what it learns (Hewitt & Manning, 2019; Liu et al., 2019; DeRose et al., 2020; Hawkins et al., 2020; Hoover et al., 2020; Manning et al., 2020; Rogers et al., 2020; Linzen & Baroni, 2021; Pavlick, 2022).^1^ Words are first assigned non-contextual static embeddings, which are submitted as input to the model. Unlike recurrent architectures (Elman, 1990; Hochreiter & Schmidhuber, 1997), which process word embeddings serially, BERT considers long “context windows” of up to 512 tokens and processes all words in the context window in parallel. Like most Transformer architectures, BERT is composed of a repeated self-attention motif: 12 sequential *layers,* each consisting of 12 parallel *attention heads*. A single *attention head* consists of three separate *query*, *key*, and *value* matrices. At each layer, the input word embeddings are multiplied by these matrices to produce a *query*, *key*, and *value* vector for each word. For an individual word, self-attention then computes the dot product of that word’s *query* vector with the *key* vector from all words in the context window, resulting in a vector of *attention weights* quantifying the relevance of other words. These weights are used to produce a weighted sum of all *value* vectors, which is the output of the attention head. We refer to these 64-dimensional vectors as the “transformation” produced by that attention head. The transformation vectors are then concatenated across all heads within a layer and passed through a feed-forward module (a multilayer perceptron; MLP) to produce a fused 768-dimensional output embedding, which serves as the input for the next layer (Fig. 1B).

Embeddings are sometimes referred to as the “residual stream”: the transformations at one layer are *added* to the embedding from the previous layer, so the embeddings accumulate previous computations that subsequent computations may access (Elhage et al., 2021).

The self-attention mechanism can be thought of as a “soft,” or weighted, key-value store. Each word in the input issues a *query* which is checked against the *keys* of all context words. However, unlike a traditional key-value store (which would only return a single, exact query-key match), the attention head first computes how well the query matches with all keys (the attention weights), and returns a weighted sum of each word’s *value* based on how closely they match. The query, key, and value matrices are learned, and recent work has revealed an emergent functional specialization where specific attention heads approximate certain syntactic relationships (Clark et al., 2019; Tenney et al., 2019). For example, prior work has discovered that a specific head in BERT reliably represents the *direct object* relationship between tokens (Clark et al., 2019). In a phrase such as “hatch the secret plan,” the “plan” token would attend heavily to the “hatch” token and update its representation accordingly. More precisely, **q_plan_^T^ · k_hatch_** will yield a large attention weight from “plan” to “hatch.” As a result, the transformation for “plan,” **z_plan_**, will be heavily weighted towards **v_hatch_** (Eq. 1). This allows the attention head to update the “plan” token’s representation to reflect the fact that it is being “hatched” (as opposed to being executed, revised, or abandoned; or being used in another sense entirely, e.g. an architectural blueprint).

### Transformer-based features outperform other linguistic features

Before disassembling the transformations into specialized circuit computations, we first evaluated how well the transformations, considered in aggregate, perform against other commonly-studied language features in predicting brain activity. We compared the encoding performance of features from three families of language models: (1) traditional linguistic features comprising parts of speech and syntactic dependencies; (2) GloVe word embeddings (Pennington et al., 2014) that capture the “static” or non-contextual meanings of words; and (3) contextualized Transformer features extracted from BERT—namely, layer-wise embeddings, transformations, and transformation magnitudes.^2^ For each TR, we appended the words from the preceding 20 TRs as context.^3^ BERT was allowed to perform bidirectional attention across the tokens in these 21 TRs, after which the context tokens were discarded and the TR tokens were averaged to obtain the Transformer features. To summarize the overall performance of these different Transformer features, we concatenated features across all heads and layers, allowing the regularized encoding model to select the best-performing combination of those features across the entire network. We use these features to predict response time series in cortical parcels comprising ten language ROIs ranging from early auditory cortex to high-level, left-hemisphere language areas (Fig. 2; see Fig. S2 for right-hemisphere results). Based on prior work (Schrimpf et al., 2021; Goldstein, Zada, et al., 2022), we expected the BERT embeddings to outperform the GloVe and linguistic features. We further hypothesized that the set of transformations would perform on par with the embeddings. Finally, the transformation magnitudes, intended to capture the relative contribution of each head, abstracted away from the semantic content, would more closely match the performance of linguistic features.

**Figure 2.**
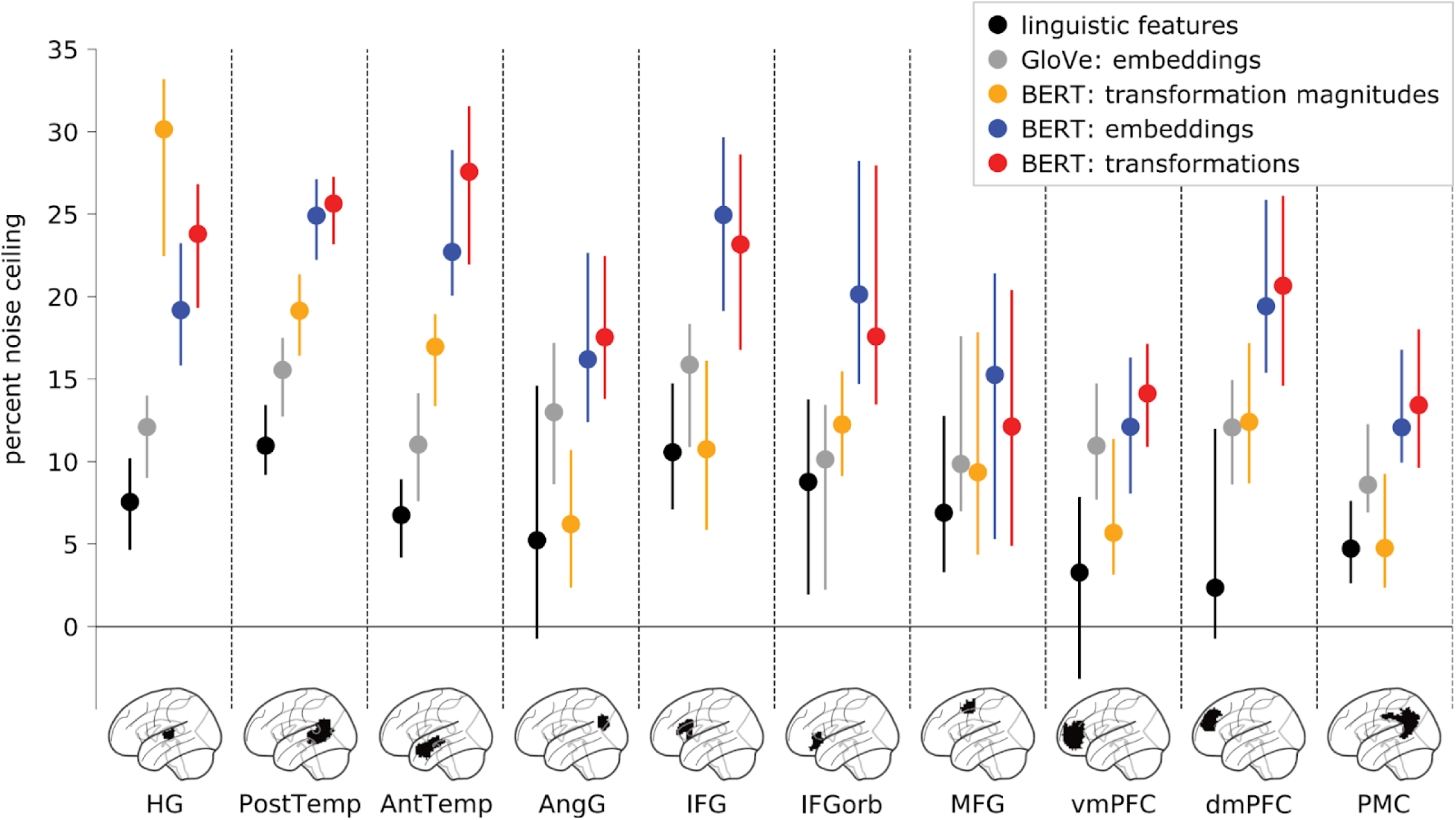
Comparing three classes of language models across cortical language areas. We used encoding models to evaluate the performance of three different classes of language models: traditional linguistic features, non-contextual word embeddings (GloVe), and contextual Transformer features (BERT). Among the Transformer features, embeddings capture the contextual semantic content of words, transformations capture the contextual transformations that yield these embeddings, and transformation magnitudes capture the non-semantic contribution of each head to a given token. For this analysis, Transformer features were concatenated across all heads and layers. Only left-hemisphere language ROIs are included here; right-hemisphere language ROIs yielded qualitatively similar results (Fig. S2). Model performance is evaluated in terms of the percent of a noise ceiling estimated using intersubject correlation (see “Noise ceiling estimation” in “Materials and methods” for further details; Fig. S1). Markers indicate median performance across participants and error bars indicate 95% bootstrap confidence intervals. HG: Heschl’s gyrus; PostTemp: posterior temporal lobe; AntTemp: anterior temporal lobe; AngG: angular gyrus; IFG: inferior frontal gyrus; IFGorb: orbital inferior frontal gyrus; MFG: middle frontal gyrus; vmPFC: ventromedial prefrontal cortex; dmPFC: dorsomedial prefrontal cortex; PMC: posterior medial cortex.

First, we confirmed that Transformer embeddings and transformations outperform traditional linguistic features in most language ROIs (*p* < .005 in HG, PostTemp, AntTemp, AngG, IFG, IFGorb, vmPFC, dmPFC, and PMC for both embeddings and transformations; permutation test; FDR corrected; Table S1). Contextual Transformer embeddings also outperform non-contextual GloVe embeddings across several ROIs in keeping with prior work (Schrimpf et al., 2021; Goldstein, Zada, et al., 2022). Interestingly, transformation magnitudes outperform GloVe embeddings and traditional linguistic features in lateral temporal areas but not in higher-level language areas: for example, transformation magnitudes outperform GloVe embeddings in posterior and anterior temporal areas, but this pattern is reversed in the angular gyrus. Finally, we found that the transformations roughly match the embeddings across all ROIs. This overall pattern of results was replicated in the autoregressive GPT-2 model (Fig. S3). Note that despite yielding similar encoding performance, the embeddings and transformations are fundamentally different; for example, the average TR-by-TR correlation between embeddings and transformations across both stimuli is effectively zero (-.004 ±.009 SD), the embeddings and transformations yield visibly different TR-by-TR representational geometries (Fig. S4), and the transformations have considerably higher temporal autocorrelation than the embeddings (Fig. S5). As a control analysis, we evaluated these features in a non-language ROI (early visual cortex) and found that no models captured a significant amount of variance (Fig. S6). Overall, these findings suggest that the transformations capture a considerable proportion of variance of neural activity across the cortical language network and motivate more detailed treatment of their functional properties.

### Layerwise performance of embeddings and transformations

Based on prior work (e.g. Toneva & Wehbe, 2019), we next segregated the Transformer features into separate layers. Layerwise embeddings are increasingly contextualized, with word representations in later layers reflecting more complex linguistic relationships (Tenney et al., 2019). Transformations, on the other hand, have a very different layerwise structure. While the embeddings generally accumulate information across layers, the transformations are largely independent from layer to layer (Fig. S7) and produce more layer-specific representational geometries (Figs. S8, S9). We found that, across language ROIs, the performance of contextual embeddings increased roughly monotonically across layers, peaking in late-intermediate or final layers (Figs. S10A, S11), replicating prior work (Toneva & Wehbe, 2019; Caucheteux & King, 2022; Caucheteux et al., 2022; Goldstein, Ham, et al., 2022). Interestingly, this pattern was observed across most ROIs, suggesting that the hierarchy of layerwise embeddings does not cleanly map onto the cortical hierarchy for language comprehension. Transformations, on the other hand, seem to yield more layer-specific fluctuations in performance than embeddings and tend to peak at earlier layers than embeddings (Figs. S10B, C, S12).

We next visualized layer preference across cortex—that is, which layer yielded the peak performance for a given cortical parcel (Fig. 3A). Across language parcels, the average performance (across participants) for transformations peaked at significantly earlier layers than performance for embeddings (mean preferred transformation layer = 7.2; mean preferred embedding layer = 8.9; *p* < .001, permutation test; Fig. 3B). Finally, we quantified the magnitude of difference in predictive performance from layer to layer and found that transformations have larger differences in performance between neighboring layers (mean layerwise embedding difference = 7.6, mean layerwise transformation difference = 14.3; *p* < .001, permutation test; Fig. 3C). These results recapitulate the progression of layer specificity reported in the literature (Toneva & Wehbe, 2019; Caucheteux & King, 2022; Caucheteux et al., 2022; Goldstein, Ham, et al., 2022), and suggest that the computations implemented by the transformations are more layer-specific than the embeddings. However, we contend that these layerwise trends provide only a coarse view of functional specialization: individual attention heads perform strikingly diverse linguistic operations even within a given layer.

**Figure 3.**
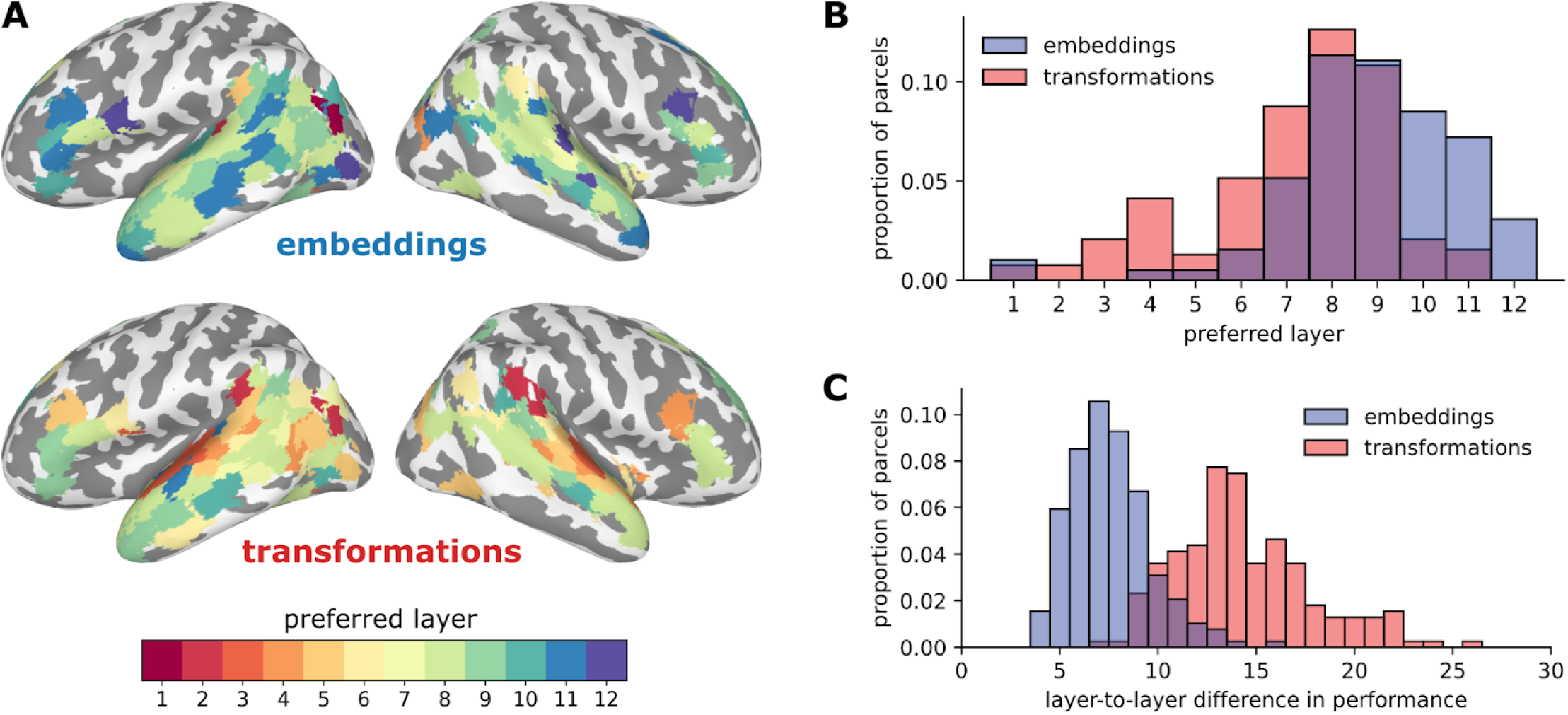
Layer preferences for embeddings and transformations. (**A**) Layer preferences are visualized on the cortical surface for embeddings (upper) and transformations (lower). While most cortical parcels prefer the final embedding layers, the transformations reveal a cortical hierarchy of increasing layer preference. Only cortical parcels with encoding performance greater than 20% of the noise ceiling for both embeddings and transformations are included for visualization purposes. The same color map for preferred layer is used for both embeddings and transformations. (**B**) Histogram of the preferred layer across cortical parcels in language ROIs. Across language parcels, performance for transformations (red) peaks at intermediate layers, while performance for embeddings (blue) peaks in later layers. (**C**) Distribution of the magnitude of layer-to-layer differences in encoding performance for embeddings and transformations; this metric of layer specificity is quantified as the L2 norm of the first differences between encoding performance for neighboring layers. Transformations (red) yield more layer-specific deviations in performance than embeddings (blue).

### Interpreting transformations via headwise analysis

Does the emergent functional specialization of internal computations in the language model reflect functional specialization observed in the cortical language network? To begin answering this core question, we first directly examined how well classical linguistic features—indicator variables identifying parts of speech and syntactic dependencies—map onto cortical activity. Despite a large body of work using experimental manipulations of phrases and sentences to dissociate syntax from semantics and localize syntactic operations in the brain (Dapretto & Bookheimer, 1999; Embick et al., 2000; Kuperberg et al., 2000; Ni et al., 2000; Friederici et al., 2003; Glaser et al., 2013; Schell et al., 2017), results have been mixed, leading some authors to suggest that syntactic computations may be fundamentally entangled with semantic representation and distributed throughout the language network (Fedorenko et al., 2012, 2020; Blank et al., 2016; Reddy & Wehbe, 2020; Caucheteux et al., 2021c). Along these lines, we found that classical linguistic features are poor predictors of brain activity and do not provide a good basis for examining functional specialization in the cortical language network in the context of naturalistic narratives (Figs. 2, S13).

Unlike the hand-crafted syntactic features of classical linguistics, BERT’s training regime yields an emergent headwise functional specialization for particular linguistic operations (Clark et al., 2019; Tenney et al., 2019). BERT is not explicitly instructed to represent syntactic dependencies, but nonetheless learns functionally comparable operations from the structure of real-world language (Clark et al., 2019). We split the transformations at each layer into their functionally specialized components—the constituent transformations implemented by each attention head. In the following analyses, we leverage these functionally-specialized headwise transformations to map between syntactic operations and the brain in a way that retains some level of interpretability, but respects the contextual nature of real-world language. Note that the embeddings incorporate information from all the transformations at a given layer (and prior layers), and therefore cannot be meaningfully disassembled in this way. We trained an encoding model on all transformations and then evaluated the prediction performance for each head individually, yielding an estimate of how well each head predicts each cortical parcel (or *headwise brain prediction* score). For each attention head, we also trained a set of decoding models to determine how much information that head contains about a given syntactic dependency (or *headwise dependency prediction* score; Fig. S14).

For each classical syntactic dependency, we first identified the attention head that best predicts that dependency (for example, head 11 of layer 6 best predicts the direct object dependency, Fig. S13). We compared the encoding performance for each classical dependency with the encoding performance for the headwise transformation that best predicts that dependency. We found that the head most associated with a given dependency generally outperformed the dependency itself (Fig. S13; see Fig. S15 replication using functionally specialized heads derived from larger text corpora; Clark et al., 2019). This confirmed our expectation that the dense, emergent headwise transformations are better predictors of brain activity than the sparse, classical linguistic indicator variables. Note that the headwise transformations are considerably higher-dimensional (64 dimensions) than the corresponding one-dimensional dependency indicators. However, we found that even after reducing a given transformation to a single dimension that best predicts the corresponding dependency, the one-dimensional transformation still better predicts brain activity than the dependency itself (Fig. S16). We found that these one-dimensional transformation time series are highly correlated with the corresponding dependency indicators, but reflect a continuous, graded representation of the dependency over the course of a narrative (Fig. S17). Although the computations performed by these heads approximate particular syntactic operations, they capture a more holistic relationship between words in the context of the narrative. That is, the transformations do not simply indicate the presence, for example, of a direct object relationship; rather, they capture an approximation of the direct object relationship *in the context* of the ongoing narrative.

Critically, BERT does not *just* learn to reproduce classical syntactic operations; it learns a rich multiplicity of linguistic and contextual relations from natural language, a subset of which can be said to approximate classical syntactic labels (Rogers et al. 2020). With this in mind, we pursued a data-driven analysis to summarize the contributions of all headwise transformations across the entire language network (Fig. 4A). We first obtained the trained encoding model for all transformations (Fig. 1, red) and averaged the regression coefficients (i.e. weight matrices) assigned to the transformation features across subjects and stimuli. To summarize the importance of each head for a given parcel, we segmented the learned weight matrix from the encoding model for that parcel into the individual attention heads at each layer and computed the L2 norm of the headwise encoding weights. This results in a single value for each of the 144 heads reflecting the magnitude of each head’s contribution to encoding performance at each parcel; these vectors capture each parcel’s “tuning curve” across the attention heads. In order to summarize the contribution of headwise transformations across the language network, we aggregated the headwise encoding weight vectors across parcels in language ROIs and used principal component analysis (PCA) to reduce the dimensionality of this transformation weight matrix across parcels, following Huth and colleagues (2016). This yields a basis set of orthogonal (uncorrelated), 144-dimensional weight vectors capturing the most variance in the headwise transformation weights across all language parcels; each head corresponds to a location in this low-dimensional brain space. The first two principal components (PCs) accounted for 92% of the variance in weight vectors across parcels, while the first nine PCs accounted for 95% of the variance. A given PC can be projected into (i.e. reconstructed in) the original space of cortical parcels, yielding a brain map where positive and negative values indicate positive and negative transformation weights along that PC (Fig. S18). Visualizing these PCs directly on the brain reveals that PC1 ranges from strongly positive values (red) in bilateral posterior temporal parcels and left lateral prefrontal cortex to widespread negative values (blue) in medial prefrontal cortex (Fig. 4B). PC2 ranged from positive values (red) in prefrontal cortex and left anterior temporal areas to negative values (blue) in partially right-lateralized temporal areas (Fig. 4C). Note that the polarity of these PCs is consistent across all analyses, but is otherwise arbitrary.

**Figure 4.**
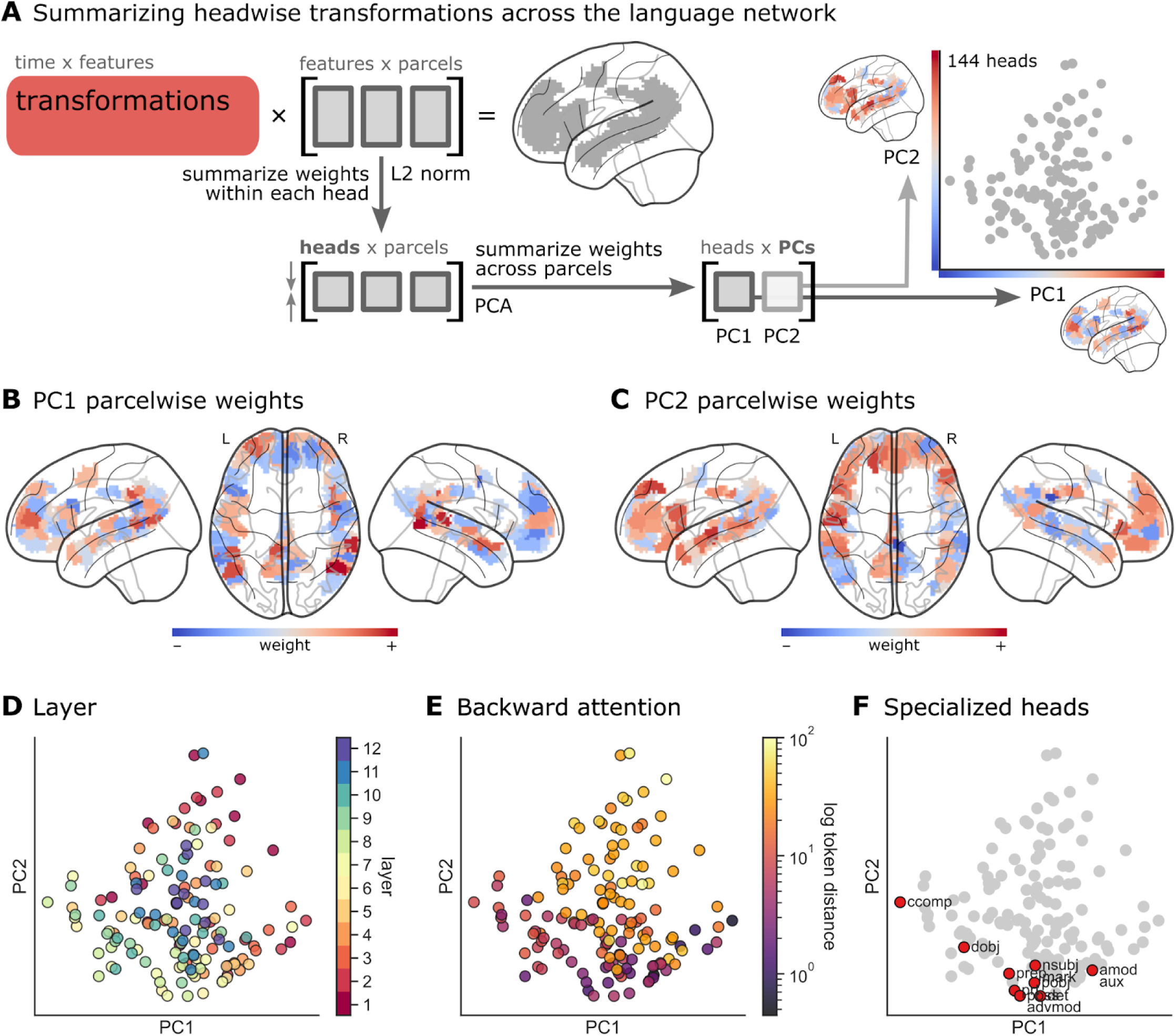
Headwise transformations in a low-dimensional brain space. (**A**) We applied PCA to the weight vectors for transformation encoding models across language parcels, effectively projecting the transformation weights into a low-dimensional brain space (Huth et al., 2016). We first obtain the parcelwise weight vectors for the encoding model trained to predict brain activity from BERT transformations. This transformation weight matrix is shaped (768 features * 12 layers) 9,216 features × 192 language parcels. We use the L2 norm to summarize the weights within each head, reducing this matrix to (12 heads * 12 layers) 144 heads × 192 language parcels. We next summarize these headwise weights across language parcels using PCA. At right, we visualize the headwise transformation weights projected onto the first two PCs. Each data point corresponds to one of 144 heads. Furthermore, each PC can be projected back onto the language network (see Fig. S22 for a control analysis). (**B**, **C**) PC1 and PC2 projected back onto the language parcels; red indicates positive weights and blue indicates negative weights along the corresponding PC. (**D**) Heads colored according to their layer in BERT in the reduced-dimension space of PC1 and PC2. (**E**) Heads colored according to their average backward attention distance in the story stimuli (look-back token distance is colored according to a log scale). (**F**) Heads highlighted in red have been reported as functionally specialized by Clark and colleagues (Clark et al., 2019).

We next examined whether there is any meaningful structure in the “geometry” of headwise transformations in this reduced-dimension cortical space. To do this, we visualized several structural and functional properties of the heads in two-dimensional projections of the language network. We first visualized the *layer* of each head in this low-dimensional brain space and found a layer gradient across heads in PC1 and PC2 (Fig. 4D). PCs 9, 5, and 1 were the PCs most correlated with layer assignment with *r* = .45, .40, and .26, respectively (Figs. S18, S19). Intermediate layers were generally located in the negative quadrant of both PC1 and PC2 (corresponding to blue parcels in Fig. 4B, C; e.g. posterolateral temporal cortex), with early and late layers located in the positive quadrant (red parcels). For each head, we next computed the *average backward attention distance* across stimuli. PCs 2, 1, and 3 were the PCs most correlated with backward attention distance with *r* = .65, .20, .19, respectively (Figs. S18, S20). We observed a strong gradient of look-back distance increasing along PC2 (Fig. 4E); that is, prefrontal and left anterior temporal parcels (red parcels in Fig. 4C) correspond to heads with longer look-back distances. Note that the upper quartile of headwise attention distances exceeds 30 tokens, corresponding to look-back distances on the scale of multiple sentences. We also found that the functionally specialized heads previously reported in the literature (Clark et al., 2019) span PC1 and cluster at the negative end of PC2 (corresponding to intermediate layers and relatively recent look-back distance; Fig. 4F). Finally, we visualized the headwise dependency prediction scores in this low-dimensional brain space and observed gradients in different directions along PC1 and PC2 for different dependencies (Fig. S21). Note that none of the aforementioned structural or functional properties of the heads that we visualize in this low-dimensional brain space are derived from the brain data; that is, the encoding models do not “know” the *layer* or *backward attention distance* of any given head.

As a control analysis, we shuffled the coefficients assigned to transformation features by the encoding model across heads within each layer of BERT. We then repeated the same analysis: we segmented the shuffled transformation features back into “pseudo-heads,” computed the L2 norm of the coefficients within each pseudo-head, and applied PCA across language parcels. This perturbation disrupts the emergent functional grouping of transformation features into particular heads observed in the unperturbed model. After this perturbation, the first two PCs accounted for only 17% of variance across language parcels (reduced from 92% in the unperturbed model). PCs were dramatically less correlated with layer assignment (maximum *r* across PCs reduced from .45 to .25) and look-back distance (maximum *r* across PCs reduced from .65 to.26). Finally, this perturbation abolished any visible geometry of layer, look-back distance, or headwise dependency decoding in the low-dimensional projection onto PCs 1 and 2 (Fig. S22). This control analysis indicates that the structure observed in Fig. 4 results from the grouping of transformation features into functionally specialized heads; transformation features map onto brain activity in a way that systematically varies head by head, and shuffling features across heads (even within layers) disrupts this structure (Fig. S23). We observed similar trends in GPT-2 (Fig. S23); interestingly, however, look-back distance was most highly correlated with PC1 (*r* = 0.77) and layer was highly correlated with PC3 (*r* = 0.60).

Finally, to quantify the correspondence between the syntactic information contained in a given head and that head’s prediction performance in the brain, we computed the correlation, across attention heads, between the brain prediction and dependency prediction scores (Fig. 5A, B). We repeated this analysis for each syntactic dependency and the parcels comprising each language ROI (Fig. S24). Headwise correspondence between dependencies and ROIs indicates that attention heads containing information about a given dependency also tend to contain information about brain activity for a given ROI—thus linking that ROI to the computation of that dependency. We found that the correspondence between brain prediction and dependency prediction scores varied considerably across ROIs. For example, in posterior superior temporal cortex, we observed headwise prediction correspondence for clausal complement (*ccomp*), direct object (*dobj*), and preposition object (*pobj*) relations. In IFG, on the other hand, headwise prediction correspondence was observed only for the clausal complement relation. Headwise prediction correspondence was high in the angular gyrus and MFG across dependencies: for example, attention heads that predict the existence of a nominal subject relationship (*nsubj)* also tend to predict the MFG, but not the dmPFC; heads that predict direct object (*dobj*) tend to predict the angular gyrus, but this relationship is weaker in the vmPFC. In the case of MFG, this is consistent with prior work implicating MFG in both language comprehension and more general cognitive demand (e.g. working memory; Fedorenko et al., 2011; Mineroff et al., 2018). Collapsing across dependencies highlights the discrepancy between ROIs (Fig. 5C). The angular gyrus and MFG display a relatively high correspondence; in contrast, the vmPFC and dmPFC display virtually no correspondence. While transformations explain significant variance in these ROIs at the scale of the full model (Fig. 2), individual layers (Fig. S10), and individual heads (Fig. 5A), their prediction performance for the brain does not correlate with their prediction performance for classic syntactic dependencies—suggesting that the shared information between transformations and certain ROIs may be semantic in nature or reflect contextual relationships beyond the scope of classical syntax. Finally, as a control analysis, we shuffled the transformation features across heads within each layer of BERT and then performed the same functional correspondence analysis.

**Figure 5.**
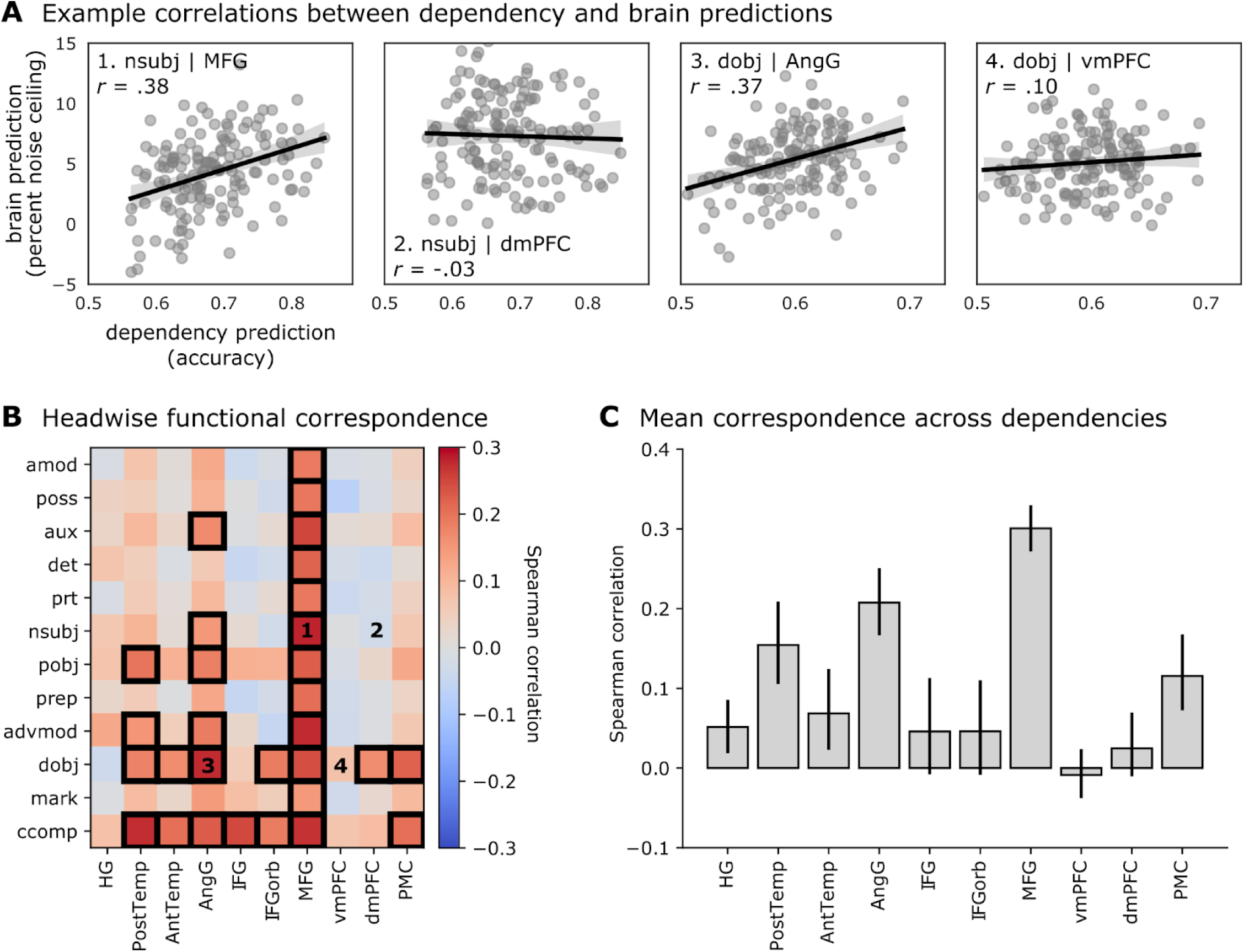
Correspondence between headwise brain and dependency predictions. (**A**) Correlation between headwise brain prediction and dependency prediction scores for example ROIs and dependencies; nominal subject (nsubj) in MFG and dmPFC, direct object (dobj) in AngG and vmPFC (see Fig. S24 for correlations plotted for all ROIs and dependencies). Each point in the scatter plot represents the dependency prediction (x-axis) and brain prediction (y-axis) scores for each of the 144 heads. Brain prediction scores reflect cross-validated encoding model performance evaluated in terms of the percent of a noise ceiling estimated using intersubject correlation. Dependency prediction scores reflect the classification accuracy of a cross-validated logistic regression model trained to predict the occurrence of a given linguistic dependency at each TR from the 64-dimensional transformation vector for a given attention head. Each of these plots corresponds to a labeled cell in the dependencies-by-ROI correlation matrix in panel B. (**B**) Correlation between headwise brain prediction and dependency prediction scores for each language ROI and syntactic dependency. Dependencies (y-axis) are ordered by their token distance; e.g. the adjectival modifier (amod) spans fewer tokens on average than the clausal complement (ccomp; see “Materials and methods” for details). Cells with black borders contain significant correlations as determined by a permutation test in which we shuffle assignments between headwise dependency prediction scores and brain prediction scores across heads (FDR controlled at p < .05). Labeled cells correspond to the example correlations in panel A. (**C**) We summarize the brain–dependency prediction correspondence for each ROI by averaging across syntactic dependencies (i.e. averaging each column of panel B).

Again, perturbing the functional grouping of transformation features into heads reduced both brain and dependency prediction performance and effectively abolished the headwise correspondence between dependencies and language ROIs (Fig. S25). Finally, replicating this analysis in GPT-2 produced higher correspondence values, particularly in IFG, but with less specificity across ROIs (Fig. S26). Note that GPT-2 and BERT have different architectures and different training regimes; given that the models have similar encoding performance overall (Fig. S3), the observed differences in functional correspondence highlight the sensitivity of our headwise analytic framework.

## Discussion

While a growing body of work has used the Transformer architecture (Vaswani et al., 2017; Devlin et al., 2019) to model the neural basis of human language processing (Toneva & Wehbe, 2019; Schrimpf et al., 2021; Caucheteux & King, 2022; Caucheteux et al., 2022; Goldstein, Zada, et al., 2022; Heilbron et al., 2022), we contribute a novel perspective on *how* to relate these models to human brain activity. Prior work has focused on the Transformer’s cumulative representations of linguistic content (the embeddings); we instead interrogate the finer-grained computations implemented by functionally-specialized internal components of the model (the transformations). These transformations are the component of the model that allows information to flow between words: all syntactic, compositional, and contextual relations *across* words are generated by transformations at one or more attention heads. Critically, unlike the embeddings, we can segregate the transformations into the computations performed by separate, functionally-specialized heads. Examining the contribution of headwise transformations to encoding performance reveals gradients in a low-dimensional cortical space that reflect structural and functional properties of the headwise transformations, including layer, look-back distance, and predictive performance on syntactic dependencies. Finally, we quantified the correspondence between headwise predictions of brain activity and syntactic dependencies for a variety of cortical language areas and dependencies, and found that certain ROIs (e.g. PostTemp, AngG, MFG) display strong headwise prediction correspondence across a variety of dependencies. This suggests that BERT and the brain converge on similar trends of functional specialization.

To build an intuition for why the transformations may provide complementary insights to the embeddings, we can compare Transformers to convolutional neural networks (CNNs) commonly used in visual neuroscience (Kriegeskorte, 2015; Yamins & DiCarlo, 2016). CNNs typically reduce dimensionality across layers (Krizhevsky et al., 2012; He et al., 2016), putting pressure on the model to gradually discard task-irrelevant, low-level information and retain only high-level semantic content. In contrast, popular Transformer architectures maintain the same dimensionality across layers. Thus Transformer embeddings can *aggregate* information (from context words) across layers, such that later layers tend to contain the most information (Tenney et al., 2019).^4^ In this light, it is unsurprising that encoding performance tends to peak at later embedding layers.

Indeed, unlike the structural correspondence between CNN layers and the visual processing hierarchy (Güçlü & van Gerven, 2015; Yamins & DiCarlo, 2016; Dupré la Tour et al., 2021), Transformer embeddings are highly predictive but relatively uninformative for localizing stages of language processing. Unlike the embeddings, the transformations reflect *updates* to word meanings at each layer. Encoding models based on the transformations must “choose” a step in the contextualization process, rather than “have it all” by simply using later layers. Our finding that these transformations are equally predictive overall (Fig. 2) but display a higher layer specificity across language areas (Fig. 3) suggests that brains and Transformers may follow a more similar overall pattern of processing than was previously believed—a finding which was potentially obfuscated by prior work’s focus on the embeddings.

Although Transformers perform remarkably well on NLP tasks, they are not positioned to serve as implementational models of cortical circuitry. Nonetheless, the human language system and end-to-end deep language models do—to some extent—share common computational principles aimed at producing well-formed, context-sensitive linguistic outputs (Hasson et al., 2020; Goldstein, Zada, et al., 2022). In both cases, some level of functional specialization appears to emerge (McClelland et al., 2010; Nasr et al., 2019; Yang et al., 2019; Dobs et al., 2022), likely reflecting the statistical structure of natural language. That said, the syntactic operations formalized by classical linguistics may not directly correspond to particular neural computations, and these computations may not be localized to macroanatomical cortical circuits (Mesulam et al., 2015; Blank et al., 2016). In practice, we found that classical linguistic features (i.e. parts of speech and syntactic dependencies) are generally poor predictors of brain activity during natural language comprehension (Figs. 2, S13, S16).

With this in mind, we adopted a data-driven approach to describe the properties of the transformations predominant in brain activity during naturalistic language processing. The distance attention heads tend to “look back” in the narrative (Fig. 4, PC2), and to a lesser extent layer assignment (Fig. 4, PC1, accounted for considerable variance in the mapping between the headwise transformation and the cortical language network. We found that the functional properties of the headwise transformations do, in fact, map onto certain cortical localization trends previously reported in the literature. For example, analyzing the encoding weights for transformations (Fig. 4) revealed that posterior temporal areas assign higher weights to heads at earlier layers (positive values along PC1) with shorter look-back distance (negative values along PC2), consistent with previous work suggesting that posterior temporal areas perform early-stage syntactic (and lexico-semantic) processing (Hickok & Poeppel, 2000, 2007; Flick & Pylkkänen, 2020; Murphy et al., 2022). Headwise correspondence in posterior temporal cortex was high for the *ccomp* and *dobj* dependencies (Fig. 5), which are involved in resolving the meaning of verb phrases, corroborating prior work implicating posterior temporal areas in verb–argument integration (Ben-Shachar et al., 2003; Friederici et al., 2003; Bornkessel et al., 2005). We also found that left-lateralized anterior temporal and prefrontal cortices were associated with longer look-back attention distances (positive values along PC2; Fig. 4), suggesting that these regions may compute longer-range contextual dependencies, including event- or narrative-level relations (Maguire et al., 1999; Vandenberghe et al., 2002; Ferstl et al., 2008; Makuuchi et al., 2009; Bašnáková et al., 2014; Baldassano et al., 2018). Interestingly, the IFG (pars opercularis and triangularis; i.e. Broca’s area) was not strongly associated with heads specialized for particular syntactic operations (Fig. 5B, C), despite being well-predicted by both BERT embeddings and transformations (Fig. 2). There are several possible explanations for this: (1) the natural language stimuli used here may not contain sufficient syntactic complexity to tax IFG; (2) the cortical parcellation used here may yield imprecise functional localization of IFG (Braga et al., 2020; Fedorenko & Blank, 2020); and (3) the IFG may be more involved in language production than comprehension (Matchin & Hickok, 2020).

Despite the formal distinction between syntax and lexico-semantics in linguistics, the neural computations supporting human language may not so cleanly dissociate syntactic and semantic processing (Fedorenko et al., 2020), especially during natural language comprehension where syntax and semantics are typically intertwined. Indeed, although Transformer models implicitly learn syntactic and compositional operations in order to produce well-formed linguistic outputs, these emergent structures are generally entangled with semantic content (Baroni, 2020; Manning et al., 2020; Linzen & Baroni, 2021). While the transformations used in the current analysis capture syntactic and contextual operations entangled with semantic content, the *transformation magnitudes* can serve to disentangle syntax and semantics. Prior work has sought to isolate the two using artificial stimuli and vector subtraction on the embeddings (Caucheteux et al., 2021c); the transformation magnitudes instead reduce the transformations down to the “activation” of individual attention heads. Insights from NLP (Clark et al., 2019) suggest this metric, which circumvents the stimulus representation entirely, nonetheless contains an emergent form of syntactic information. Our comparison to classical linguistic features (Fig. 2) suggests this is the case: transformation magnitudes outperform classical linguistic features in temporal language areas, and perform comparably elsewhere. Interestingly, the transformation magnitudes also outperform the non-contextual word embeddings in temporal areas (Fig. 2), while this relationship is reversed in angular gyrus, a putative high-level convergence zone for semantic representation (Binder et al., 2009).

It is important to note that our analysis is constrained by the temporal resolution of fMRI, which is not sufficient to capture language processing that occurs on rapid timescales. Rather, current fMRI results provide complementary insights to measurement modalities with higher temporal resolution (Goldstein, Zada, et al., 2022; Heilbron et al., 2022) because the superior spatial coverage of fMRI allows us to map language encoding throughout the entire brain. Our findings join a growing body of work demonstrating that fMRI is sensitive to slower-evolving contextual and narrative features that occur in natural language (Lerner et al., 2011; Hasson et al., 2015; Chang et al., 2022).

Our results suggest several future lines of research. Prior work has explored different Transformer architectures (Schrimpf et al., 2021; Caucheteux & King, 2022) aiming to establish a structural mapping between Transformers and the brain. Toward this end, training “bottlenecked” Transformer models that successively reduce the dimensionality of linguistic representations—similar to CNNs—may produce more hierarchical embeddings and provide a better structural mapping onto cortical language circuits (Raghu et al., 2021). Second, the current work sidesteps the acoustic and prosodic features of natural speech (Santoro et al., 2014; de Heer et al., 2017); future work, however, may benefit from models that extract high-level contextual semantic content directly from the speech signal (in the same way that CNNs operate directly on pixel values; Li et al., 2022; Millet et al., 2022; Vaidya et al., 2022). Third, a large body of prior work has utilized Transformer embeddings to predict the brain; it may be worth revisiting some of these findings with an emphasis on transformations instead. Finally, current neurobiological models of language highlight the importance of long-range fiber tracts connecting the nodes of the language network (Catani et al., 2005; Saur et al., 2008; Dick & Tremblay, 2012); we suspect that future language models with more biologically-inspired circuit connectivity may provide insights not only into functional specialization but also functional integration across specialized modules.

Transformer-based deep language models like BERT and GPT obtain state-of-the-art performance on multiple NLP tasks. BERT’s attention heads are functionally specialized and learn to approximate classical syntactic operations in order to produce contextualized natural language (Clark et al., 2019; Tenney et al., 2019). The rapidly developing field of BERTology (Rogers et al., 2020) seeks to characterize this emergent functional specialization. In this work, we took a first step towards bridging BERTology’s insights to language processing in the brain. Although we do not find a direct one-to-one mapping between attention heads, linguistic dependencies, and cortical areas, our findings suggest that certain trends in functional organization—such as a gradient of increasing contextual look-back distance—may be shared. Mapping the internal structure of deep language models to cortical language circuits can bring us closer to a mechanistic understanding of human language processing, and may ultimately provide insights into how and why this kind of functional specialization emerges in both deep language models and the brain.

## Materials and methods

### Experimental data

Models were evaluated on two story datasets from the publicly available “Narratives” collection of fMRI datasets acquired while subjects listened to naturalistic spoken stories (Nastase et al., 2021). Code used to analyze the data are available at the accompanying GitHub repository: https://github.com/tsumers/bert-brains. The “Slumlord” and “Reach for the Stars One Small Step at a Time” dataset includes 18 subjects (ages: 18–27 years, mean age: 21 years, 9 reported female) and comprises two separate stories roughly 13 minutes (550 TRs) and 2,600 words in total. The “I Knew You Were Black” dataset includes 45 subjects (ages: 18-53 years, mean age: 23.3 years, 33 reported female); the story is roughly 13 minutes (534 TRs) long and contains roughly 1,500 words. All functional MRI datasets were acquired with a 1.5 s TR (Nastase et al., 2021), and were organized in compliance with the Brain Imaging Data Structure (Gorgolewski et al., 2016).

Preprocessed MRI data were obtained from the Narratives derivatives release (Nastase et al., 2021). Briefly, the following preprocessing steps were applied to the functional MRI data using fMRIPrep (Esteban et al., 2019): susceptibility distortion correction (using fMRIPrep’s fieldmap-less approach), slice-timing correction, volume registration, and spatial normalization to MNI space (*MNI152NLin2009cAsym* template). Confound regression was then implemented using AFNI’s 3dTproject (Cox, 1996) with the following confound regressors: six head motion parameters, the first five aCompCor components from CSF and from white matter (Behzadi et al., 2007), cosine bases for high-pass filtering (cutoff: 128 s), and first- and second-order polynomial trends. Non-smoothed functional data were used for all analyses in the current study. To harmonize datasets with differing spatial resolution and reduce computational demands, we resampled all functional data to a fine-grained 1000-parcel cortical parcellation derived from intrinsic functional connectivity (Schaefer et al., 2018). That is, time series were averaged across voxels within each parcel to yield a single average response time series per parcel (within each subject and story dataset).

### Baseline language features

Language model representations were derived from the time-locked phoneme- and word-level transcripts available in the Narratives dataset (Nastase et al., 2021). Words were assigned to the fMRI volumes based on the ending timestamps; e.g. if a word began in one TR and ended in the following TR, it was assigned to the second TR. The following low-level acoustic and linguistic features were extracted for each TR to serve as confound variables in subsequent analyses: (*a*) the number of words per TR, (*b*) number of phonemes per TR, and (*c*) a binary vector indicating the presence of individual phonemes per TR. These are the same confound variables used in Huth et al. 2018.

We extracted part-of-speech and dependency relations to serve as classical linguistic features. These features were annotated using the *en_core_web_lg* (v2.3.1) model from spaCy (v2.3.7; Honnibal et al., 2020). For each TR, we created a binary feature vector across all parts-of-speech/dependency relations, indicating whether or not a given part-of-speech/dependency relation appeared within that TR. Part of speech (e.g. noun, verb, adjective) describes the function of each word in a sentence. We used 14 part-of-speech labels: pronoun, verb, noun, determiner, auxiliary, adposition, adverb, coordinating conjunction, adjective, particle, proper noun, subordinating conjunction, numeral, and interjection (Table S3). A dependency relation describes the syntactic relation between two words. For each word, a parser defines another word in the same sentence, called the “head,” to which the word is syntactically related; the dependency relation describes the way the word is related to its head. We used 25 dependency relations: *nsubj*, *ROOT*, *advmod*, *prep*, *det*, *pobj*, *aux*, *dobj*, *cc*, *ccomp*, *amod*, *compound*, *acomp*, *poss*, *xcomp*, *conj*, *relcl*, *attr*, *mark*, *npadvmod*, *advcl*, *neg*, *prt*, *nummod*, and *intj* (Table S4; https://github.com/clir/clearnlp-guidelines/blob/master/md/specifications/dependency_labels.md). For visualization, we focus on 12 dependencies with particularly high correspondence to particular heads reported by Clark and colleagues (Clark et al., 2019).

Finally, to serve as a baseline for Transformer-based language models, we used GloVe vectors (Pennington et al., 2014) which capture the “static” semantic content of a word across contexts. Conceptually, GloVe vectors are similar to the vector representations of text input to BERT prior to any contextualization applied by the Transformer architecture. We obtained GloVe vectors for each word using the *en_core_web_lg* model from spaCy, and averaged vectors for multiple words occurring within a TR to obtain a single vector per TR.

### Transformer self-attention mechanism

While language models such as GloVe (Pennington et al., 2014) assign a single “global” or “static” embedding (i.e. meaning) to a given word across all contexts, the Transformer architecture (Vaswani et al., 2017) introduced the *self-attention* mechanism which yields context-specific representations. Just as convolutional neural nets (LeCun et al., 1998; Krizhevsky et al., 2012) use convolutional filters to encode spatial inductive biases, Transformers use self-attention blocks as a sophisticated computational motif or “circuit” that is repeated both within and across layers. Self-attention represents a significant architectural shift from sequential processing of language via recurrent connections (Elman, 1990) to simultaneously processing multiple tokens. Variations on the Transformer architecture with different dimensionality and training objectives currently dominate major tasks in NLP, with BERT (Devlin et al., 2019) and GPT (Radford et al., 2019) being two of the most prominent examples.

Self-attention operates as follows. A single *attention head* consists of three separate (learned) matrices: a *query* matrix, a *key* matrix, and a *value* matrix, each of dimensionality *d_model_* × *d_head_*, where *d_model_* indicates the dimensionality of the model’s embedding layers, and *d_head_* indicates the dimensionality of the attention head. Input word vectors are multiplied by each of these three matrices independently, producing a *query*, *key*, and *value* vector for each word. To determine the contextualized representation for a given word vector, the self-attention operation takes the dot product of that word’s *query* vector with the *key* vector from all words. The resulting values are then scaled and softmaxed, producing the “attention weights” for that word.

Formally, for a Transformer head of dimensionality *d_head_* and sets of query, key, and value vectors forming matrices **Q**, **K**, **V**, the self-attention mechanism operates as follows:

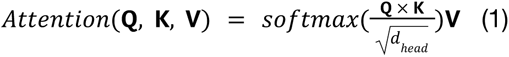

The *i*th token “attends” to tokens based on the inner product of its query vector **Q**_i_ with the key vectors for all tokens, **K**. When the query vector matches a given key, the inner product will be large; the softmax ensures the resulting “attention weights” sum to one. These attention weights are then used to generate a weighted sum of the value vectors, **V**, which is the final output of the self-attention operation (Eq. 1). We refer to the attention head’s output as the “transformation” produced by that head. Each attention head produces a separate transformation for each input token.

State-of-the-art Transformer architectures scale up the core self-attention mechanism described above in two ways. First, multiple attention heads are assembled in parallel within a given layer (“multi-headed attention”). For example, BERT-base-uncased (Devlin et al., 2019), used in most of our analyses, contains 12 attention heads in each layer. The 12 attention heads are each 64-dimensional and act in parallel to produce independent 64-dimensional “transformation” vectors. These vectors are concatenated to produce a single 768-dimensional vector which is then passed to the feed-forward module. Second, BERT-base consists of 12 identical layers stacked on top of each other, allowing the model to learn a hierarchy of transformations for increasingly complex contextualization (Tenney et al., 2019). The full model has 12 layers × 12 heads = 144 total attention heads and a total of 110 million learned parameters. The original input vectors are analogous to the static GloVe vectors; however, as they pass through successive layers in the model, repeated applications of the self-attention mechanism allow their meanings to contextualize each other.

The phenomenal success of Transformer-based models has generated an entire sub-field of NLP research, dubbed “BERTology,” dedicated to reverse-engineering their internal representations (Manning et al., 2020; Rogers et al., 2020). Researchers have linked different aspects of the model to classic NLP concepts, including functional specialization of individual “attention heads” to syntactic dependencies (Clark et al., 2019), representational subspaces corresponding to parse trees (Hewitt & Manning, 2019), and an overall layerwise progression of representations that parallels traditional linguistic pipelines (Tenney et al., 2019). Our approach builds on this work, using internal Transformer features as a bridge from these classical linguistic concepts to brain data.

### Transformer-based features

We used three separate components from the Transformer models to predict brain activity. The first features we extract are the layerwise “embeddings,” which are the de facto Transformer feature used for most applications, including prior work in neuroscience (e.g. Schrimpf et al., 2021). The embeddings represent the contextualized semantic content, with information accumulating across successive layers as the Transformer blocks extract increasingly nuanced relationships between tokens (Tenney et al., 2019). As a result, embeddings have been characterized as a “residual stream” that the attention blocks at each layer “write” to and “read” from. Later layers often represent a superset of information available in earlier layers, while the final layers are optimized to the specific pretraining task (for BERT, causal language modeling). Prior work using such models typically finds that the mid-to-late layers best predict brain activity (Schrimpf et al., 2021; Caucheteux & King, 2022).

The second set of features we extract are the headwise “transformations” (Eq. 1), which capture the contextual information introduced by a particular head into the residual stream prior to the feedforward layer (MLP). These features are the unique component of the transformer responsible for propagating information between different tokens (see Elhage et al., 2021). Consequently, transformations play a privileged role in constructing meaning from the intricate relationships between words: any contextual information that passes into a single word’s representation is incorporated by way of transformations computed at some head (Meng et al., 2022). In other words, it is simultaneously the earliest *dense* representation to result from the attention operation at a given layer (unlike the sparse attention matrices themselves) and the final *head-wise* representation (unlike MLP activations, which operate on the full residual stream after each head’s transformations have been concatenated back together). While the transformations represent the most crucial component of the self-attention mechanism, to our knowledge they have not been previously studied in human neuroscience.

Finally, the third set of features we extract are the “transformation magnitudes,” which are the L2 norm of each attention head’s transformation. This is effectively a “univariate” metric for how *active* each attention head was: how much its update “moved” the word representation, without any information about the direction of its influence. Because the transformation magnitudes lack direction, they cannot encode any semantic information: it is not possible to discern *how* a word’s meaning was changed, only *how much* it was changed. Prior work has shown that individual attention heads learn to implement particular syntactic relationships (Clark et al., 2019); therefore, the transformation magnitudes provide information about the relationships between tokens in the TR divorced from the semantic content of the TR.

### Generating Transformer features

Transformer models were accessed via the HuggingFace library (Wolf et al., 2020). We used the BERT-base-uncased model. We generated “embeddings” and “transformations” as follows. For each TR, we concatenated words assigned to that TR (3.75 words on average) together with words from the preceding 20 TRs (corresponding to the preceding 30 seconds of the auditory story stimulus). We passed the resulting set of words through the Transformer tokenizer and then model. This procedure allowed information from the preceding time window to “contextualize” the meaning of the words occurring in the present TR. At each layer, we extracted the “transformations” (output of the self-attention submodule, Eq. 1) and the “embeddings” (output of the final feedforward layer) for *only* the words in that TR. We excluded the automatically appended “SEP” token. For each TR, this left us with a tensor of dimension *n_layers_* × *d_model_* × *n_tokens_*, where *n_layers_* indicates the number of layers in the Transformer model, *d_model_* indicates the dimensionality of the model’s embedding layers, and *n_tokens_* the number of tokens spoken in that TR. For BERT, this resulted in a 12 layer × 768 dimension × *n_tokens_* tensor. To reduce this to a consistent dimensionality, we averaged over the tokens occurring within each TR, resulting in a 12 × 768 × 1 tensor for each TR. TRs with no words were assigned zeros. Finally, to generate “transformation magnitudes,” we computed the L2 norm of each headwise transformation.

To generate the “backwards attention” metric (Fig. 5), we followed a procedure similar to the “attention distance” measure in (Vig & Belinkov, 2019). Unlike the previous analyses, this required a fixed number of Transformer tokens per TR. Rather than using the preceding 20 TRs, we first encoded the entire story using the Transformer tokenizer, and for each TR selected the 128 tokens preceding the end of the TR. This corresponded to approximately 30 seconds of the auditory story stimulus. We processed each TR and extracted each head’s matrix of token-to-token attention weights (Eq. 1). We selected the token-to-token attention weights corresponding to information flow from earlier words in the stimulus into tokens in the present TR.^5^ We multiplied each token-to-token attention weight by the distance between the two tokens, and divided by the number of tokens in the TR to obtain the per-head attention distance in that TR. Finally, we averaged this metric over all TRs in the stimulus to obtain the headwise attention distances.

### Decoding dependency relations

In addition to using the linguistic features to predict brain activity, we used the “transformation” representations to predict dependencies. For each TR, the “transformation” consists of a *n_layers_* × *d_model_* tensor; for BERT, this yields a 12 layers × 768 dimension tensor. Each of the layers is thus a 768-dimensional vector, which itself consists of 12 concatenated 64-dimensional vectors, each corresponding to the output of a single attention head. These 64-dimensional headwise vectors were used as predictors. Note that this decomposition is only valid for transformations, which are the direct results of the multi-head attention mechanism; due to the feedforward layer after the multi-head attention mechanism, there is no trivial way to reduce embeddings to headwise representations.

We performed logistic regression with the L2 penalty (implemented using scikit-learn; Pedregosa et al., 2011) to predict the occurrences of each binary dependency relation over the course of each story from the headwise transformations. The regularization hyperparameter was determined for each head and each dependency relation using nested five-fold cross-validation over a log-scale grid with 11 values ranging from 10^-30^ to 10^30^. Since some dependency relations are relatively rare, labels are imbalanced. We corrected for this imbalance by weighting samples according to the inverse frequency of occurrence during training and by using balanced accuracy for evaluation (Brodersen et al., 2010).

### Encoding model estimation and evaluation

Encoding models were estimated using banded ridge regression with three-fold cross-validation (Dupré la Tour et al., 2022). Ridge regression is a formulation of regularized linear regression using a penalty term to ensure that the learned coefficients minimize the squared L2 norm of the model parameters. This effectively imposes a multivariate normal prior with zero mean and spherical covariance on the model parameters (Nunez-Elizalde et al., 2019). Relative to ordinary least squares, ridge regression tends to better accommodate collinear parameters and better generalize to unseen data (by reducing overfitting). In banded ridge regression, the full model is composed of several concatenated submodels including both features of interest (e.g. BERT features) and confound variables. The following confound variables were included based on prior work (Huth et al., 2016): phoneme and word rate per TR, a 32-dimensional phoneme indicator matrix, and an indicator vector indicating silent TRs. Each of these submodels was assigned to a separate “band” in the banded ridge regression and thus received a different regularization parameter when fitting the full model.

Specifically, we have separate bands for: the main model (e.g. BERT features), the silence indicator vector, word count and phoneme count vectors, and the phoneme indicator matrix. When evaluating the model, we generate predictions using only the main model band, discarding any confound features. For each band, we duplicated and horizontally stacked the feature space four times comprising lags of 1, 2, 3, and 4 TRs (1.5, 3.0, 4.5, 6.0 s) in order to account for parcelwise variation in hemodynamic lag (Huth et al., 2016). The regularization parameters were selected via a random search of 100 iterations. In each iteration, we sample regularization parameters for each band uniformly from the simplex by sampling from a dirichlet distribution (as implemented in the *himalaya* package; Dupré la Tour et al., 2022). We performed nested three-fold cross-validation within each training set of the outer cross-validation partition. Encoding models were fit for each of the 1,000 parcels within each subject and each story dataset. The cross-validation procedure was implemented so as to partition the time series into temporally contiguous segments. When conducting the headwise encoding analyses (Fig. 6), we also discard the learned coefficients corresponding to all heads except for the particular head of interest for prediction and evaluation on the test set (Lee Masson & Isik, 2021).

Encoding model performance was evaluated by computing the Pearson correlation between the predicted and actual time series for the test partition. Correlation was used as the evaluation metric in both the nested cross-validation loop for regularization hyperparameter optimization, and in the outer cross-validation loop. For each partition of the outer cross-validation loop, the regularization parameter with the highest correlation from the nested cross-validation loop within the training set was selected. This procedure yields a correlation value for each test set of the outer cross-validation loop for each parcel and subject. These correlation values were then averaged across cross-validation folds and Fisher-z transformed prior to statistical assessment.

### Noise ceiling estimation

The raw correlation values described above depend on the signal-to-noise ratio (SNR), duration, and other particular features of the data. In order to provide a more interpretable metric of model performance, we compute the proportion of the correlation value relative to a noise ceiling—effectively the proportion of explained variance relative to the total variance available. Acquiring replicates of brain responses to the same stimulus in the same subjects is effectively impossible under naturalistic contexts due to repetition effects; i.e. listening to a narrative for a second time does not engage the same cognitive processes as listening to the narrative for the first time (Aly et al., 2018). To circumvent this challenge, we used intersubject correlation (ISC) as an estimate of the noise ceiling (Nili et al., 2014; Nastase et al., 2019). In this approach, time series from all subjects are averaged to derive a surrogate model intended to represent the upper limit of potential model performance. For each subject, the test time series for each outer cross-validation fold is first averaged with the test time series of all other subjects in the dataset, then the test time series for that subject is correlated with the average time series. Including each subject in the average biases the noise ceiling upward, thus yielding more conservative estimates of the proportion. Note that under circumstances where individual subjects are expected to vary considerably in their functional architecture, ISC may provide a suboptimal noise ceiling. However, in the current context, we do not hypothesize or model any such differences.

### Statistical assessment

To assess the statistical significance of encoding model performance, we used two nonparametric randomization tests. First, when testing whether model performance was significantly greater than zero, we used a one-sample bootstrap hypothesis test (Hall & Wilson, 1991). For each iteration of the bootstrap, we randomly sampled subject-level correlation values (averaged across cross-validation folds), then computed the Fisher-transformed mean across the bootstrap sample of subjects to construct a bootstrap distribution around the mean model performance value across subjects for each parcel. We then subtracted the observed mean performance value from this bootstrap distribution, thus shifting the mean roughly to zero (the null hypothesis). Finally, we computed a one-sided *p*-value by determining how many samples from the shifted bootstrap distribution exceeded the observed mean.

Second, when comparing performance between two models, we used a permutation test. For each iteration of the permutation test, we took the subject-wise differences in performance between the two models, randomly flipped their signs, then recomputed the mean difference in correlation across subjects to populate a null distribution. We then computed a two-sided *p*-value by determining how many samples from either tail of the distribution exceeded the mean observed difference.

Statistical tests for population inference were performed by concatenating subjects across story datasets prior to randomization in order to produce one *p*-value across stories; however, randomization was stratified within each story. When assessing *p*-values across parcels or ROIs, we corrected for multiple tests by controlling the false discovery rate (FDR) at *q* < .05 (Benjamini & Hochberg, 1995).

### Summarizing headwise transformation weights

To summarize the contribution of headwise transformations across the language network, we first obtained the regression coefficients—i.e. weight matrices—from the encoding model trained to predict brain activity from the BERT transformations concatenated across layers (Fig. 1, red; corresponding to the performance in Fig. 2, red). We averaged the parcelwise weight vectors across both subjects and stimuli. We next computed the L2 norm of the regression coefficients within each head at each layer, summarizing the contribution of the transformation at each head for each parcel. Following Huth and colleagues (Huth et al., 2012, 2016), we then used principal component analysis (PCA) to summarize these headwise transformation weights across all parcels in the language ROIs. This yields a reduced-dimension brain space where each data point corresponds to the transformation implemented by each of the 144 attention heads. To visualize the structure of these headwise transformations in the reduced-dimension brain space, we colored the data points according to structural and functional properties of the heads, including their *layer*, *backward attention distance*, and *dependency prediction scores*.

## Acknowledgments and funding sources

S.K. was supported by NIH T32MH065214 for the duration of this work. T.S. is supported by an NDSEG Fellowship. R.H. is supported by a C.V. Starr Fellowship. S.A.N. is supported by NIH R01MH112566.

## Supporting information

**Figure S1.**
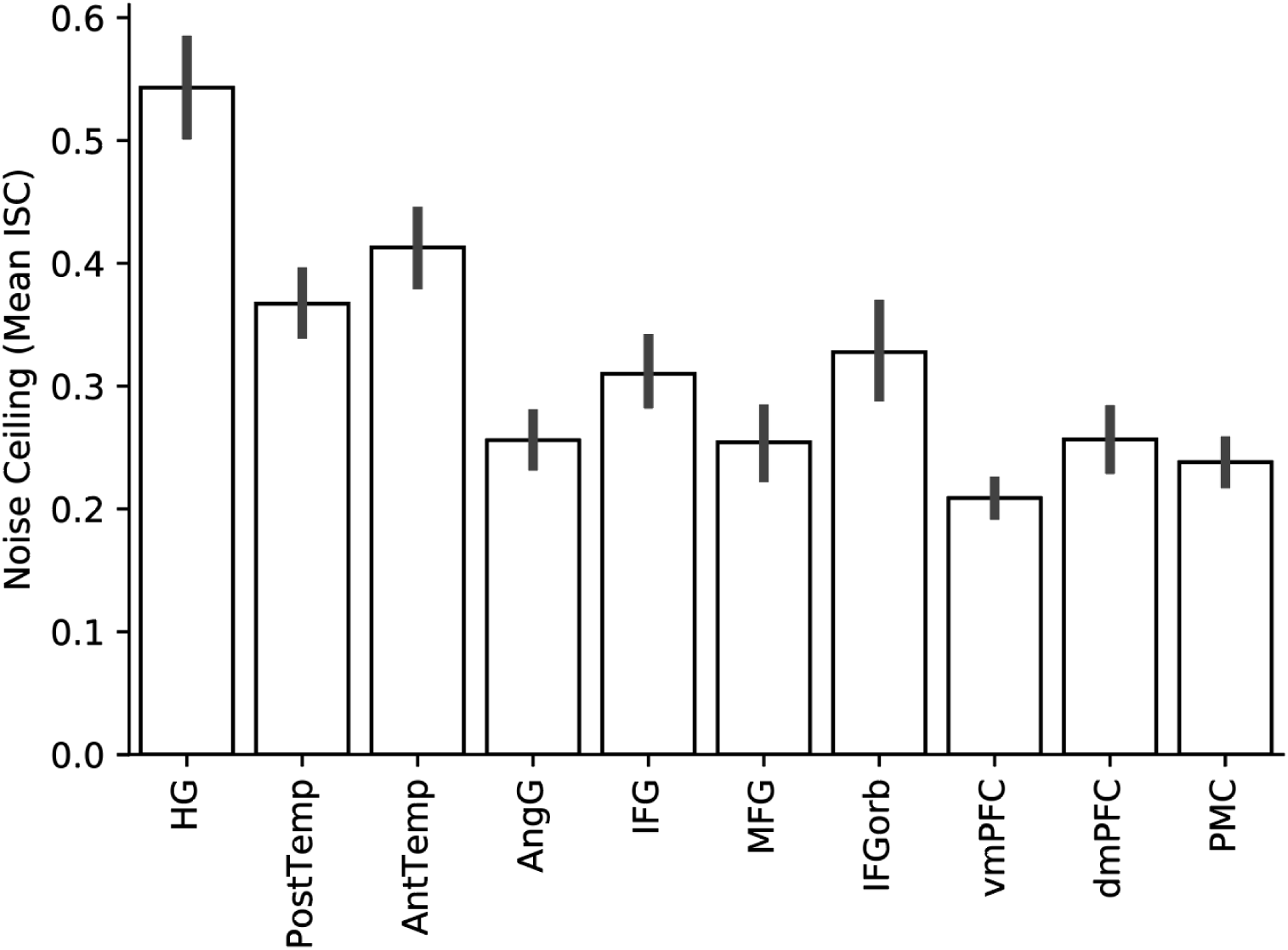
Noise ceilings for each language ROI as measured by intersubject correlation (ISC) analysis (Nastase et al., 2019; cf. Fig. 2). We evaluate the performance of the encoding models by computing the correlation between predicted and actual parcel time series for left-out test segments of each stimulus. To construct a noise ceiling, for each parcel we compute the correlation between each subject’s time series and the average time series of the subjects (i.e. leave-one-out ISC). ISC is computed for test segments of each stimulus to match the inputs to the encoding analysis. Here, we report the mean ISC across subjects for each language ROI; error bars represent 95% bootstrap confidence intervals of the mean across subjects.

**Figure S2.**
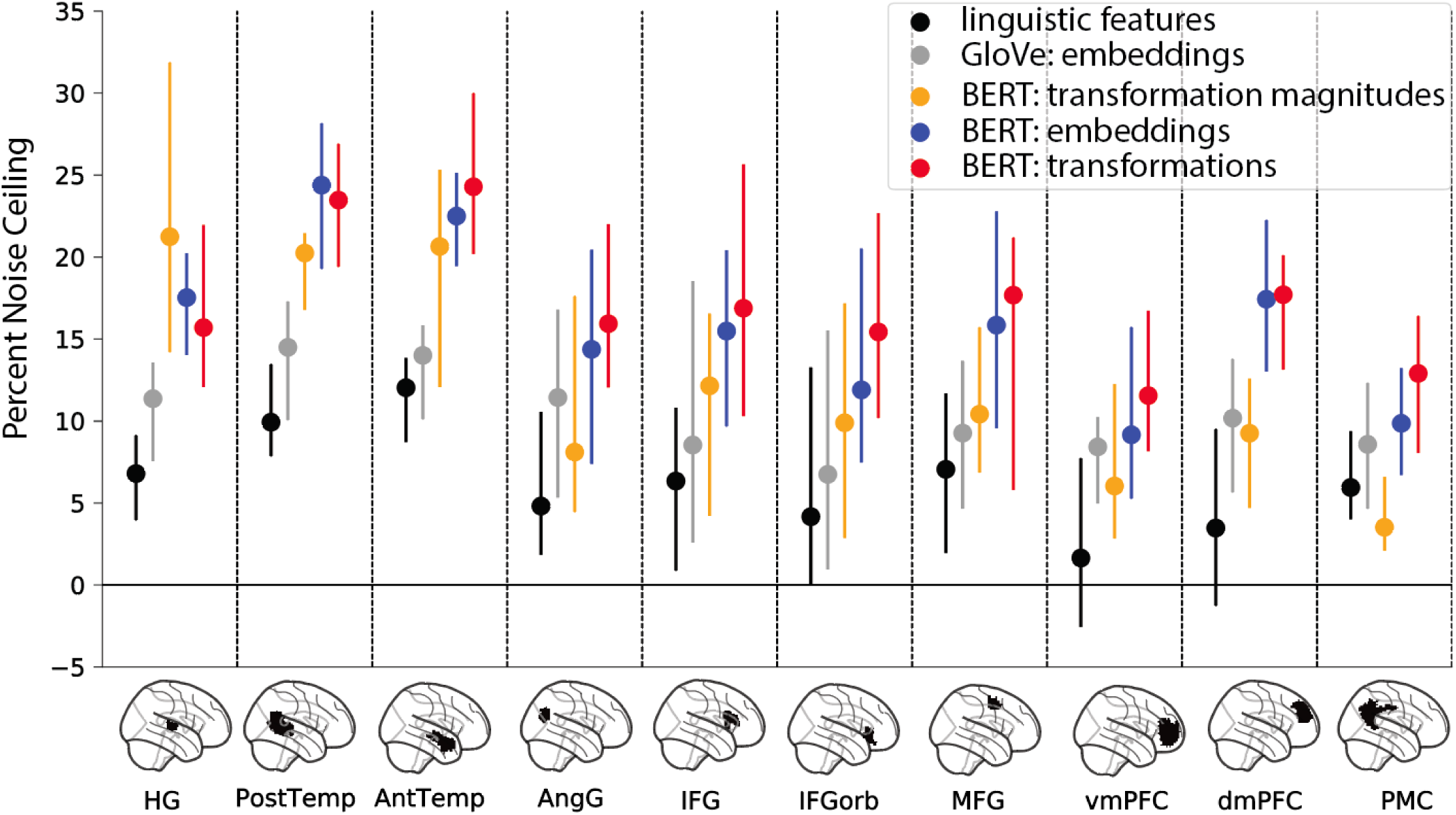
Comparing three classes of language models in right-hemisphere language areas (cf. Fig. 2). Right-hemisphere language areas yield qualitatively similar encoding results to left-hemisphere language areas. Model performance is evaluated in terms of the percent of a noise ceiling estimated using intersubject correlation. Markers indicate median performance and error bars indicate 95% bootstrap confidence intervals.

**Figure S3.**
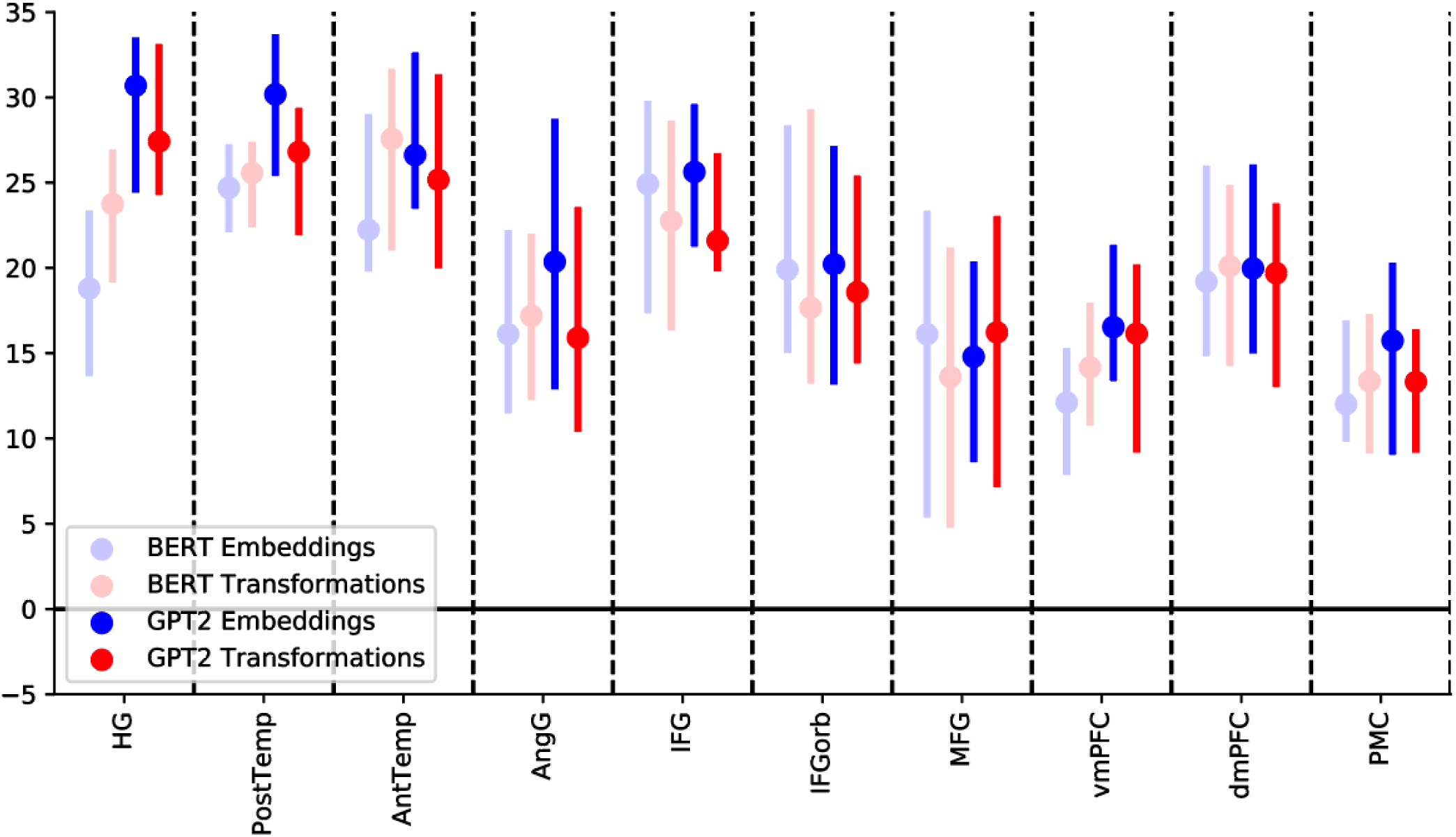
Encoding performance for GPT-2 is comparable to BERT across left-hemisphere language ROIs (cf. Fig. 2). We extracted embeddings (blue) and transformations (red) from GPT-2 and repeated the same encoding analysis as was performed using BERT. Results for BERT are reproduced from Fig. 2 for comparison. Model performance is evaluated in terms of the same noise ceiling used to evaluate BERT. Markers indicate median performance across participants and error bars indicate 95% bootstrap confidence intervals.

**Figure S4.**
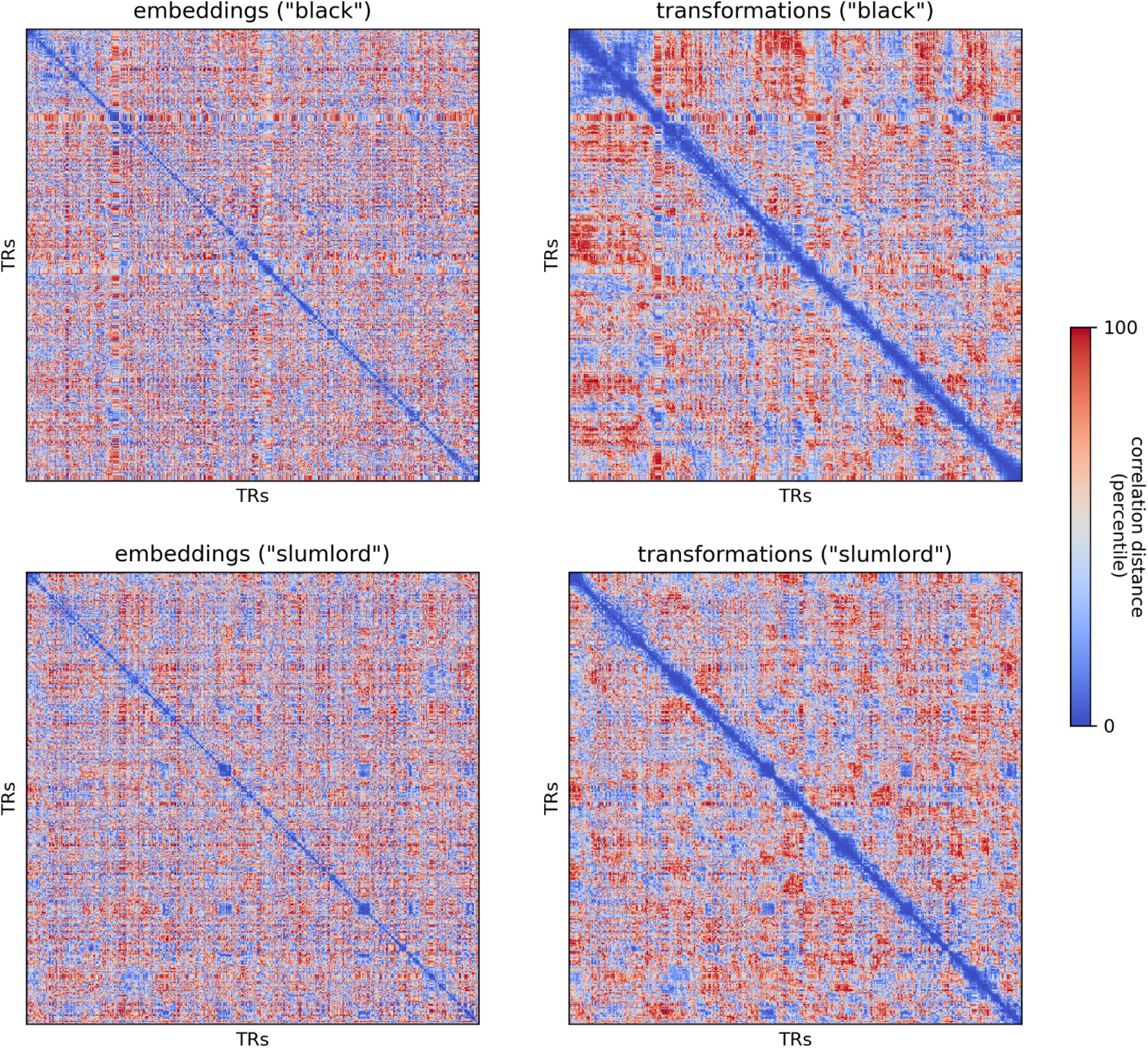
Time-point-by-time-point representational dissimilarity matrices (RDMs) for BERT embeddings (left) and transformations (right) concatenated across all layers. Despite yielding comparable encoding performance across many ROIs, the embeddings and transformations yield different representational geometries. The time-point-by-time-point RDMs are correlated at Spearman r = .768 and r = .815 for the “black” and “slumlord” stimuli, respectively. Relatively high correlations are expected when comparing representational geometries for the entire model (all 12 layers) because the transformations contextually sculpt the embeddings layer-by-layer; the layerwise representational geometries reveal a more obvious difference between embeddings and transformations (Figs. S6, S7). Dissimilarities between the vectors at each TR are measured using correlation distance (1 – Pearson correlation). Dissimilarities are colored according to percentiles.

**Figure S5.**
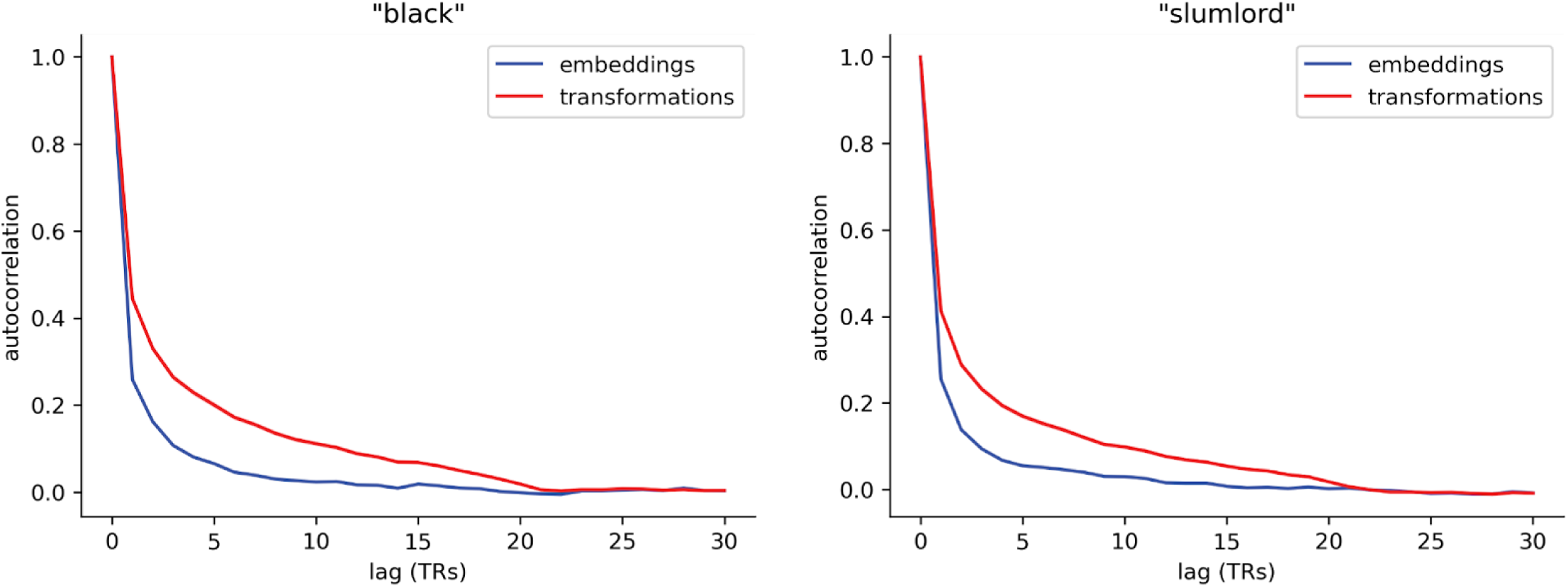
Transformations (red) have higher temporal autocorrelation than embeddings (blue) in both the “black” (left) and “slumlord” (right) story stimuli. We computed the TR-by-TR autocorrelation up to a lag of 30 TRs (45 s) separately for each feature in the embedding and transformation vectors. The blue and red lines indicate the mean temporal autocorrelation across features for embeddings and transformations, respectively. Bootstrap 95% confidence intervals are plotted around the means, but are not visible given the highly consistent temporal autocorrelation functions across features.

**Figure S6.**
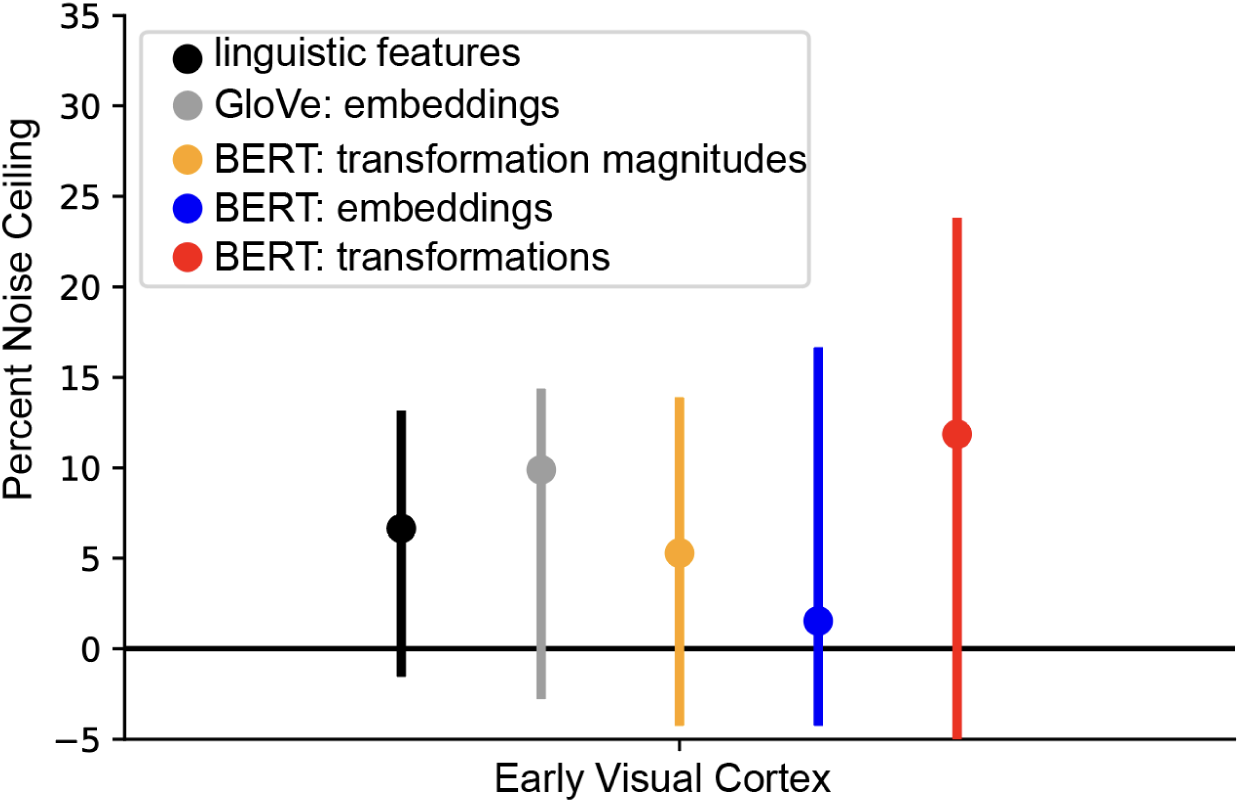
Encoding performance for three classes of language models in an early visual control ROI (cf. Fig. 2). Early visual cortex was anatomically defined as parcels overlapping with Brodmann Area 17 (primary visual cortex). As expected, none of these language features yield statistically significant prediction performance in early visual parcels. Model performance is evaluated in terms of the percent of a noise ceiling estimated using intersubject correlation. Markers indicate median performance and error bars indicate 95% bootstrap confidence intervals.

**Figure S7.**
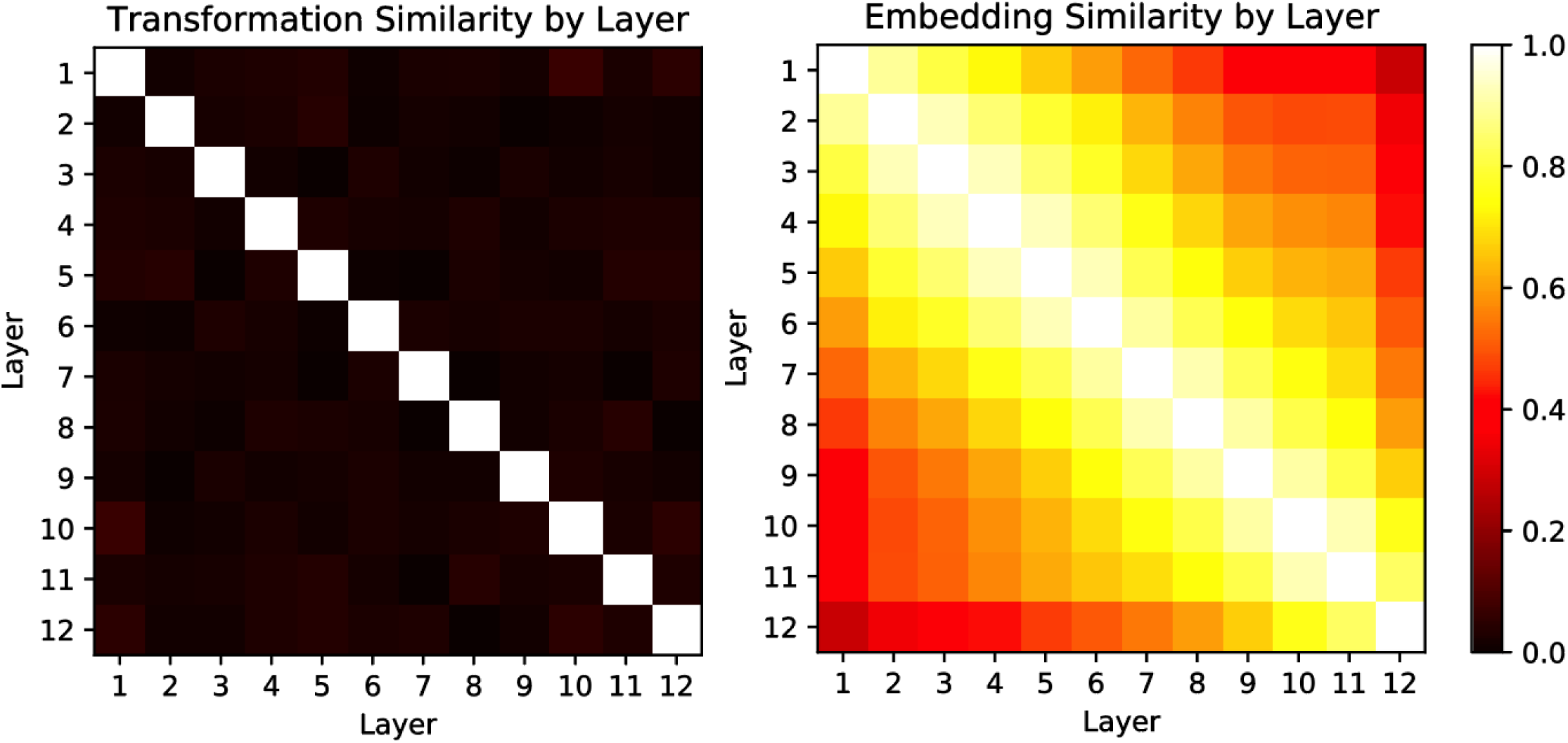
Correlation between transformations across layers (left) and embeddings across layers (right). Each word of the story stimulus was propagated through the model and the transformations and embeddings were extracted at each layer. We averaged transformations and embeddings for each word within a TR (see “Generating Transformer features” in Methods). We computed the pairwise correlation of the transformations and embeddings across the 12 layers for each TR (each TR yields a 12 × 12 correlation matrix), then averaged the resulting correlation matrices across TRs. Embeddings are much more similar across layers because embeddings accumulate contextual information layer-by-layer; on the other hand, transformations capture layer-by-layer “updates” to the embedding and are largely uncorrelated across layers.

**Figure S8.**
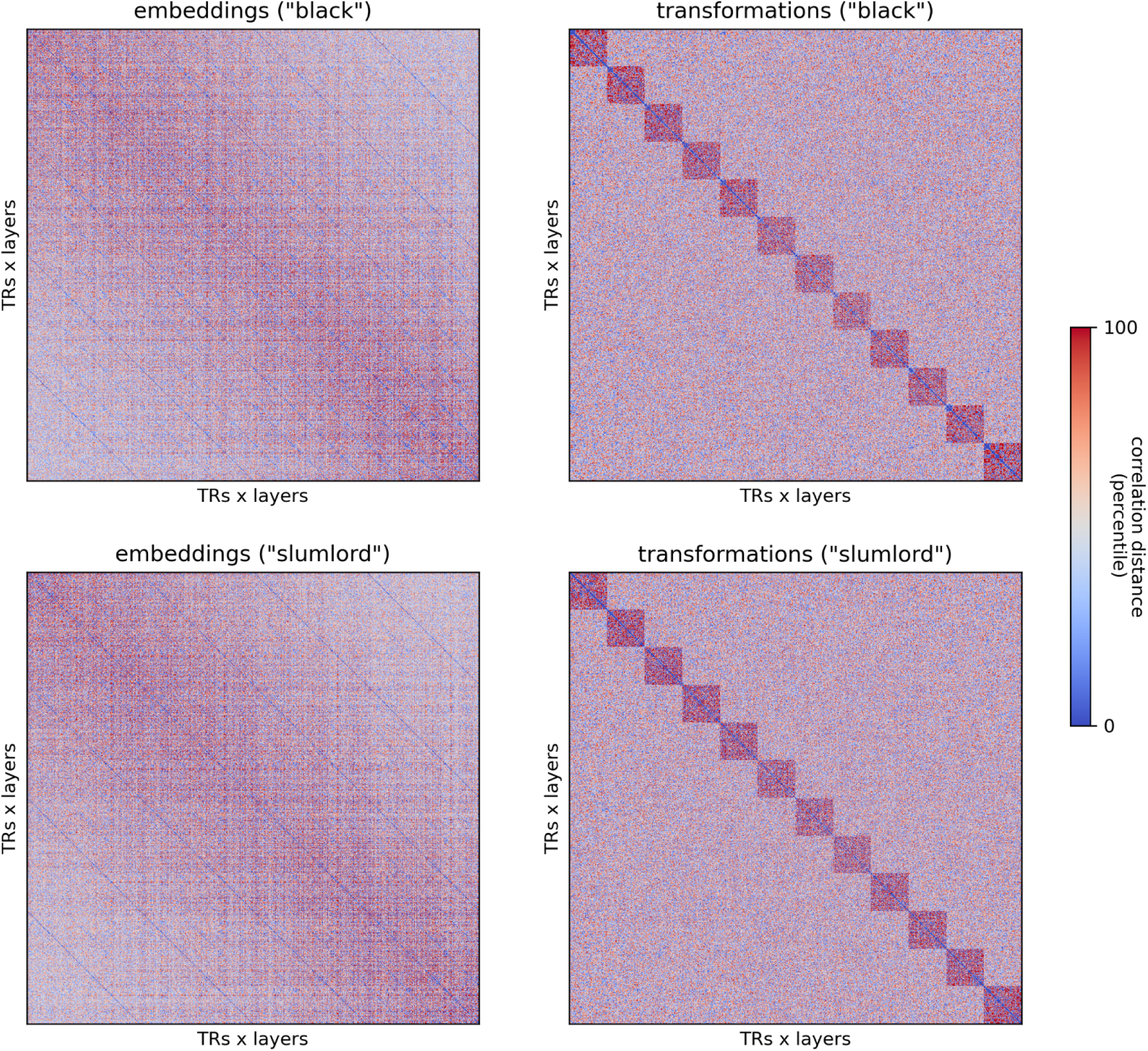
Time-point-by-time-point RDMs for embeddings (left) and transformations (right) within and across each layer of BERT. To compute the RDMs, we first split the embeddings and transformations by layer, then stacked these layers along the time axis; for example, the 12 diagonal blocks in the transformation RDMs (right) correspond to time-point-by-time-point RDMs at each layer. Cross-layer dissimilarities assume that the 768 features at each layer are shared across layers, and this is not strictly true by design. However, the embedding vectors are much more similar across layers than the transformation vectors. Overall, the RDMs for embeddings and transformations are highly dissimilar; Spearman r = .080 and .083 for the “black” and “slumlord” story stimuli, respectively. Dissimilarities between the vectors at each TR are measured using correlation distance (1 – Pearson correlation). Dissimilarities are colored according to percentiles.

**Figure S9.**
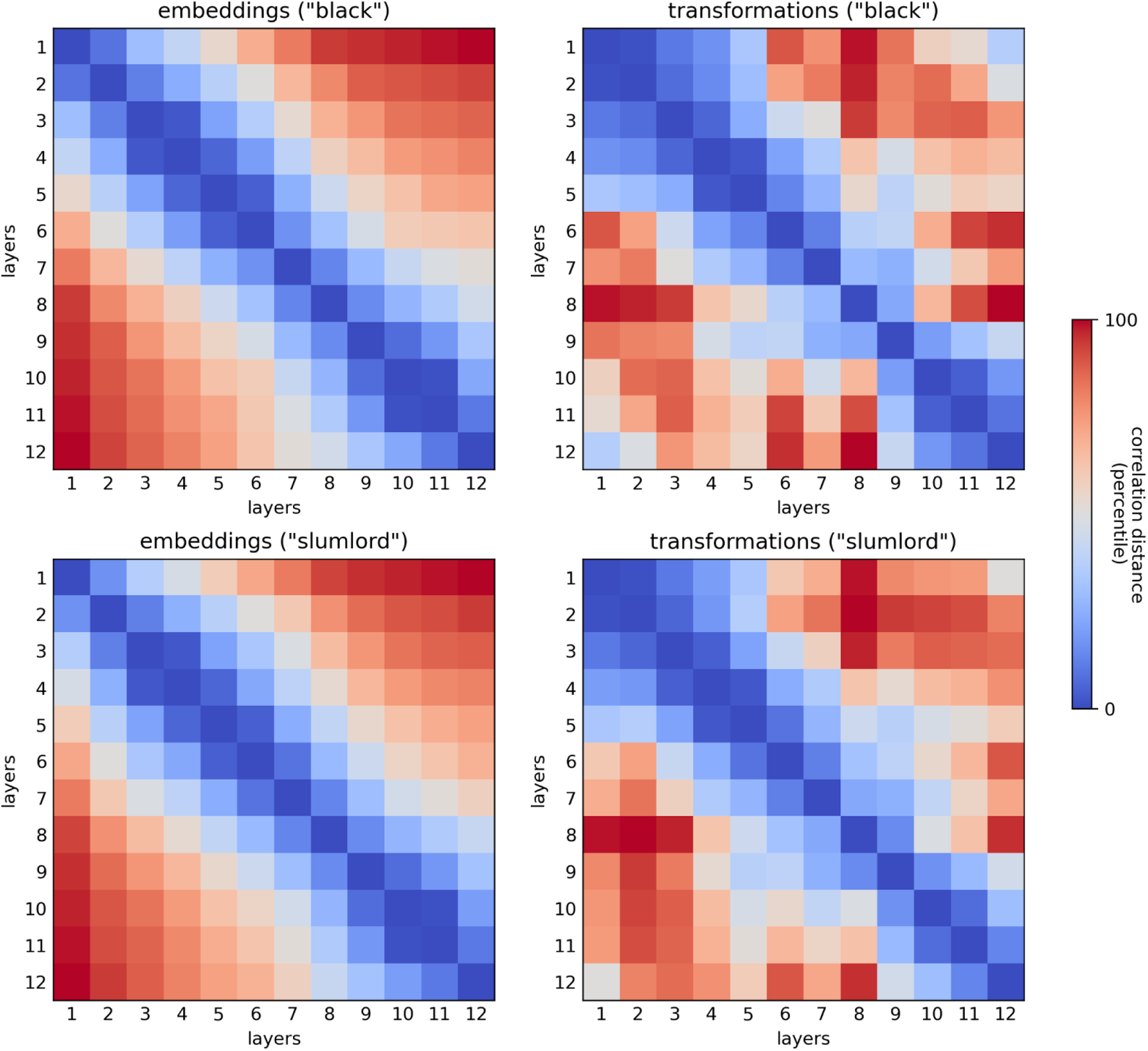
Second-order layer-by-layer representational geometry of the time-point-by-time-point RDMs for embeddings (left) and transformations (right). We first computed a time-point-by-time-point RDM for each of the 12 layers. We computed the pairwise dissimilarity of these time-point RDMs across all 12 layers, resulting in a 12-by-12 RDM. The layerwise embedding RDMs (left) demonstrate that the layerwise representational geometries evolve sequentially across layers; i.e. the representational geometry at each layer is most similar to neighboring layers and most distinct from the most distant layers. The layerwise transformation RDMs (right), on the other hand, demonstrate that the layerwise representational geometries for transformations are not as sequentially organized; for example, early-layer transformations are more similar to particular late-layer transformations than they are to particular intermediate-layer transformations. The second-order layer-by-layer RDMs are correlated at Spearman r = .750 and r = .848 for the “black” and “slumlord” stimuli, respectively. Second-order dissimilarities between the time-point-by-time-point RDMs are measured using correlation distance (1 – Pearson correlation). Dissimilarities are colored according to percentiles.

**Figure S10.**
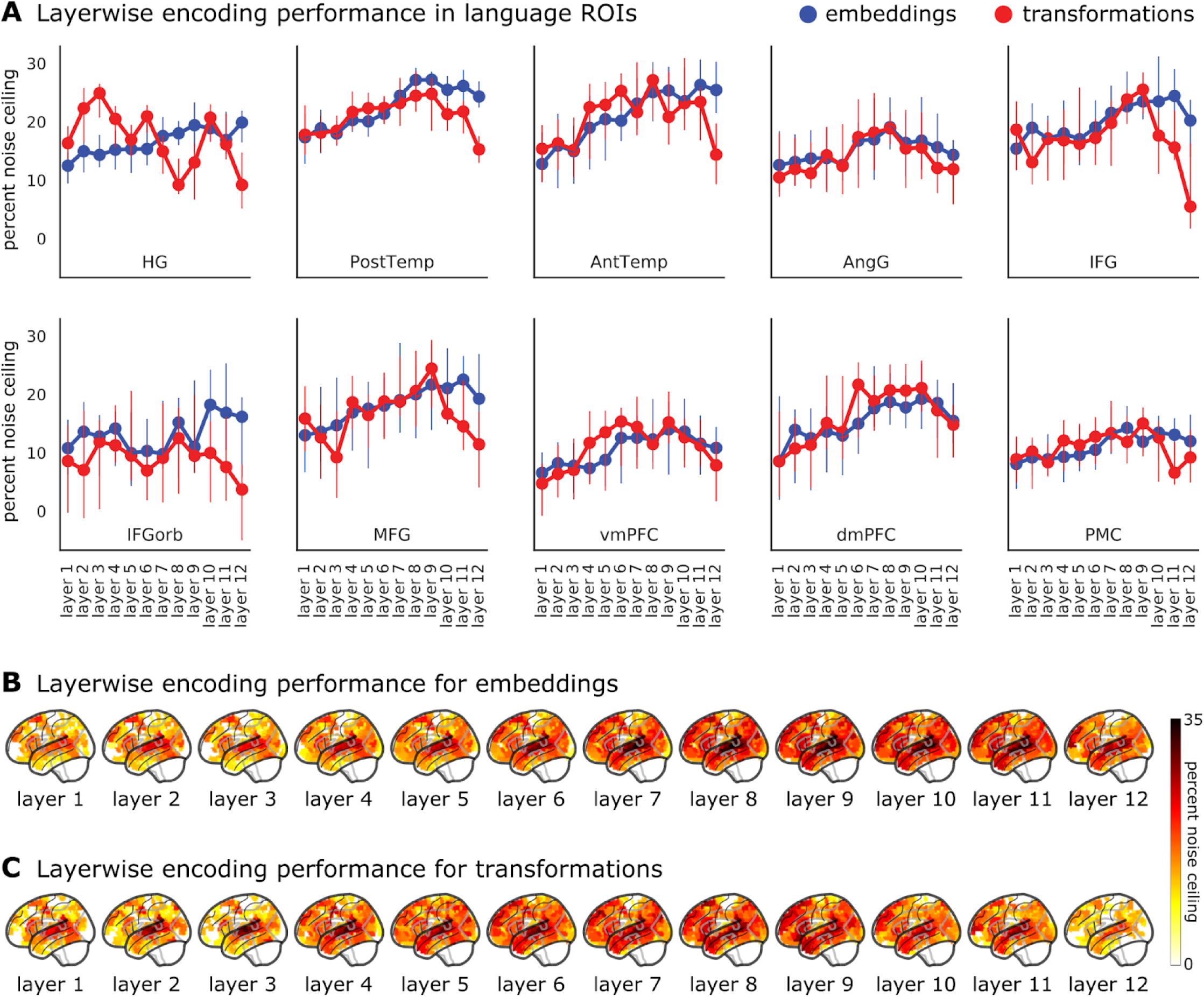
Layerwise performance of embeddings and transformations. (**A**) Layerwise model performance for embeddings (blue) and transformations (red) in ten left-hemisphere language ROIs (see Fig. S11 for right-hemisphere language areas). Model performance for embeddings tends to increase monotonically across layers, whereas performance for embeddings tends to peak for intermediate layers. Markers indicate median performance and error bars indicate 95% bootstrap confidence intervals. (**B**) Model performance for each layer of embeddings and (**C**) transformations across all cortical parcels. Cortical maps are thresholded to display only parcels with statistically significant model performance (nonparametric bootstrap hypothesis test; FDR controlled at p < .05). Model performance is evaluated in terms of the percent of a noise ceiling estimated using intersubject correlation.

**Figure S11.**
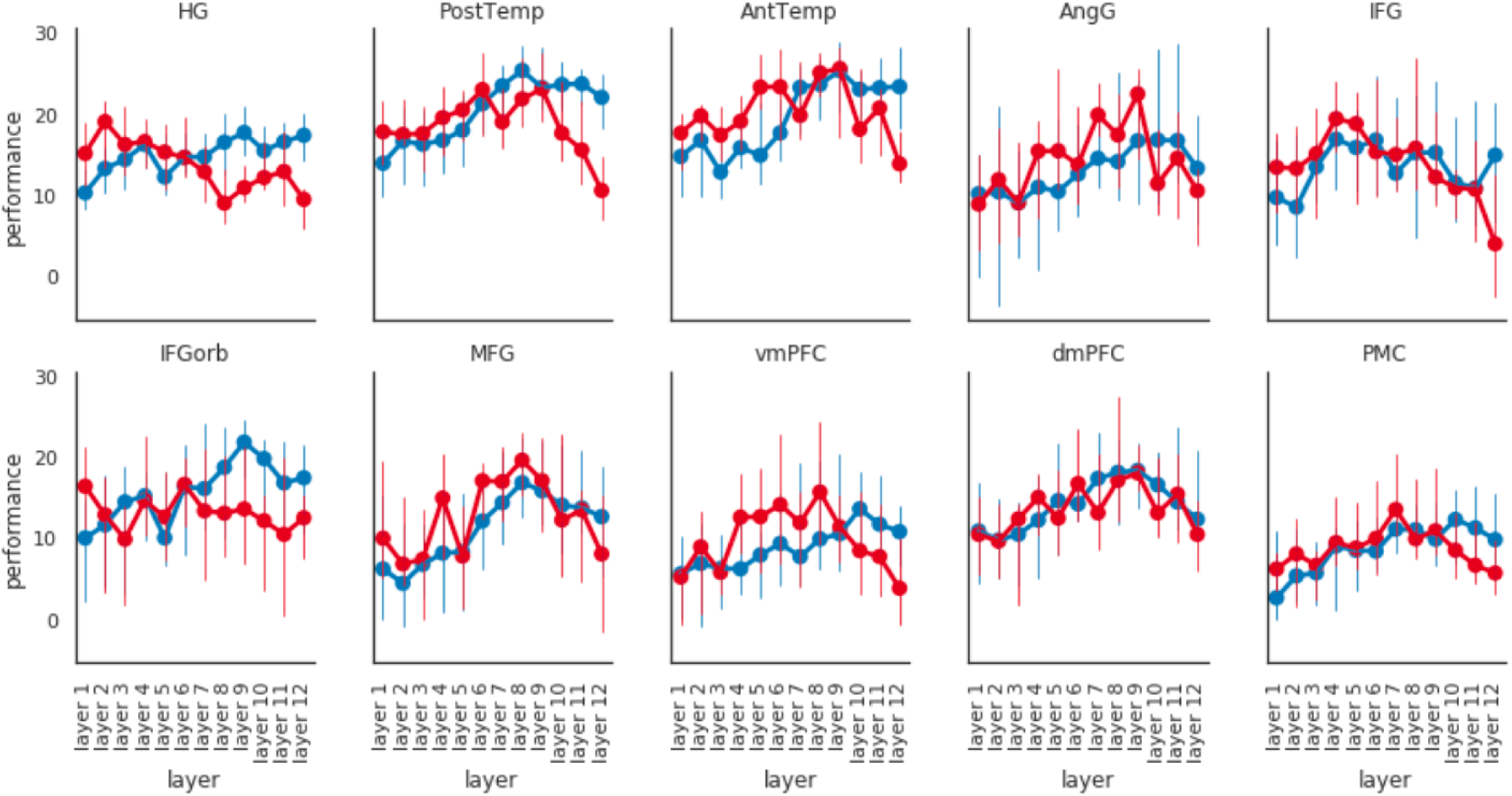
Layerwise encoding performance for embeddings (blue) and transformations (red) in right-hemisphere language areas (cf. Fig. S10A). Model performance is evaluated in terms of the percent of a noise ceiling estimated using intersubject correlation. Markers indicate median performance and error bars indicate 95% bootstrap confidence intervals.

**Figure S12.**
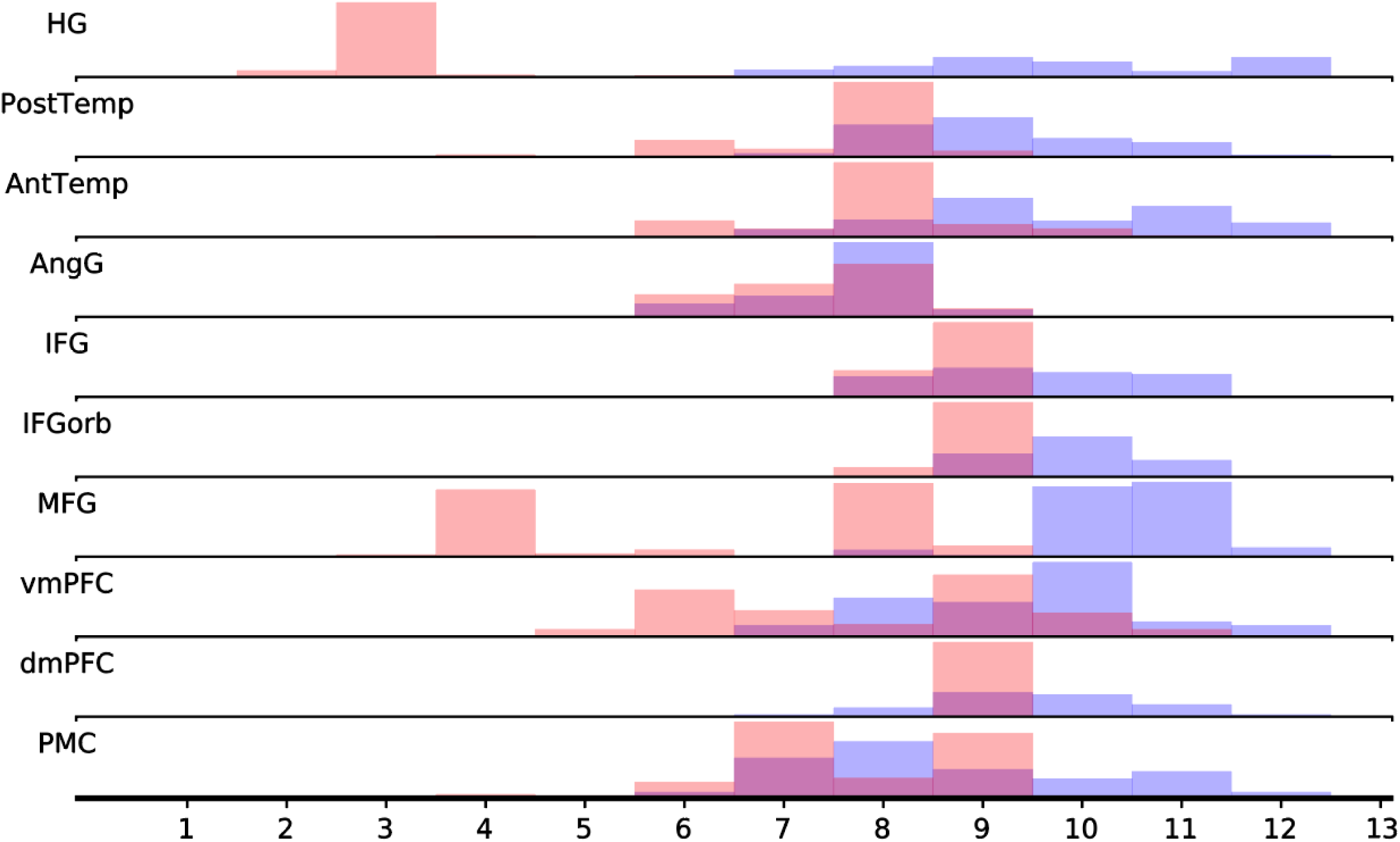
To examine the relative specificity of these layer preferences, we examined the entropy of the softmax distribution across layer performances (Tenney et al., 2019) for language ROIs, where higher entropy corresponds to a relatively uniform distribution while lower entropy corresponds to a relatively concentrated distribution. We found that performance scores for transformations were significantly more layer-specific than embeddings overall (p < .05 for all language ROIs; Table S2) where performance scores for embeddings were relatively diffuse across final layers (Fig. S9). Softmax distribution of transformation and embedding performances across different layers. The distribution of layer preferences for embeddings is more diffuse across layers than that of the transformations. To quantify this layer specificity, we quantified the entropy of these softmax distributions and found that the entropy of the embedding distribution was significantly higher than that of the transformation distribution in all of the ROIs (Table S2).

**Figure S13.**
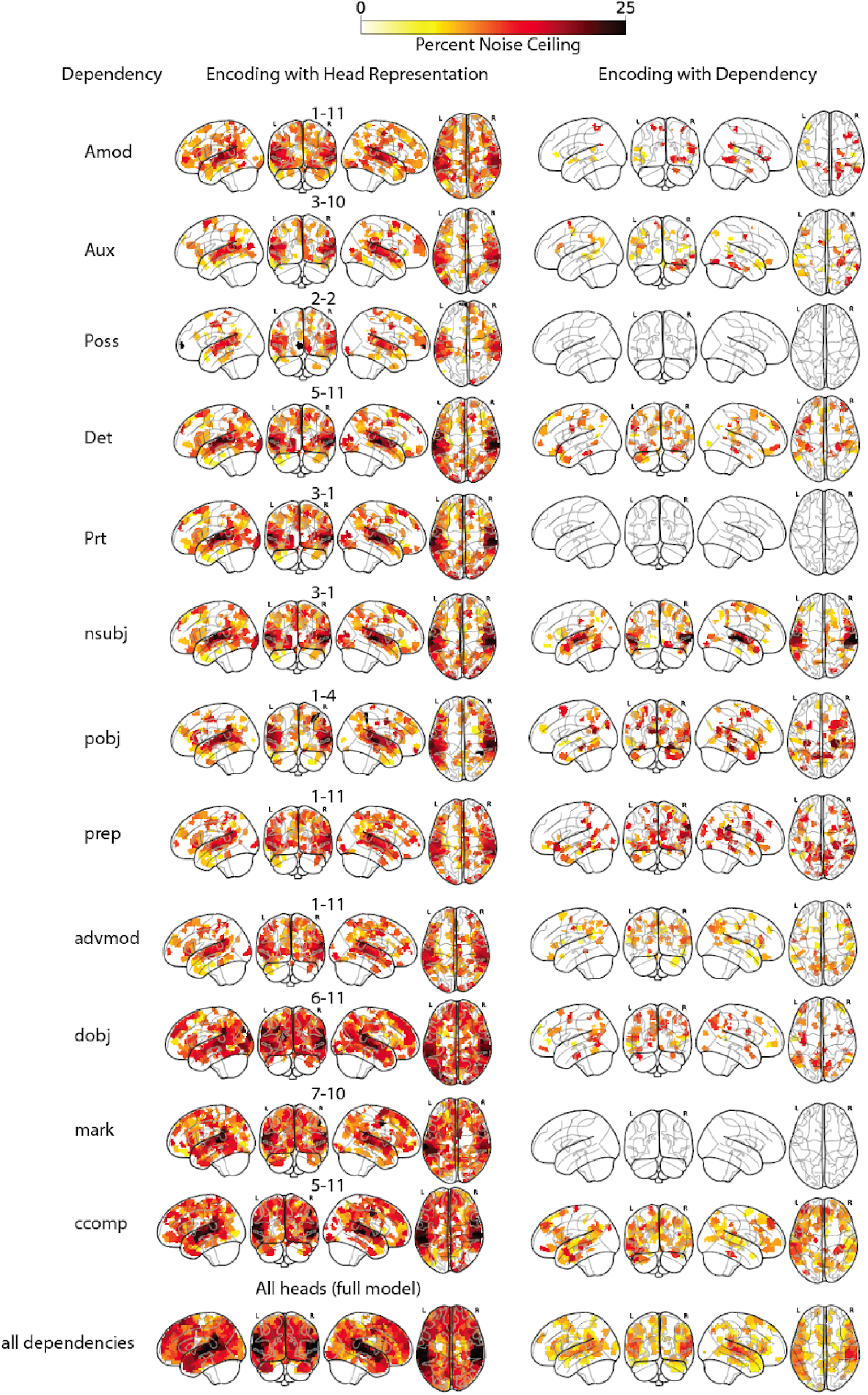
Comparison between encoding performance using the headwise transformation that best decodes a particular classical linguistic dependency (left) and the indicator variable for the linguistic dependency itself (right). The best-performing head is labeled (layer-head) for each dependency. Cortical maps are thresholded to display only parcels with statistically significant model performance (nonparametric bootstrap hypothesis test; FDR controlled at p < .05). Model performance is evaluated in terms of the percent of a noise ceiling estimated using intersubject correlation. Note that for the poss, prt, and mark dependencies, no significant parcels were found.

**Figure S14.**
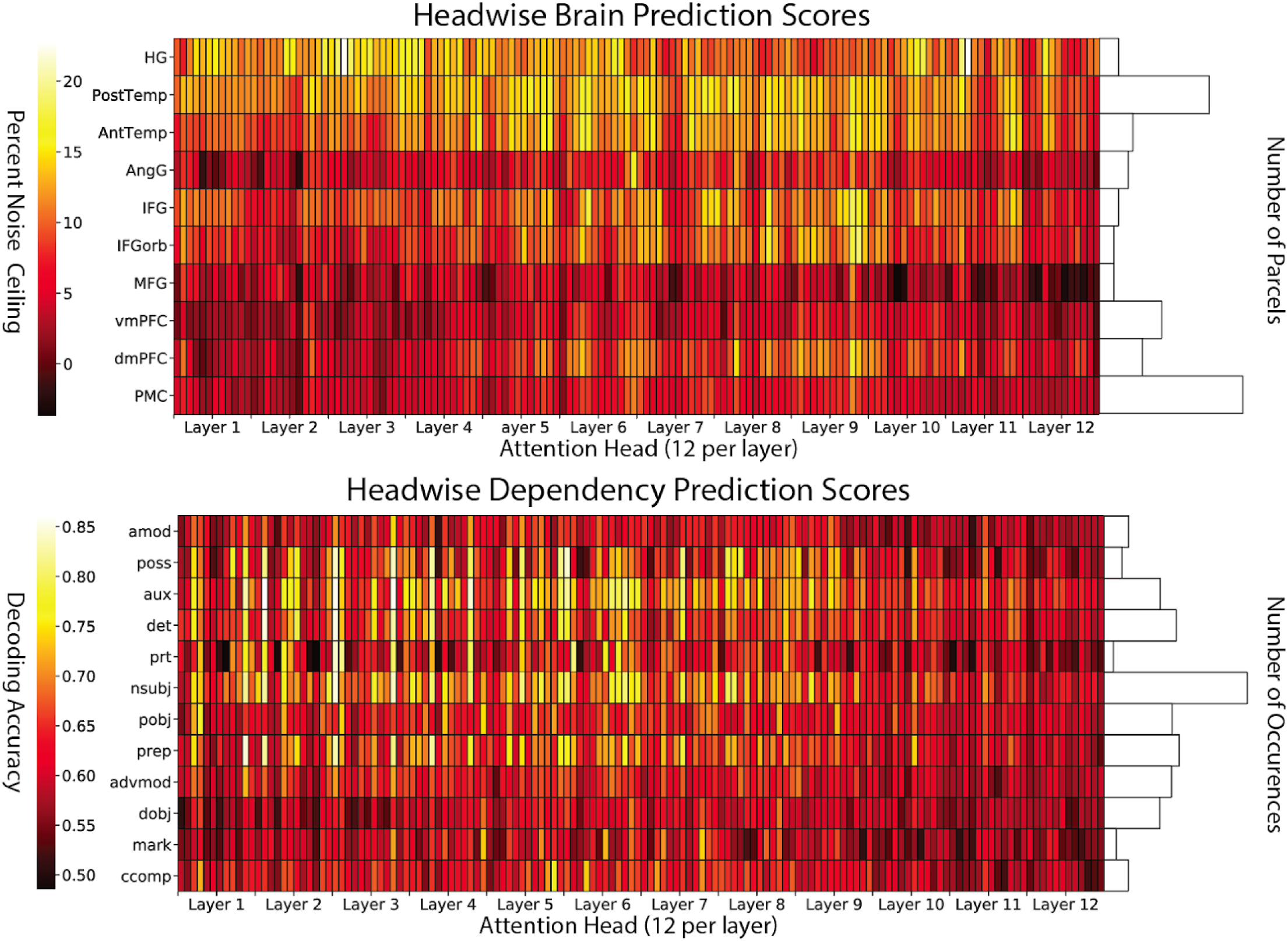
Headwise brain prediction scores (top) and headwise dependency prediction scores (bottom). At top, the headwise brain prediction scores are shown for the 12 layers of BERT (x-axis) and 10 language ROIs (y-axis). There are 12 heads per layer in BERT. The histogram at right reflects the number of parcels in each language ROI. Brain prediction scores reflect cross-validated encoding model performance evaluated in terms of the percent of a noise ceiling estimated using intersubject correlation. At bottom, the headwise dependency prediction scores are shown for the 12 layers of BERT (x-axis) and 12 classical linguistic dependencies. Dependency prediction scores reflect the classification accuracy of a cross-validated logistic regression model trained to predict the occurrence of a given linguistic dependency at each time point from the 64-dimensional transformation vector for a given attention head. The histogram at right reflects the number of occurrences of each linguistic dependency across both story stimuli (Table S5).

**Figure S15.**
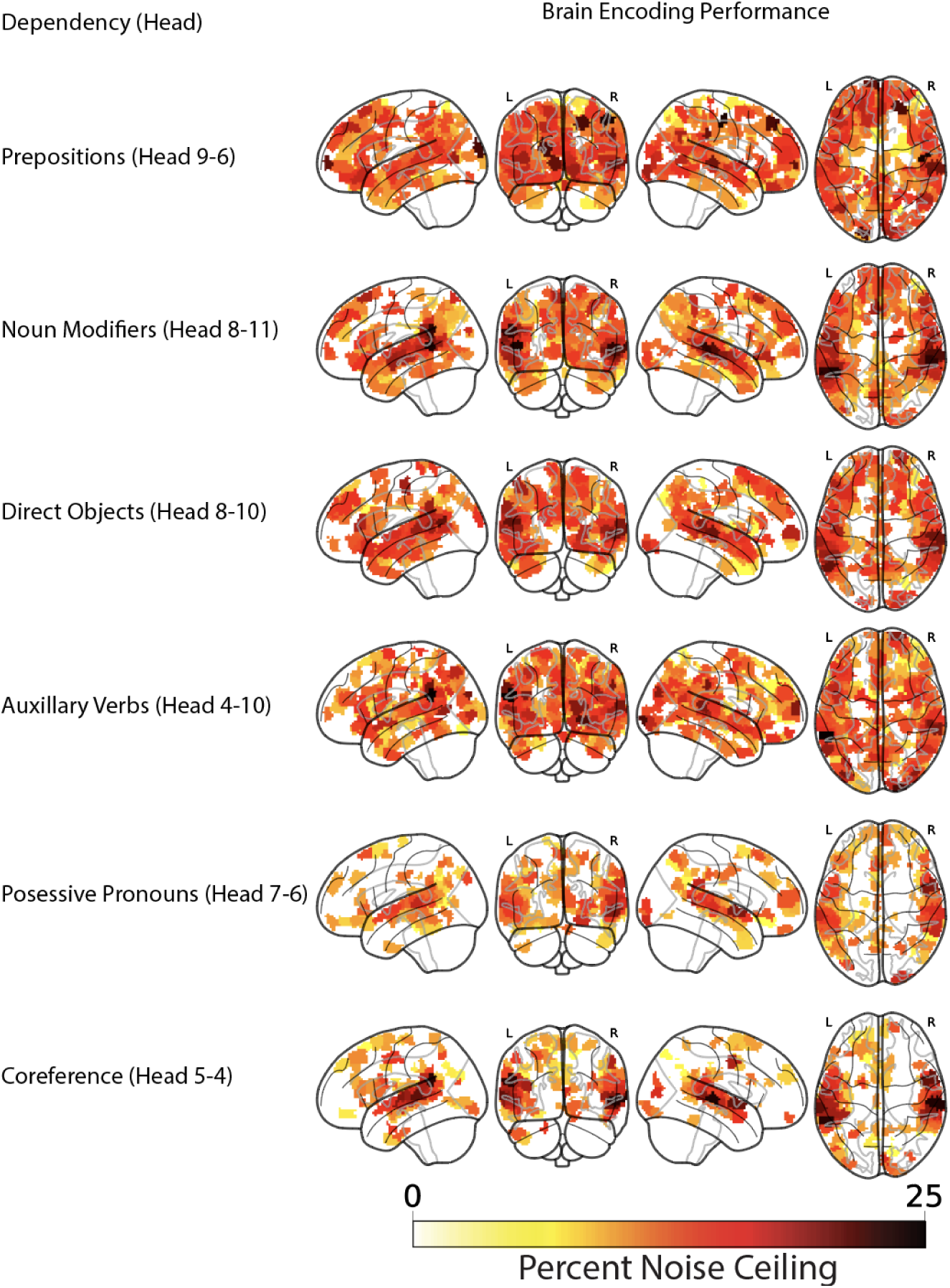
Encoding performance for heads previously highlighted in the literature for functional specialization (Clark et al., 2019). Heads are labeled according to the linguistic phenomenon for which they are specialized as well as their index in BERT (layer-head). Model performance is evaluated in terms of the percent of a noise ceiling estimated using intersubject correlation.

**Figure S16.**
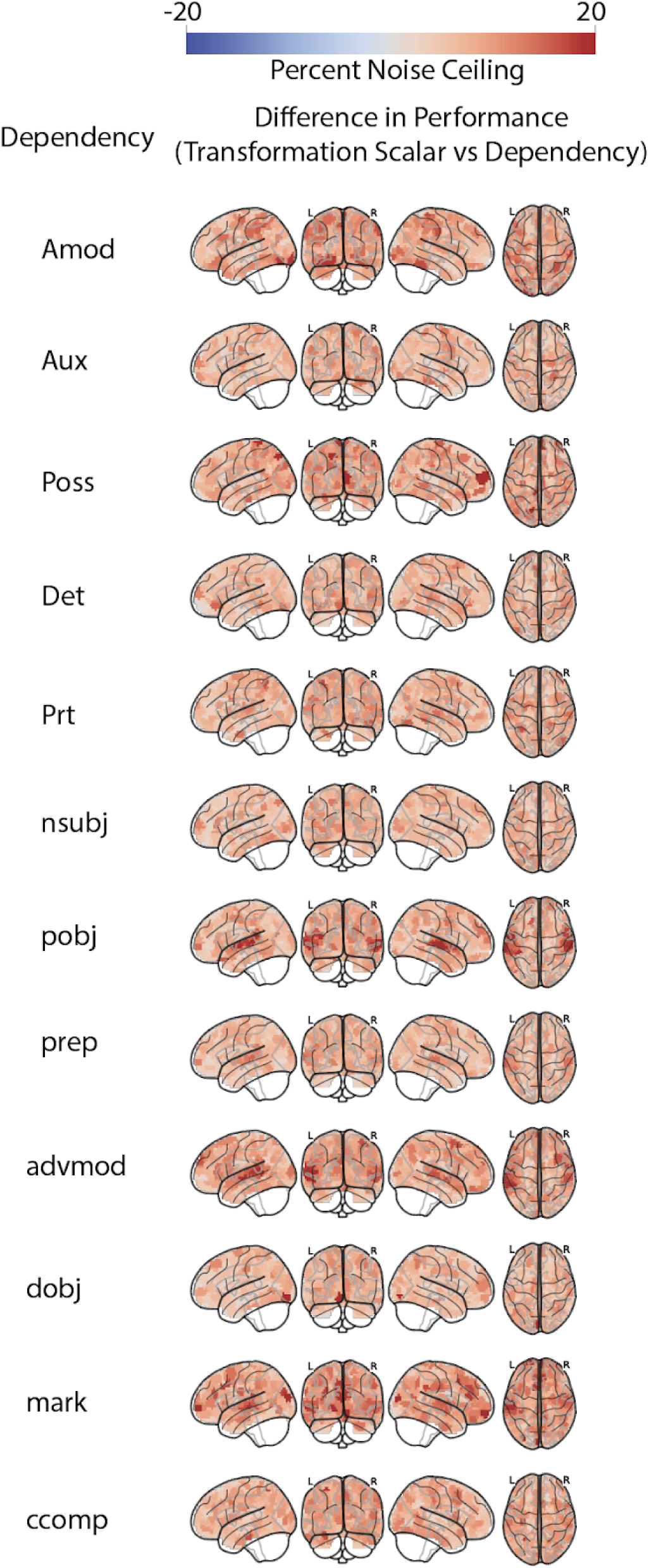
Difference in encoding performance between headwise transformations reduced to a one-dimensional scalar vector and the dependency indicator variable (unthresholded). We reduced the 64-dimensional transformation vector for the head that best decodes a particular dependency down to a single scalar by computing a weighted sum across dimensions based on the weights estimated from the logistic regression used to decode that dependency (Fig. S13). Despite the matched dimensionality, warm colors indicate that the head-specific transformation values at each TR still outperform the corresponding classical dependency indicator variable in a cross-validated encoding analysis.

**Figure S17.**
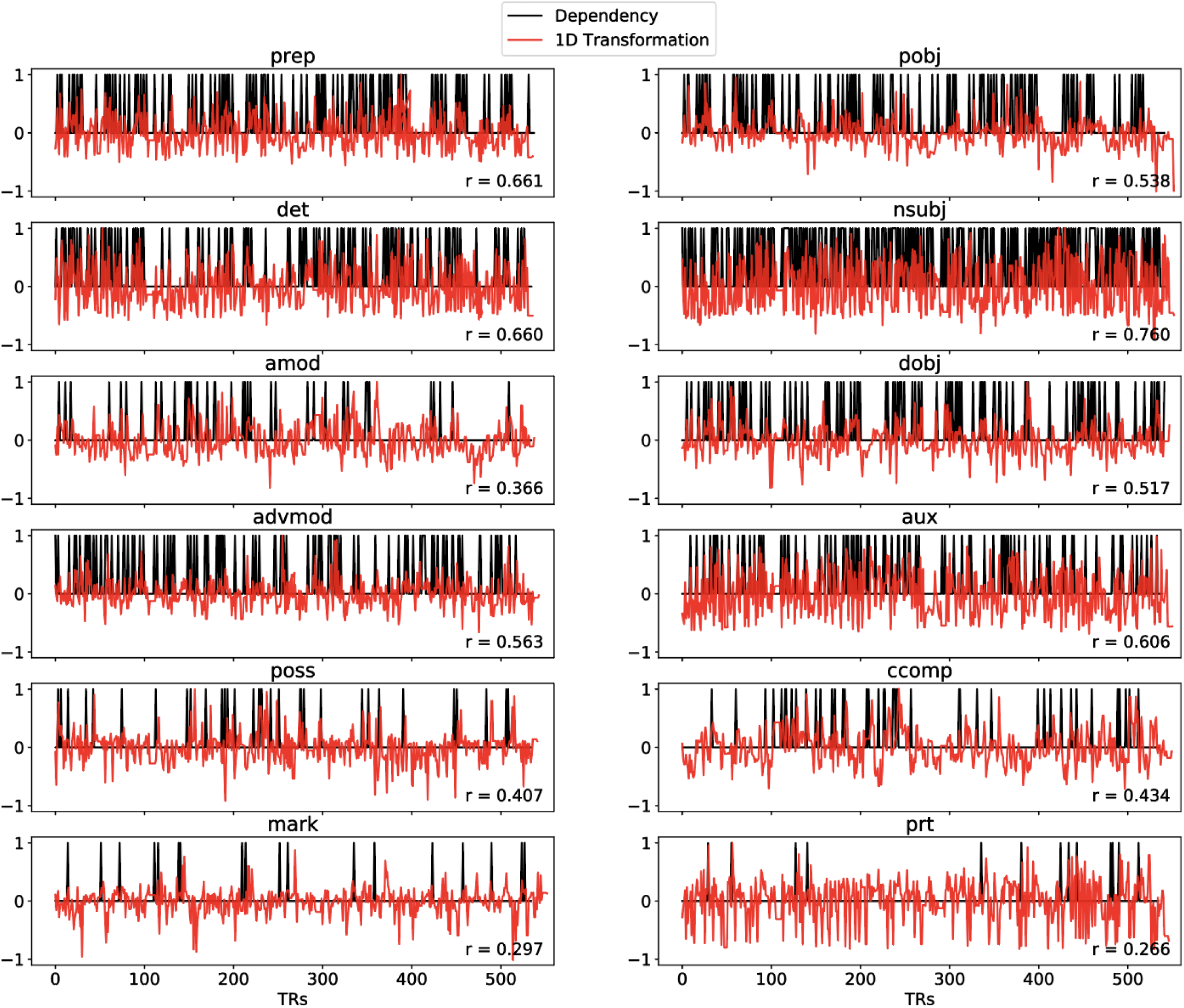
Time series of the one-dimensional transformation that best predicts a given linguistic dependency (red) and the indicator time series of the dependency itself (black) for the “I Knew You Were Black” story stimulus. We first obtained the 64-dimensional transformation vector (corresponding to an attention head) that best predicted a given classical linguistic dependency, assessed using logistic regression (see the “Decoding dependence relations” section of “Materials and methods”). To better match the dimensionality of the transformation vector and the one-dimensional linguistic dependency indicator, we computed a weighted sum of the coefficients estimated using logistic regression. This reduces the 64-dimensional transformation vector a single scalar value (Fig. S13). We then plotted how this one-dimensional transformation fluctuates over time (red) with respect to the linguistic dependency indicator (black). We mean-centered and normalized the one-dimensional transformation time series by the maximum absolute value for visualization. The one-dimensional transformation captures a graded, continuous representation of the linguistic dependency with peaks corresponding to occurrences of the linguistic dependency; correlations between the one-dimensional transformation and linguistic dependency time series are plotted at the bottom right of each panel.

**Figure S18.**
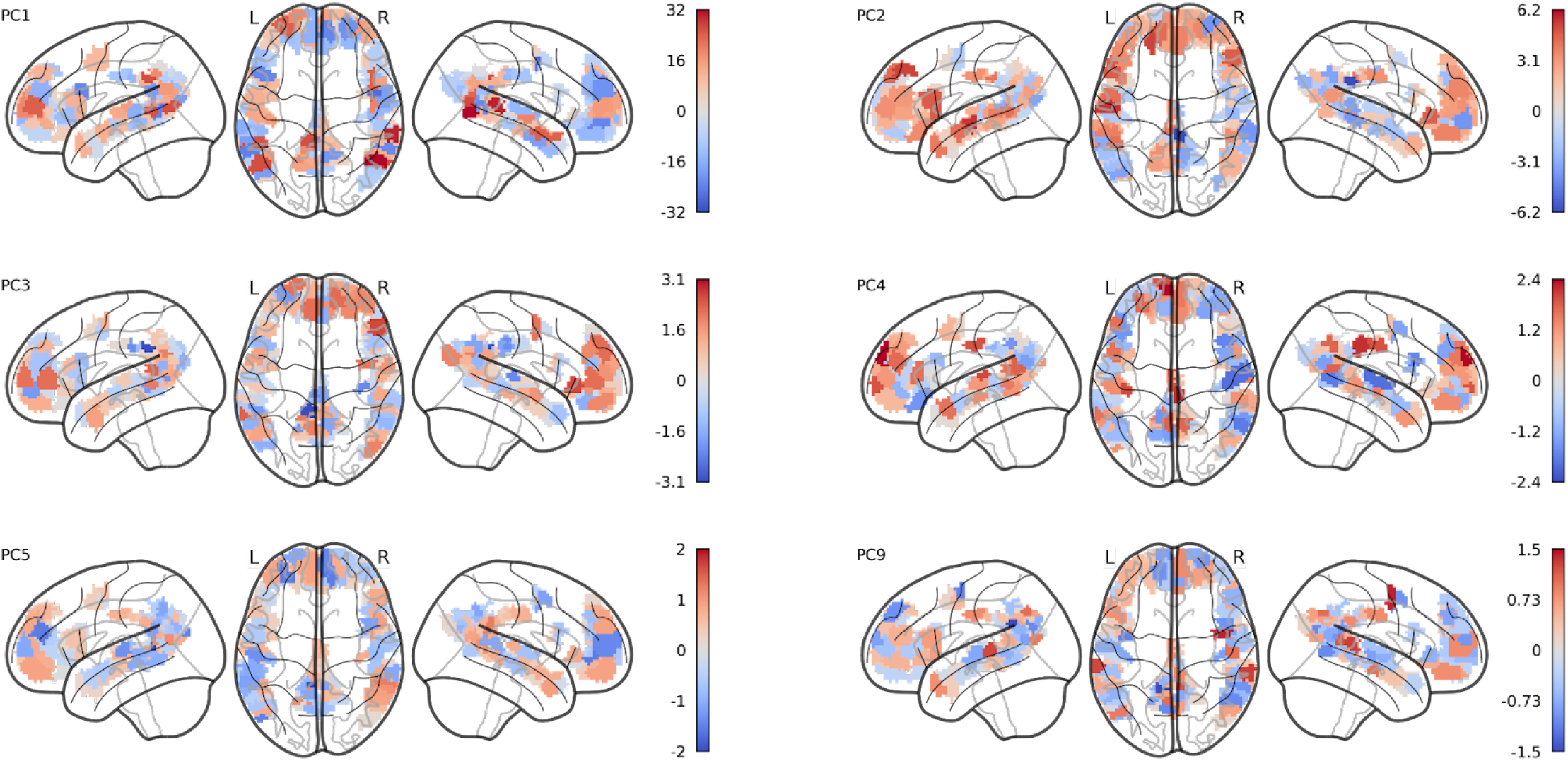
Projection of six PCs summarizing regression coefficients for the transformation-based encoding models (i.e. headwise transformation weights) onto the parcels of the language network (cf. Fig. 4). The PCs are orthogonal and capture non-redundant structure in the transformation weight matrix across the language network. Color bars reflect transformation weights for each PC (the polarity of each PC is arbitrary).

**Figure S19.**
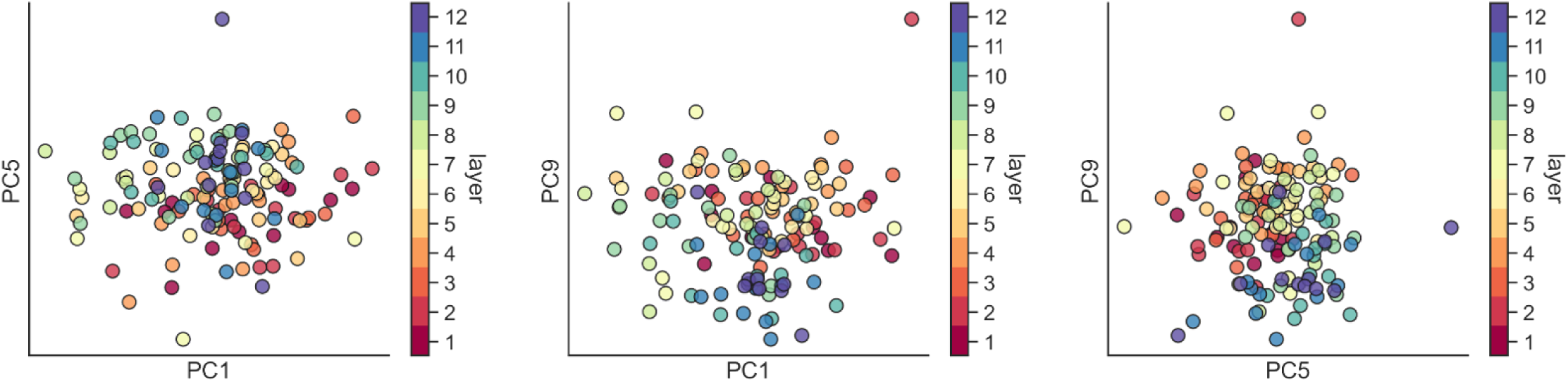
Headwise transformations weights projected into a two-dimensional space summarizing the language network colored according to the layer of each head (cf. Fig. 4D). PCs 9, 5, and 1 were the PCs most correlated with layer assignment with r = .45, .40, and .26, respectively. The PCs are orthogonal and capture non-redundant structure in the transformation weight matrix across the language network. Each PC can be projected back onto the cortical language network (Fig. S18). Each point in a scatter plot corresponds to one of the 144 attention heads.

**Figure S20.**
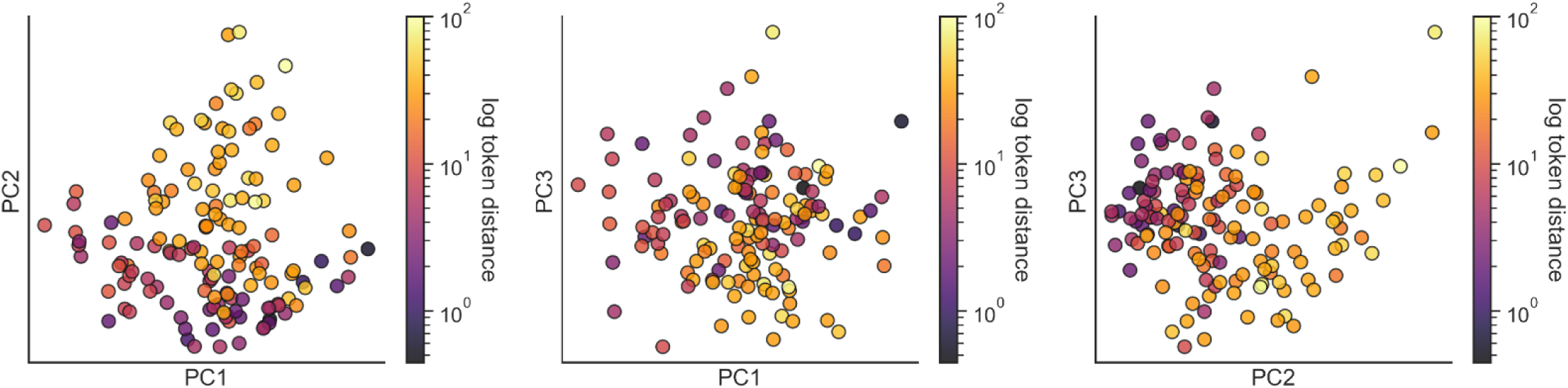
Headwise transformation weights projected into a two-dimensional space summarizing the language network colored according to the backward attention distance of each head (cf. Fig. 4E). PCs 2, 1, and 3 were the PCs most correlated with backward attention distance with r = .65, .20, .19, respectively. The PCs are orthogonal and capture non-redundant structure in the transformation weight matrix across the language network. Each PC can be projected back onto the cortical language network (Fig. S18). Each point in a scatter plot corresponds to one of the 144 attention heads.

**Figure S21.**
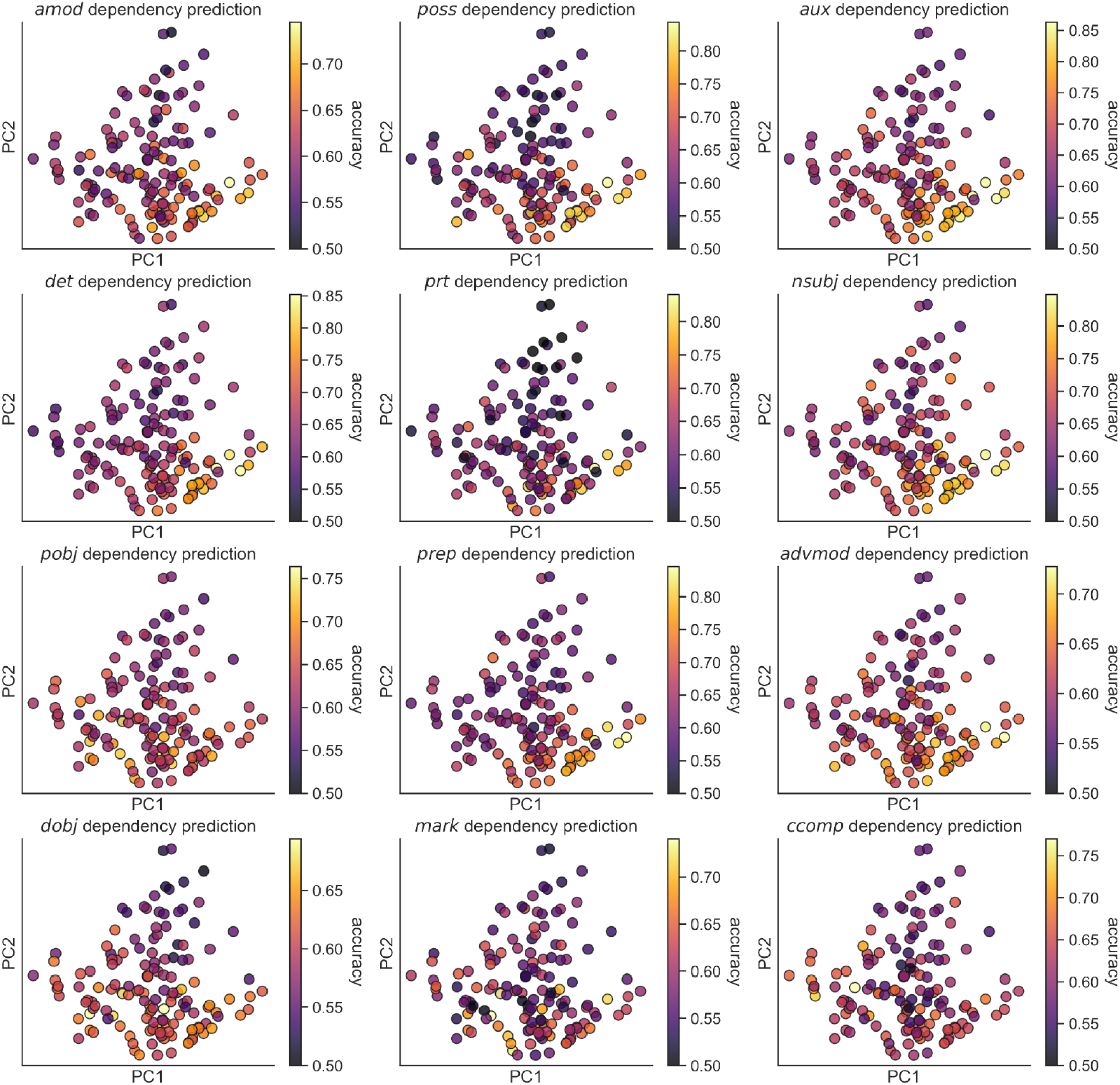
Headwise transformation weights projected into a two-dimensional space summarizing the language network colored according to each head’s dependency prediction score for each classical linguistic dependency (cf. Fig. 4). Dependency prediction scores reflect the classification accuracy of a cross-validated logistic regression model trained to predict the occurrence of a given linguistic dependency at each TR from the 64-dimensional transformation vector for a given attention head. The PCs are orthogonal and capture non-redundant structure in the transformation weight matrix across the language network. Each PC can be projected back onto the cortical language network (Fig. S18). Each point in a scatter plot corresponds to one of the 144 attention heads.

**Figure S22.**
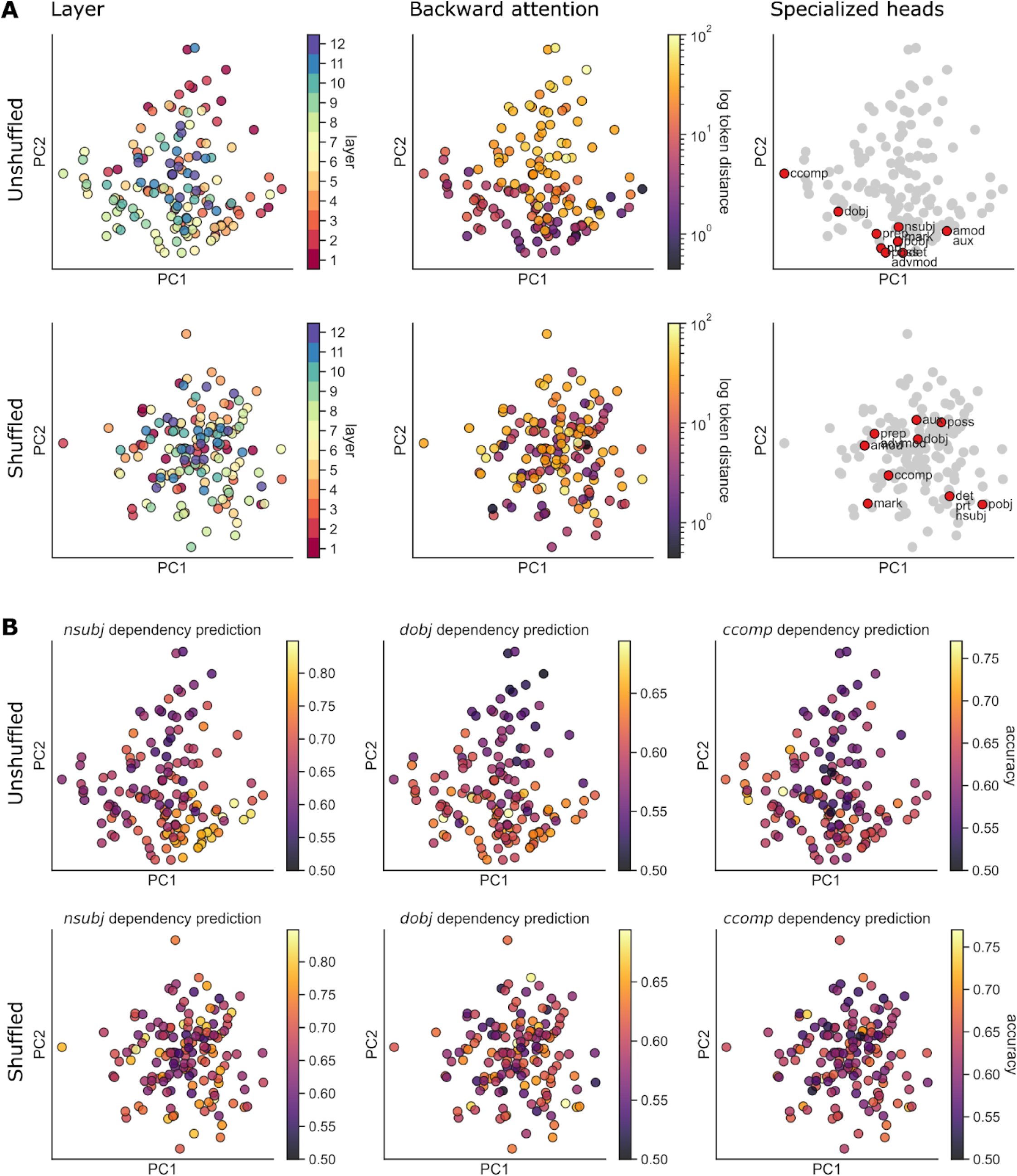
Control analysis in which we shuffle the coefficients assigned to transformation features by the encoding model across heads within each layer of BERT prior to running the analysis in Fig. 4 (i.e. immediately prior to computing the L2 norm in Fig. 4A). This perturbation effectively disrupts the functional grouping of transformation features into particular heads. (**A**) Heads are colored according to layer assignment, look-back distance (log-scale token distance), and functionally specialized heads (Clark et al., 2019) in the projection onto PCs 1 and 2. The original, unshuffled PCA solution (upper) is reproduced from Fig. 4 for comparison. In the shuffled PCA solution (lower), there is no visible structure for layer assignment or look-back distance. Disrupting the grouping of transformation features into functionally-specialized heads abolishes the low-dimensional structure of headwise contributions to predicting brain activity. (**B**) Heads are colored according to their dependency prediction scores in the projection onto PCs 1 and 2. The unshuffled PCA solution (upper) is reproduced from Fig. S21 for the nsubj, dobj, and ccomp dependencies. Dependency decoding performance falls along visible gradients in this low-dimensional brain space. In the shuffled PCA solution (lower), there are no visible gradients in dependency decoding performance.

**Figure S23.**
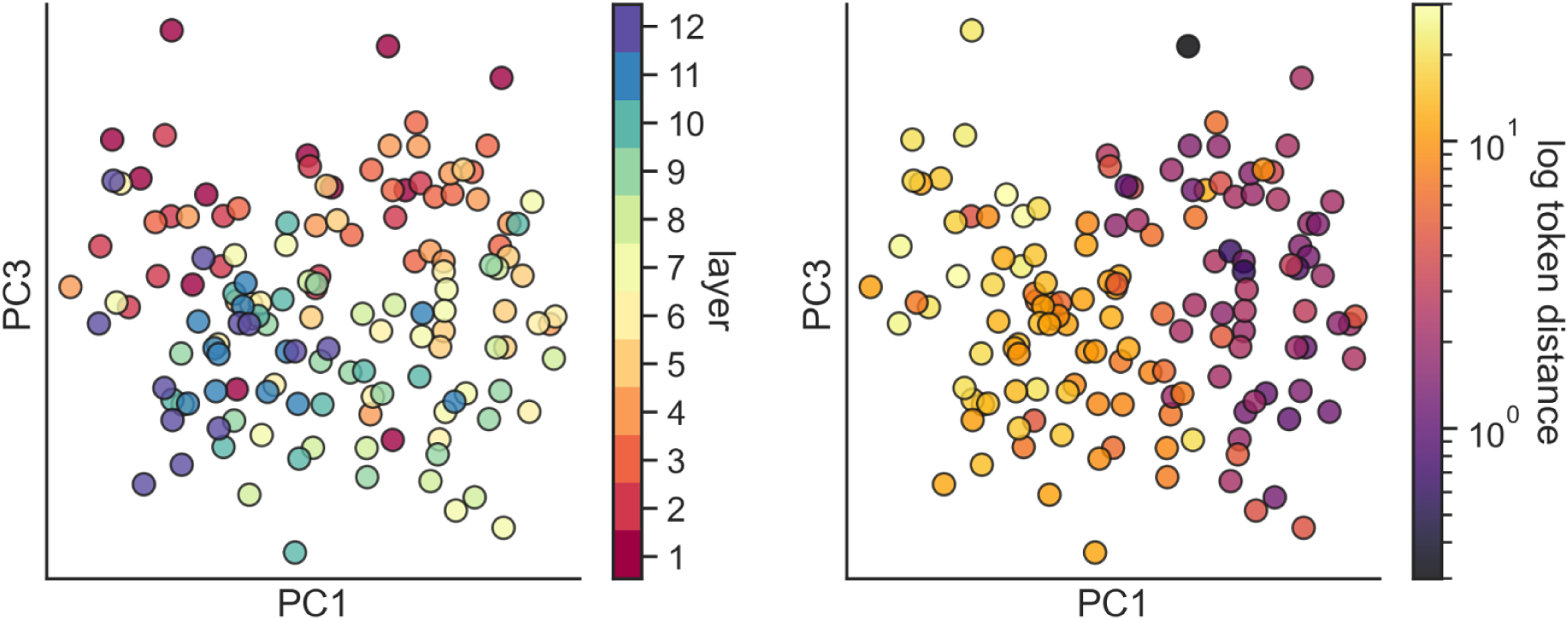
Headwise transformations in a low-dimensional brain space derived from GPT-2 (cf. Fig. 4). Following the same procedure used for BERT, we applied PCA to the weight vectors for transformation encoding models based on GPT-2 across language parcels, effectively projecting the transformation weights into a low-dimensional brain space. Each data point in the scatter plot corresponds to one of 144 heads in GPT-2. At left, heads are colored according to their layer in a reduced-dimension space of PC1 and PC3. A layer gradient is visible along PC3 (r = .60), similarly to PC1 and PC2 in BERT. At right, heads are colored according to their average backward attention distance in the story stimuli for PC1 and PC3 (look-back token distance is colored according to a log scale; color scale maximum = 50 tokens). Whereas look-back distance in BERT is most highly correlated with PC2, in GPT-2, PC1 reflects a strong gradient of look-back distance (r = .77).

**Figure S24.**
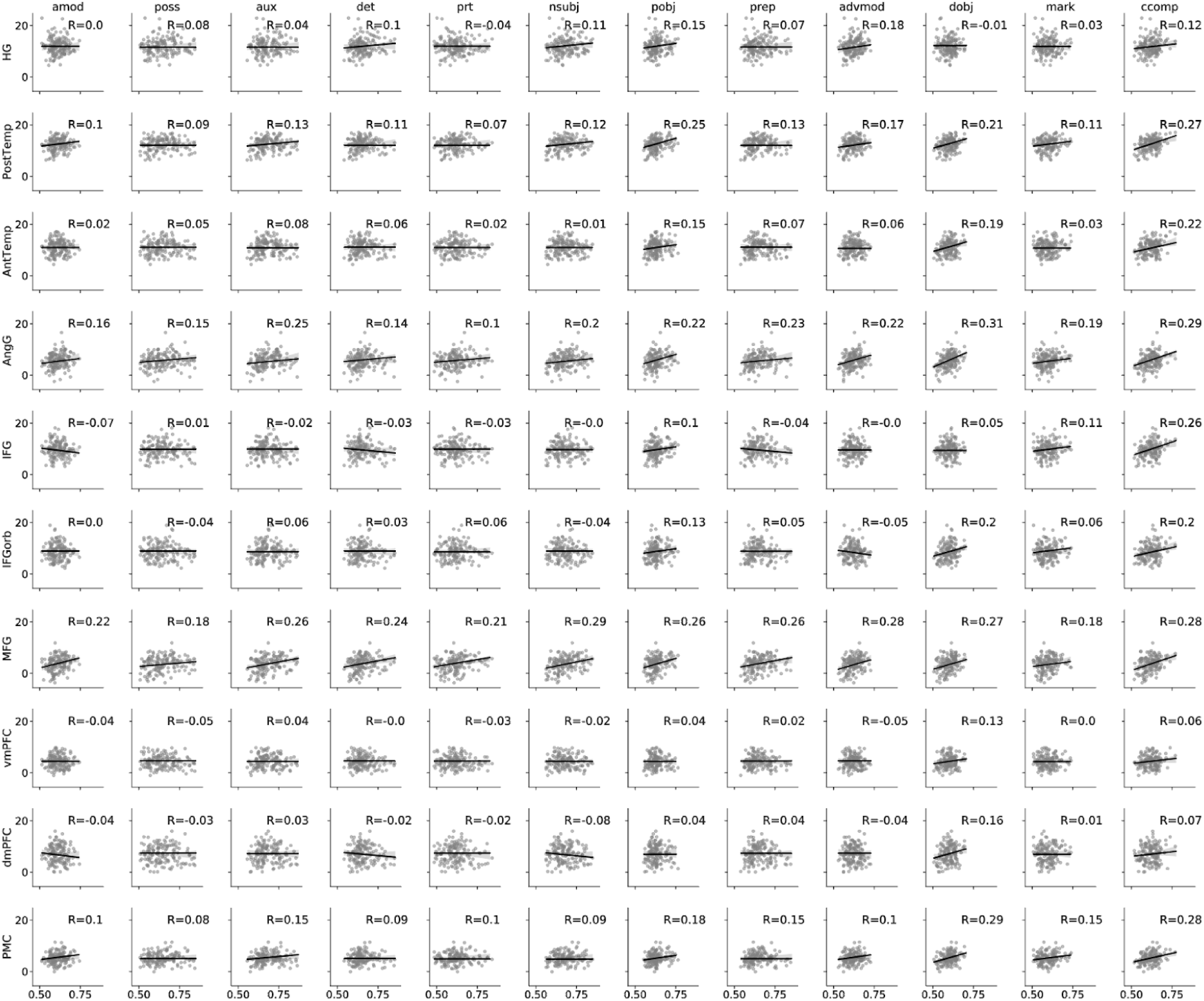
Correspondence between headwise brain prediction and dependency prediction scores for all dependencies (x-axis) and language ROIs (y-axis; cf. Fig. 5). Each point in a given scatter plot represents the dependency prediction (x-axis) and brain prediction (y-axis) scores for each of the 144 heads. Brain prediction scores reflect cross-validated encoding model performance evaluated in terms of the percent of a noise ceiling estimated using intersubject correlation. Dependency prediction scores reflect the classification accuracy of a cross-validated logistic regression model trained to predict the occurrence of a given linguistic dependency at each time point from the 64-dimensional transformation vector for a given attention head.

**Figure S25.**
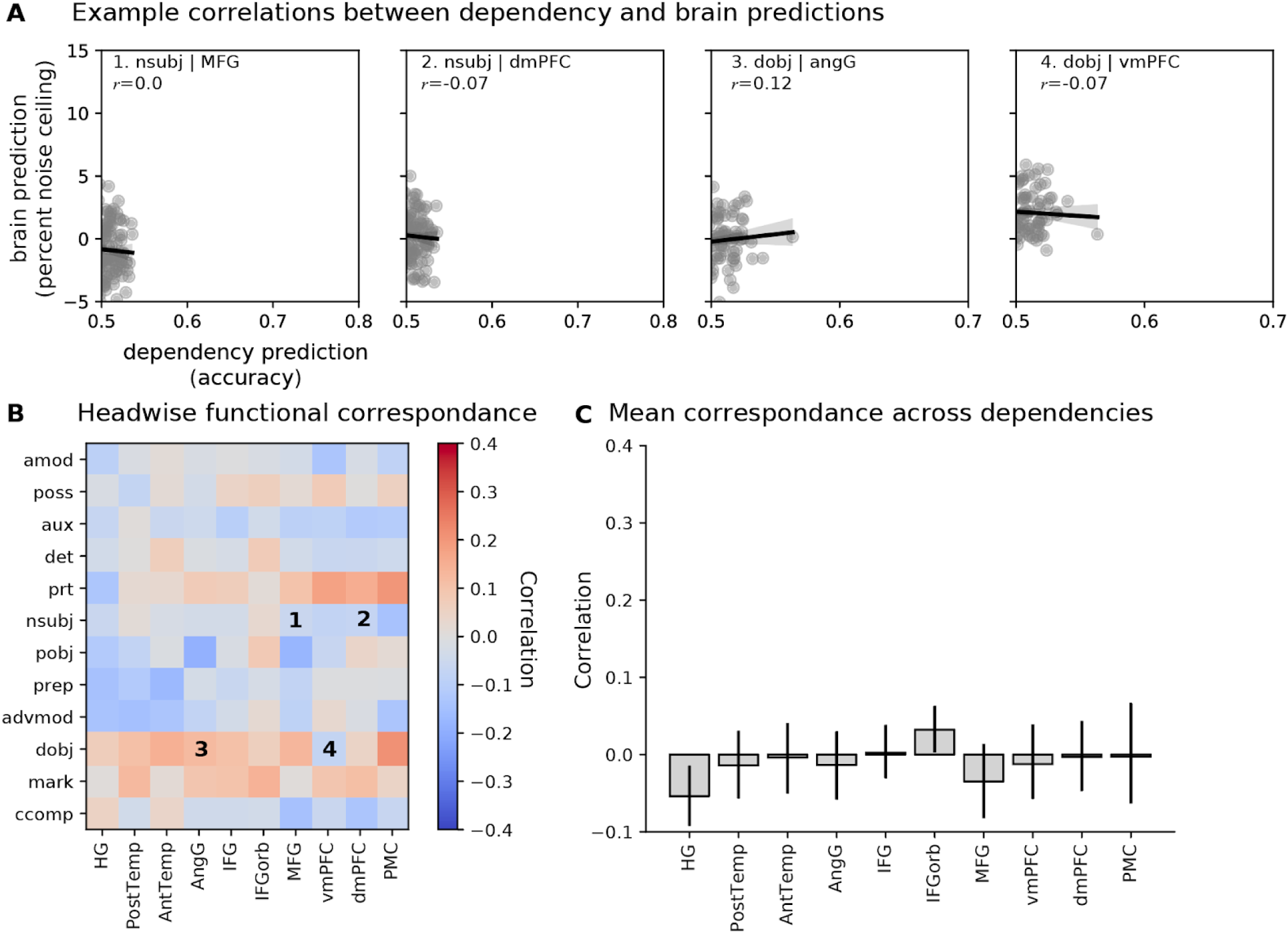
As a control analysis, we reevaluated headwise functional correspondence after shuffling the transformation features across heads within each layer (cf. Fig. 5). To do this, we first shuffled the transformation dimensions across heads within each layer, then recomputed both the brain prediction scores (the encoding model mapping transformations onto parcelwise brain activity) and the dependency prediction scores (the logistic regression model for predicting the occurrence of a given linguistic dependency) for each ROI and each linguistic dependency. We then segmented these shuffled transformations back into “pseudo-heads” and recomputed the functional correspondence of brain and dependency prediction scores across heads (cf. Fig. 5). This effectively abolishes the headwise structure of the transformations and any functional specialization therein. This yields lower brain and dependency prediction scores for the shuffled pseudo-heads (A) and reduces functional correspondence (none of the correlations were significant). (B). The 95% bootstrap confidence intervals for the mean functional correspondence across dependencies for each ROI cross zero (C), suggesting that there is no significant functional correspondence for any ROI.

**Figure S26.**
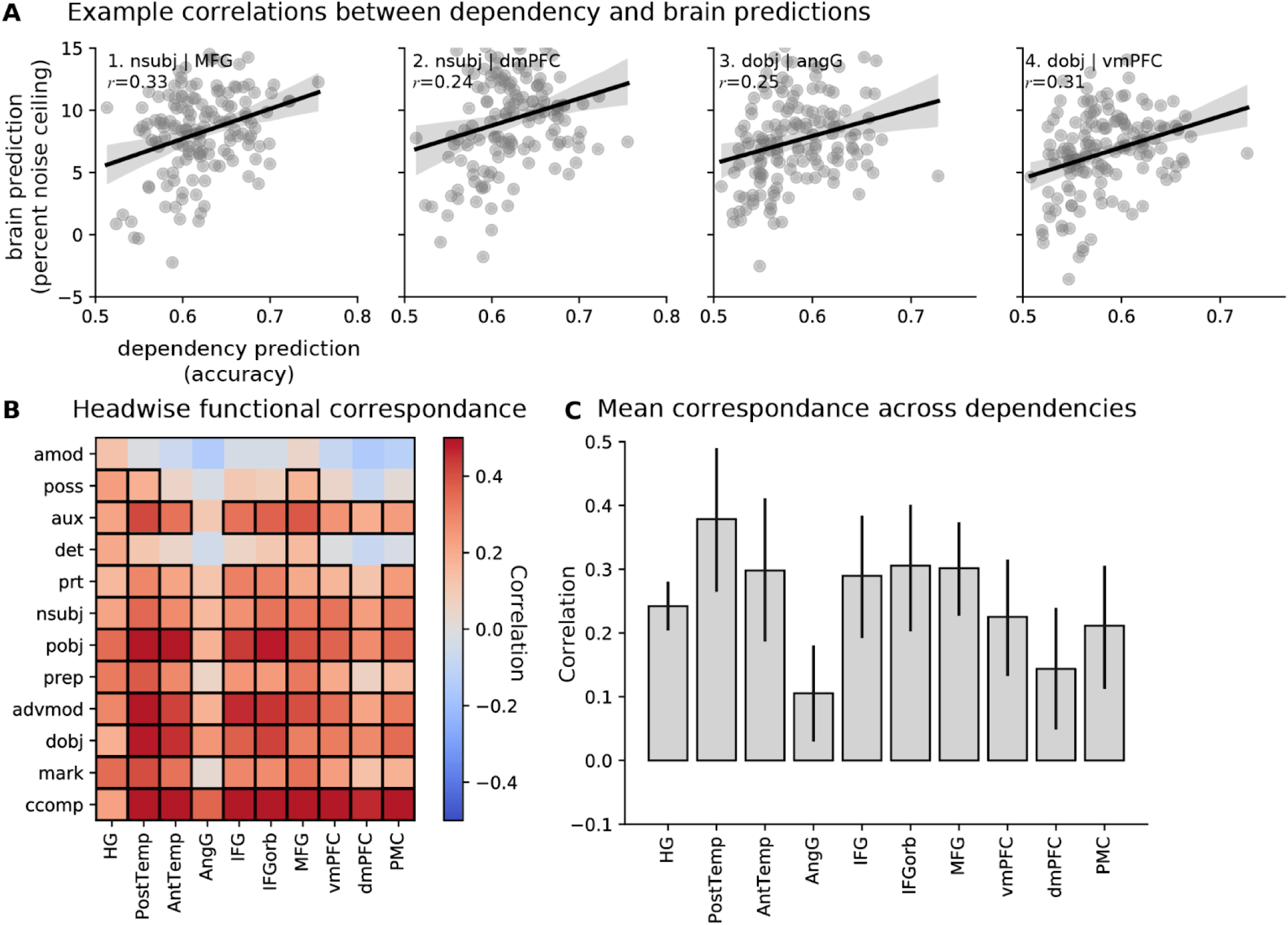
Correspondence between headwise brain and dependency predictions derived from GPT-2 (cf. Fig. 5). (**A**) Correlation between headwise brain prediction and dependency prediction scores for the same example ROIs and dependencies as Fig. 5. Each point in the scatter plot represents the dependency prediction (x-axis) and brain prediction (y-axis) scores for each of the 144 heads from GPT-2. (**B**) Correlation between headwise brain prediction and dependency prediction scores for each language ROI and syntactic dependency. Cells with black borders contain significant correlations as determined by a permutation test in which we shuffle assignments between headwise dependency prediction scores and brain prediction scores across heads (FDR controlled at p < .05). Labeled cells correspond to the example correlations in panel A. (**C**) We summarize the brain–dependency prediction correspondence for each ROI by averaging across syntactic dependencies (i.e. averaging each column of panel B). GPT-2 yields generally higher headwise correspondence than BERT (Fig. 5), but with less specificity across ROIs.

**Table S1.** Statistical significance of comparisons between BERT features (embeddings, transformations, transformation magnitudes), static GloVe embeddings, and classical linguistic features at each language ROI. For each comparison, two-tailed p-values reflect the statistical significance of the difference in encoding performance across subjects for a given ROI (permutation test; FDR controlled at p < .05).

| Comparison | HG | PostTemp | AntTemp | AngG | MFG | IFG | IFGorb | vmPFC | dmPFC | PMC |
| --- | --- | --- | --- | --- | --- | --- | --- | --- | --- | --- |
| BERT transformations vs BERT embeddings | 0.6620 | 0.0567 | 0.6700 | 0.9140 | 0.7887 | 0.7322 | 0.1517 | 1.0000 | 0.1867 | 0.0660 |
| BERT transformations vs BERT transformation magnitudes | 0.0114 | 0.0057 | 0.0025 | 0.0017 | 0.7000 | 0.0250 | 0.0025 | 0.0071 | 0.0017 | 0.0533 |
| BERT transformations vs linguistic features | 0.0017 | 0.0017 | 0.0025 | 0.0017 | 0.7000 | 0.0025 | 0.0025 | 0.0020 | 0.0017 | 0.0017 |
| BERT transformations vs GloVe embeddings | 0.0017 | 0.0017 | 0.0500 | 0.0017 | 0.7887 | 0.0025 | 1.0000 | 0.0020 | 0.0171 | 0.0017 |
| BERT embeddings vs BERT transformation magnitudes | 0.0017 | 0.0017 | 0.0025 | 0.0017 | 1.0000 | 0.0025 | 0.0025 | 0.0020 | 0.0017 | 0.0425 |
| BERT embeddings vs linguistic features | 0.0017 | 0.0017 | 0.0025 | 0.0017 | 0.7887 | 0.0025 | 0.0025 | 0.0020 | 0.0017 | 0.0017 |
| BERT embedding vs GloVe embeddings | 0.0017 | 0.0017 | 0.0140 | 0.0017 | 0.7887 | 0.0140 | 0.2256 | 0.0020 | 0.0017 | 0.0017 |
| BERT transformation magnitudes vs linguistic features | 0.0017 | 0.0017 | 0.6089 | 0.8087 | 0.7000 | 0.7322 | 0.1986 | 0.1212 | 0.7500 | 0.0017 |
| BERT transformation magnitudes vs GloVe embeddings | 0.0175 | 0.0488 | 0.0267 | 0.4814 | 0.8678 | 0.8540 | 0.2256 | 1.0000 | 0.1375 | 0.0017 |
| Linguistic features vs GloVe embeddings | 0.2633 | 0.0940 | 0.0663 | 0.8144 | 0.7000 | 0.6014 | 0.0200 | 0.0033 | 0.0017 | 0.0229 |

**Table S2.** Mean entropy of the distribution of encoding performance across layers for embeddings and transformations at each language ROIs. Statistical significance for the difference between entropy for embeddings and transformations was evaluated by bootstrap resampling the differences (FDR controlled at p < .05).

| ROI | Embedding Entropy | Transformation Entropy | Difference | p-value |
| --- | --- | --- | --- | --- |
| PostTemp | 1.2130171 | 0.68240984 | 5.126 | 0.00016667 |
| AntTemp | 0.967746731 | 0.3861663 | 6.664 | 0.00016667 |
| AngG | 0.635415239 | 0.38440445 | 2.635 | 0.0122 |
| IFG | 0.849934771 | 0.37969937 | 4.214 | 0.00342857 |
| MFG | 0.642895319 | 0.39010882 | 3.005 | 0.0122 |
| IFGorb | 0.710395586 | 0.3873914 | 3.356 | 0.0122 |
| vmPFC | 0.819666132 | 0.41612847 | 3.975 | 0.00016667 |
| dmPFC | 0.767067125 | 0.37410996 | 5.21 | 0.00016667 |
| PMC | 1.005453546 | 0.6119654 | 3.604 | 0.00016667 |
| HG | 1.067141906 | 0.51129539 | 4.432 | 0.00016667 |

**Table S3.** Descriptions and examples of the parts-of-speech. Examples are excerpted from https://universaldependencies.org/u/pos/. Phrasal verb particles, e.g. [give] in, are not included; they are tagged as ADP or ADV.

| Part-of-speech | Description | Examples |
| --- | --- | --- |
| PRON | pronoun | she, somebody, mine |
| VERB | verb | run, eating |
| NOUN | noun | girl, tree, air |
| DET | determiner | a, the, this |
| AUX | auxiliary | is, will, should |
| ADP | adposition | in, to, during |
| ADV | adverb | well, tomorrow, very |
| CCONJ | coordinating conjunction | and, or, but |
| ADJ | adjective | big, green, first |
| PART* | particle | 's, not |
| PROPN | proper noun | Mary, John, London |
| SCONJ | subordinating conjunction | if, while |
| NUM | numeral | one, two, three |
| INTJ | interjection | psst, bravo |

**Table S4.** Descriptions and examples of the dependency relations in Fig. 5. The labels are part of ClearNLP dependency labels (https://github.com/clir/clearnlp-guidelines/blob/master/md/specifications/dependency_labels.md). Examples are adapted from https://universaldependencies.org/u/dep/all.html. Note that poss, prt, and dobj in the table correspond to nmod:poss, compound:prt, and obj in the universal dependencies, respectively; pobj and prep are not part of the universal dependencies.

| Dependency | Description | Example |
| --- | --- | --- |
| amod | adjectival modifier | Sam eats [large]<sub>amod</sub> hot [dogs]<sub>head</sub>. |
| poss | possessive nominal modifier | [Marie]<sub>poss</sub> 's [book]<sub>head</sub> |
| aux | auxiliary | He [should]<sub>aux</sub> [leave]<sub>head</sub>. |
| det | determiner | [Which]<sub>det</sub> [book]<sub>head</sub> do you prefer? |
| prt | phrasal verb particle | They [shut]<sub>head</sub> [down]<sub>prt</sub> the station. |
| nsubj | nominal subject | The [car]<sub>nsubj</sub> is [red]<sub>head</sub>. |
| pobj* | object of preposition | We went [to]<sub>head</sub> the grocery [store]<sub>pobj</sub>. |
| prep* | preposition | We [went]<sub>head</sub> [to]<sub>prep</sub> the grocery store. |
| advmod | adverbial modifier | [Genetically]<sub>advmod</sub> [modified]<sub>head</sub> food |
| dobj | direct object | She [gave]<sub>head</sub> me a [raise]<sub>dobj</sub>. |
| mark | subordinate clause marker | He says [that]<sub>mark</sub> you [like]<sub>head</sub> to swim. |
| ccomp | clausal complement | He [says]<sub>head</sub> that you [like]<sub>ccomp</sub> to swim. |

**Table S5.** Number of TRs in which linguistic dependency occurs in the two story stimuli. The column headers indicate the total number of TRs for that stimulus.

| Dependency | “black” (~/534) | “slumlord” (~/619) |
| --- | --- | --- |
| prep | 121 | 214 |
| pobj | 109 | 195 |
| det | 117 | 206 |
| nsubj | 245 | 392 |
| amod | 43 | 66 |
| dobj | 95 | 153 |
| advmod | 122 | 177 |
| aux | 99 | 150 |
| poss | 35 | 45 |
| ccomp | 46 | 61 |
| mark | 19 | 35 |
| prt | 13 | 29 |

## Footnotes

1 There is an ongoing debate as to whether autoregressive transformers (e.g., GPT; Radford et al., 2018, 2019; Brown et al., 2020) or bi-directional transformers (e.g., BERT; Devlin et al., 2019) are more appropriate models for predicting brain activity (Antonello and Huth, 2022). In this work, we chose BERT as a more plausible model for narrative comprehension. This is because BERT’s “bidirectional” attention allows later words in a sentence to affect the meaning of earlier words, whereas GPT’s “causal” attention does not. To understand the implications, consider the example shown in Fig. 1 above, containing the words “the secret plan.” GPT is autoregressive, using a “causal” (rather than “bidirectional”) attention mechanism. This means that information can only flow forwards in time: the representation of “plan” can be updated based on “secret”, but the representation of “secret” cannot be retroactively updated based on “plan.” In contrast, BERT allows bidirectional attention within the context window, so the two words can affect each others’ representations. Humans clearly can and do operate bidirectionally: we are able to update our representation of earlier words based on later ones (e.g., cataphora; Carden, 1982). Given our focus on the contextualization process itself (i.e. transformation vectors), we chose BERT as the more realistic model; however, we replicate our key findings in GPT-2 to showcase the generalizability of our approach.

2 We omit the original static BERT embeddings (which are sometimes termed “Layer 0”) and compare BERT layers 1-12 to the 12 transformation layers.

3 We chose 20 TRs as it represents 30 seconds of audio data, averaging around 100 Transformer tokens. The majority of BERT’s training occurred on sequences of 128 tokens, so this ensured that BERT was exposed to sequence lengths that were similar to its training distribution.

4 Albeit overspecialized for a particular downstream training objective; i.e. the cloze task for BERT.

5 We excluded the special [SEP] token.

## References

Aly, M., Chen, J., Turk-Browne, N. B., & Hasson, U. (2018). Learning naturalistic temporal structure in the posterior medial network. Journal of Cognitive Neuroscience, 30(9), 1345–1365. https://doi.org/10.1162/jocn_a_01308

Antonello, R., Turek, J. S., Vo, V., & Huth, A. (2021). Low-dimensional structure in the space of language representations is reflected in brain responses. In M. Ranzato, A. Beygelzimer, Y. Dauphin, P. S. Liang, & J. W. Vaughan (Eds.), Advances in Neural Information Processing Systems (Vol. 34, pp. 8332–8344). Curran Associates, Inc. https://proceedings.neurips.cc/paper/2021/file/464074179972cbbd75a39abc6954cd12-Paper.pdf

Baldassano, C., Hasson, U., & Norman, K. A. (2018). Representation of real-world event schemas during narrative perception. Journal of Neuroscience, 38(45), 9689–9699. https://doi.org/10.1523/JNEUROSCI.0251-18.2018

Baroni, M. (2020). Linguistic generalization and compositionality in modern artificial neural networks. *Philosophical Transactions of the Royal Society of London: Series B*, Biological Sciences, 375(1791), 20190307. https://doi.org/10.1098/rstb.2019.0307

Bašnáková, J., Weber, K., Petersson, K. M., van Berkum, J., & Hagoort, P. (2014). Beyond the language given: the neural correlates of inferring speaker meaning. Cerebral Cortex, 24(10), 2572–2578. https://doi.org/10.1093/cercor/bht112

Behzadi, Y., Restom, K., Liau, J., & Liu, T. T. (2007). A component based noise correction method (CompCor) for BOLD and perfusion based fMRI. NeuroImage, 37(1), 90–101. https://doi.org/10.1016/j.neuroimage.2007.04.042

Benjamini, Y., & Hochberg, Y. (1995). Controlling the false discovery rate: a practical and powerful approach to multiple testing. *Journal of the Royal Statistical Society: Series B*, Statistical Methodology, 57(1), 289–300. http://www.jstor.org/stable/2346101

Ben-Shachar, M., Hendler, T., Kahn, I., Ben-Bashat, D., & Grodzinsky, Y. (2003). The neural reality of syntactic transformations: evidence from functional magnetic resonance imaging. Psychological Science, 14(5), 433–440. https://doi.org/10.1111/1467-9280.01459

Berwick, R. C., Friederici, A. D., Chomsky, N., & Bolhuis, J. J. (2013). Evolution, brain, and the nature of language. Trends in Cognitive Sciences, 17(2), 89–98. https://doi.org/10.1016/j.tics.2012.12.002

Binder, J. R., Desai, R. H., Graves, W. W., & Conant, L. L. (2009). Where is the semantic system? A critical review and meta-analysis of 120 functional neuroimaging studies. Cerebral Cortex, 19(12), 2767–2796. https://doi.org/10.1093/cercor/bhp055

Blank, I., Balewski, Z., Mahowald, K., & Fedorenko, E. (2016). Syntactic processing is distributed across the language system. NeuroImage, 127, 307–323. https://doi.org/10.1016/j.neuroimage.2015.11.069

Bookheimer, S. (2002). Functional MRI of language: new approaches to understanding the cortical organization of semantic processing. Annual Review of Neuroscience, 25, 151–188. https://doi.org/10.1146/annurev.neuro.25.112701.142946

Bornkessel, I., Zysset, S., Friederici, A. D., von Cramon, D. Y., & Schlesewsky, M. (2005). Who did what to whom? The neural basis of argument hierarchies during language comprehension. NeuroImage, 26(1), 221–233. https://doi.org/10.1016/j.neuroimage.2005.01.032

Braga, R. M., DiNicola, L. M., Becker, H. C., & Buckner, R. L. (2020). Situating the left-lateralized language network in the broader organization of multiple specialized large-scale distributed networks. Journal of Neurophysiology, 124(5), 1415–1448. https://doi.org/10.1152/jn.00753.2019

Brodersen, K. H., Ong, C. S., Stephan, K. E., & Buhmann, J. M. (2010). The balanced accuracy and its posterior distribution. 2010 20th International Conference on Pattern Recognition, 3121–3124. https://doi.org/10.1109/ICPR.2010.764

Bruner, J. S. (1985). Actual Minds, Possible Worlds. Harvard University Press.

Carden, G. (1982). Backwards anaphora in discourse context. Journal of Linguistics, 18(2), 361–387. https://doi.org/10.1017/S0022226700013657

Catani, M., Jones, D. K., & Ffytche, D. H. (2005). Perisylvian language networks of the human brain. Annals of Neurology, 57(1), 8–16. https://doi.org/10.1002/ana.20319

Caucheteux, C., Gramfort, A., & King, J.-R. (2021a). Model-based analysis of brain activity reveals the hierarchy of language in 305 subjects. arXiv. http://arxiv.org/abs/2110.06078

Caucheteux, C., Gramfort, A., & King, J.-R. (2021b). Long-range and hierarchical language predictions in brains and algorithms. arXiv. http://arxiv.org/abs/2111.14232

Caucheteux, C., Gramfort, A., & King, J.-R. (2022). Deep language algorithms predict semantic comprehension from brain activity. Scientific Reports, 12(1), 16327. https://doi.org/10.1038/s41598-022-20460-9

Caucheteux, C., Gramfort, A., & King, J.-R. (2021c). Disentangling syntax and semantics in the brain with deep networks. In M. Meila & T. Zhang (Eds.), Proceedings of the 38th International Conference on Machine Learning (Vol. 139, pp. 1336–1348). PMLR. https://proceedings.mlr.press/v139/caucheteux21a.html

Caucheteux, C., & King, J.-R. (2022). Brains and algorithms partially converge in natural language processing. Communications Biology, 5(1), 134. https://doi.org/10.1038/s42003-022-03036-1

Chang, C. H. C., Nastase, S. A., & Hasson, U. (2022). Information flow across the cortical timescale hierarchy during narrative construction. Proceedings of the National Academy of Sciences of the United States of America, 119(51), e2209307119. https://doi.org/10.1073/pnas.2209307119

Chomsky, N. (1965). Aspects of the Theory of Syntax. MIT Press.

Christiansen, M. H., & Chater, N. (2016). The now-or-never bottleneck: a fundamental constraint on language. Behavioral and Brain Sciences, 39, e62. https://doi.org/10.1017/S0140525X1500031X

Clark, K., Khandelwal, U., Levy, O., & Manning, C. D. (2019). What does BERT look at? An analysis of BERT’s attention. Proceedings of the 2019 ACL Workshop BlackboxNLP: Analyzing and Interpreting Neural Networks for NLP, 276–286. https://doi.org/10.18653/v1/W19-4828

Cox, R. W. (1996). AFNI: software for analysis and visualization of functional magnetic resonance neuroimages. Computers and Biomedical Research, an International Journal, 29(3), 162–173. https://doi.org/10.1006/cbmr.1996.0014

Dapretto, M., & Bookheimer, S. Y. (1999). Form and content: dissociating syntax and semantics in sentence comprehension. Neuron, 24(2), 427–432. https://doi.org/10.1016/s0896-6273(00)80855-7

de Heer, W. A., Huth, A. G., Griffiths, T. L., Gallant, J. L., & Theunissen, F. E. (2017). The hierarchical cortical organization of human speech processing. Journal of Neuroscience, 37(27), 6539–6557. https://doi.org/10.1523/JNEUROSCI.3267-16.2017

DeRose, J. F., Wang, J., & Berger, M. (2020). Attention flows: analyzing and comparing attention mechanisms in language models. arXiv. http://arxiv.org/abs/2009.07053

Devlin, J., Chang, M.-W., Lee, K., & Toutanova, K. (2019). BERT: pre-training of deep bidirectional transformers for language understanding. Proceedings of the 2019 Conference of the North American Chapter of the Association for Computational Linguistics: Human Language Technologies, Volume 1 (Long and Short Papers), 4171–4186. https://doi.org/10.18653/v1/N19-1423

Dick, A. S., & Tremblay, P. (2012). Beyond the arcuate fasciculus: consensus and controversy in the connectional anatomy of language. Brain, 135(Pt 12), 3529–3550. https://doi.org/10.1093/brain/aws222

Ding, N., Melloni, L., Zhang, H., Tian, X., & Poeppel, D. (2016). Cortical tracking of hierarchical linguistic structures in connected speech. Nature Neuroscience, 19(1), 158–164. https://doi.org/10.1038/nn.4186

Dobs, K., Martinez, J., Kell, A. J. E., & Kanwisher, N. (2022). Brain-like functional specialization emerges spontaneously in deep neural networks. Science Advances, 8(11), eabl8913. https://doi.org/10.1126/sciadv.abl8913

Dupré la Tour, T., Eickenberg, M., Nunez-Elizalde, A. O., & Gallant, J. L. (2022). Feature-space selection with banded ridge regression. NeuroImage, 264, 119728. https://doi.org/10.1016/j.neuroimage.2022.119728

Dupré la Tour, T., Lu, M., Eickenberg, M., & Gallant, J. L. (2021). A finer mapping of convolutional neural network layers to the visual cortex. SVRHM 2021 Workshop @ NeurIPS. https://openreview.net/pdf?id=EcoKpq43Ul8

Elhage, N., Nanda, N., Olsson, C., Henighan, T., Joseph, N., Mann, B., Askell, A., Bai, Y., Chen, A., Conerly, T., DasSarma, N., Drain, D., Ganguli, D., Hatfield-Dodds, Z., Hernandez, D., Jones, A., Kernion, J., Lovitt, L., Ndousse, K., … Olah, C. (2021). A Mathematical Framework for Transformer Circuits. Transformer Circuits Thread.

Elman, J. L. (1990). Finding structure in time. Cognitive Science, 14(2), 179–211. https://doi.org/10.1207/s15516709cog1402_1

Embick, D., Marantz, A., Miyashita, Y., O’Neil, W., & Sakai, K. L. (2000). A syntactic specialization for Broca’s area. Proceedings of the National Academy of Sciences of the United States of America, 97(11), 6150–6154. https://doi.org/10.1073/pnas.100098897

Esteban, O., Markiewicz, C. J., Blair, R. W., Moodie, C. A., Ilkay Isik, A., Erramuzpe, A., Kent, J. D., Goncalves, M., DuPre, E., Snyder, M., Oya, H., Ghosh, S. S., Wright, J., Durnez, J., Poldrack, R. A., & Gorgolewski, K. J. (2019). fMRIPrep: a robust preprocessing pipeline for functional MRI. In Nature Methods (Vol. 16, Issue 1, pp. 111–116). https://doi.org/10.1038/s41592-018-0235-4

Fedorenko, E., Behr, M. K., & Kanwisher, N. (2011). Functional specificity for high-level linguistic processing in the human brain. Proceedings of the National Academy of Sciences of the United States of America, 108(39), 16428–16433. https://doi.org/10.1073/pnas.1112937108

Fedorenko, E., & Blank, I. A. (2020). Broca’s area is not a natural kind. Trends in Cognitive Sciences, 24(4), 270–284. https://doi.org/10.1016/j.tics.2020.01.001

Fedorenko, E., Blank, I. A., Siegelman, M., & Mineroff, Z. (2020). Lack of selectivity for syntax relative to word meanings throughout the language network. Cognition, 203, 104348. https://doi.org/10.1016/j.cognition.2020.104348

Fedorenko, E., Hsieh, P.-J., Nieto-Castañón, A., Whitfield-Gabrieli, S., & Kanwisher, N. (2010). New method for fMRI investigations of language: defining ROIs functionally in individual subjects. Journal of Neurophysiology, 104(2), 1177–1194. https://doi.org/10.1152/jn.00032.2010

Fedorenko, E., Nieto-Castañon, A., & Kanwisher, N. (2012). Lexical and syntactic representations in the brain: an fMRI investigation with multi-voxel pattern analyses. Neuropsychologia, 50(4), 499–513. https://doi.org/10.1016/j.neuropsychologia.2011.09.014

Ferstl, E. C., Neumann, J., Bogler, C., & von Cramon, D. Y. (2008). The extended language network: a meta-analysis of neuroimaging studies on text comprehension. Human Brain Mapping, 29(5), 581–593. https://doi.org/10.1002/hbm.20422

Flick, G., & Pylkkänen, L. (2020). Isolating syntax in natural language: MEG evidence for an early contribution of left posterior temporal cortex. Cortex, 127, 42–57. https://doi.org/10.1016/j.cortex.2020.01.025

Friederici, A. D. (2011). The brain basis of language processing: from structure to function. Physiological Reviews, 91(4), 1357–1392. https://doi.org/10.1152/physrev.00006.2011

Friederici, A. D., Chomsky, N., Berwick, R. C., Moro, A., & Bolhuis, J. J. (2017). Language, mind and brain. Nature Human Behaviour, 1(10), 713–722. https://doi.org/10.1038/s41562-017-0184-4

Friederici, A. D., Rüschemeyer, S.-A., Hahne, A., & Fiebach, C. J. (2003). The role of left inferior frontal and superior temporal cortex in sentence comprehension: localizing syntactic and semantic processes. Cerebral Cortex, 13(2), 170–177. https://doi.org/10.1093/cercor/13.2.170

Glaser, Y. G., Martin, R. C., Van Dyke, J. A., Hamilton, A. C., & Tan, Y. (2013). Neural basis of semantic and syntactic interference in sentence comprehension. Brain and Language, 126(3), 314–326. https://doi.org/10.1016/j.bandl.2013.06.006

Goldberg, A. E. (2006). Constructions at Work: The Nature of Generalization in Language. Oxford University Press.

Goldstein, A., Ham, E., Nastase, S. A., Zada, Z., Grinstein-Dabus, A., Aubrey, B., Schain, M., Gazula, H., Feder, A., Doyle, W., Devore, S., Dugan, P., Friedman, D., Brenner, M., Hassidim, A., Devinsky, O., Flinker, A., Levy, O., & Hasson, U. (2022). Correspondence between the layered structure of deep language models and temporal structure of natural language processing in the human brain. bioRxiv. https://doi.org/10.1101/2022.07.11.499562

Goldstein, A., Zada, Z., Buchnik, E., Schain, M., Price, A., Aubrey, B., Nastase, S. A., Feder, A., Emanuel, D., Cohen, A., Jansen, A., Gazula, H., Choe, G., Rao, A., Kim, C., Casto, C., Fanda, L., Doyle, W., Friedman, D., … Hasson, U. (2022). Shared computational principles for language processing in humans and deep language models. Nature Neuroscience, 25(3), 369–380. https://doi.org/10.1038/s41593-022-01026-4

Gorgolewski, K. J., Auer, T., Calhoun, V. D., Craddock, R. C., Das, S., Duff, E. P., Flandin, G., Ghosh, S. S., Glatard, T., Halchenko, Y. O., Handwerker, D. A., Hanke, M., Keator, D., Li, X., Michael, Z., Maumet, C., Nichols, B. N., Nichols, T. E., Pellman, J., … Poldrack, R. A. (2016). The brain imaging data structure, a format for organizing and describing outputs of neuroimaging experiments. Scientific Data, 3, 160044. https://doi.org/10.1038/sdata.2016.44

Graesser, A. C., Singer, M., & Trabasso, T. (1994). Constructing inferences during narrative text comprehension. Psychological Review, 101(3), 371–395. https://doi.org/10.1037/0033-295x.101.3.371

Güçlü, U., & van Gerven, M. A. J. (2015). Deep neural networks reveal a gradient in the complexity of neural representations across the ventral stream. Journal of Neuroscience, 35(27), 10005–10014. https://doi.org/10.1523/JNEUROSCI.5023-14.2015

Hall, P., & Wilson, S. R. (1991). Two guidelines for bootstrap hypothesis testing. Biometrics, 47(2), 757–762. https://doi.org/10.2307/2532163

Hamilton, L. S., & Huth, A. G. (2020). The revolution will not be controlled: natural stimuli in speech neuroscience. *Language*, Cognition and Neuroscience, 35(5), 573–582. https://doi.org/10.1080/23273798.2018.1499946

Hasson, U., Chen, J., & Honey, C. J. (2015). Hierarchical process memory: memory as an integral component of information processing. Trends in Cognitive Sciences, 19(6), 304–313. https://doi.org/10.1016/j.tics.2015.04.006

Hasson, U., Nastase, S. A., & Goldstein, A. (2020). Direct fit to nature: an evolutionary perspective on biological and artificial neural networks. Neuron, 105(3), 416–434. https://doi.org/10.1016/j.neuron.2019.12.002

Hawkins, R. D., Yamakoshi, T., Griffiths, T. L., & Goldberg, A. E. (2020). Investigating representations of verb bias in neural language models. arXiv. http://arxiv.org/abs/2010.02375

Heilbron, M., Armeni, K., Schoffelen, J.-M., Hagoort, P., & de Lange, F. P. (2022). A hierarchy of linguistic predictions during natural language comprehension. Proceedings of the National Academy of Sciences of the United States of America, 119(32), e2201968119. https://doi.org/10.1073/pnas.2201968119

Hewitt, J., & Manning, C. D. (2019). A structural probe for finding syntax in word representations. Proceedings of the 2019 Conference of the North American Chapter of the Association for Computational Linguistics: Human Language Technologies, Volume 1 (Long and Short Papers), 4129–4138. https://www.aclweb.org/anthology/N19-1419.pdf

He, Zhang, Ren, & Sun. (2016). Deep residual learning for image recognition. Proceedings of the IEEE Conference on Computer Vision and Pattern Recognition (CVPR), 770–778. http://openaccess.thecvf.com/content_cvpr_2016/html/He_Deep_Residual_Learning_CVPR_2016_paper.html

Hickok, G., & Poeppel, D. (2000). Towards a functional neuroanatomy of speech perception. Trends in Cognitive Sciences, 4(4), 131–138. https://doi.org/10.1016/s1364-6613(00)01463-7

Hickok, G., & Poeppel, D. (2007). The cortical organization of speech processing. Nature Reviews. Neuroscience, 8(5), 393–402. https://doi.org/10.1038/nrn2113

Hochreiter, S., & Schmidhuber, J. (1997). Long short-term memory. Neural Computation, 9(8), 1735–1780. https://doi.org/10.1162/neco.1997.9.8.1735

Honnibal, M., Montani, I., Van Landeghem, S., & Boyd, A. (2020). SpaCy: industrial-strength natural language processing in python. Zenodo.

Hoover, B., Strobelt, H., & Gehrmann, S. (2020). exBERT: a visual analysis tool to explore learned representations in transformer models. Proceedings of the 58th Annual Meeting of the Association for Computational Linguistics: System Demonstrations, 187–196. https://doi.org/10.18653/v1/2020.acl-demos.22

Huth, A. G., de Heer, W. A., Griffiths, T. L., Theunissen, F. E., & Gallant, J. L. (2016). Natural speech reveals the semantic maps that tile human cerebral cortex. Nature, 532(7600), 453–458. https://doi.org/10.1038/nature17637

Huth, A. G., Nishimoto, S., Vu, A. T., & Gallant, J. L. (2012). A continuous semantic space describes the representation of thousands of object and action categories across the human brain. Neuron, 76(6), 1210–1224. https://doi.org/10.1016/j.neuron.2012.10.014

Jain, S., & Huth, A. (2018). Incorporating context into language encoding models for fMRI. In S. Bengio, H. Wallach, H. Larochelle, K. Grauman, N. Cesa-Bianchi, & R. Garnett (Eds.), Advances in Neural Information Processing Systems (Vol. 31, pp. 6628–6637). Curran Associates, Inc. http://papers.nips.cc/paper/7897-incorporating-context-into-language-encoding-models-for-fmri.pdf

Kriegeskorte, N. (2015). Deep neural networks: a new framework for modeling biological vision and brain information processing. Annual Review of Vision Science, 1, 417–446. https://doi.org/10.1146/annurev-vision-082114-035447

Krizhevsky, A., Sutskever, I., & Hinton, G. E. (2012). ImageNet classification with deep convolutional neural networks. In F. Pereira, C. J. Burges, L. Bottou, & K. Q. Weinberger (Eds.), Advances in Neural Information Processing Systems (Vol. 25, pp. 1097–1105). Curran Associates, Inc. https://proceedings.neurips.cc/paper/4824-imagenet-classification-with-deep-convolutional-neural-netw orks.pdf

Kuperberg, G. R., McGuire, P. K., Bullmore, E. T., Brammer, M. J., Rabe-Hesketh, S., Wright, I. C., Lythgoe, D. J., Williams, S. C., & David, A. S. (2000). Common and distinct neural substrates for pragmatic, semantic, and syntactic processing of spoken sentences: an fMRI study. Journal of Cognitive Neuroscience, 12(2), 321–341. https://doi.org/10.1162/089892900562138

Landauer, T. K., & Dumais, S. T. (1997). A solution to Plato’s problem: the latent semantic analysis theory of acquisition, induction, and representation of knowledge. Psychological Review, 104(2), 211–240. https://doi.org/10.1037/0033-295X.104.2.211

LeCun, Bottou, L., Bengio, Y., & Haffner, P. (1998). Gradient-based learning applied to document recognition. Proceedings of the IEEE, 86(11), 2278–2324. https://doi.org/10.1109/5.726791

Lee Masson, H., & Isik, L. (2021). Functional selectivity for social interaction perception in the human superior temporal sulcus during natural viewing. NeuroImage, 245, 118741. https://doi.org/10.1016/j.neuroimage.2021.118741

Lerner, Y., Honey, C. J., Silbert, L. J., & Hasson, U. (2011). Topographic mapping of a hierarchy of temporal receptive windows using a narrated story. Journal of Neuroscience, 31(8), 2906–2915. https://doi.org/10.1523/JNEUROSCI.3684-10.2011

Linzen, T., & Baroni, M. (2021). Syntactic structure from deep learning. Annual Review of Linguistics, 7(1), 195–212. https://doi.org/10.1146/annurev-linguistics-032020-051035

Liu, N. F., Gardner, M., Belinkov, Y., Peters, M. E., & Smith, N. A. (2019). Linguistic knowledge and transferability of contextual representations. Proceedings of the 2019 Conference of the North American Chapter of the Association for Computational Linguistics: Human Language Technologies, Volume 1 (Long and Short Papers), 1073–1094. https://doi.org/10.18653/v1/N19-1112

Li, Y., Anumanchipalli, G. K., Mohamed, A., Lu, J., Wu, J., & Chang, E. F. (2022). Dissecting neural computations of the human auditory pathway using deep neural networks for speech. bioRxiv. https://doi.org/10.1101/2022.03.14.484195

Lyu, B., Marslen-Wilson, W. D., Fang, Y., & Tyler, L. K. (2021). Finding structure in time: humans, machines, and language. bioRxiv. https://doi.org/10.1101/2021.10.25.465687

MacDonald, M. C., Pearlmutter, N. J., & Seidenberg, M. S. (1994). The lexical nature of syntactic ambiguity resolution. Psychological Review, 101(4), 676–703. https://doi.org/10.1037/0033-295x.101.4.676

Maguire, E. A., Frith, C. D., & Morris, R. G. (1999). The functional neuroanatomy of comprehension and memory: the importance of prior knowledge. Brain, 122 ( *Pt 10*), 1839–1850. https://doi.org/10.1093/brain/122.10.1839

Makuuchi, M., Bahlmann, J., Anwander, A., & Friederici, A. D. (2009). Segregating the core computational faculty of human language from working memory. Proceedings of the National Academy of Sciences of the United States of America, 106(20), 8362–8367. https://doi.org/10.1073/pnas.0810928106

Manning, C. D., Clark, K., Hewitt, J., Khandelwal, U., & Levy, O. (2020). Emergent linguistic structure in artificial neural networks trained by self-supervision. Proceedings of the National Academy of Sciences of the United States of America, 117(48), 30046–30054. https://doi.org/10.1073/pnas.1907367117

Martin, A. E. (2020). A compositional neural architecture for language. Journal of Cognitive Neuroscience, 32(8), 1407–1427. https://doi.org/10.1162/jocn_a_01552

Martin, A. E., & Doumas, L. A. A. (2017). A mechanism for the cortical computation of hierarchical linguistic structure. PLoS Biology, 15(3), e2000663. https://doi.org/10.1371/journal.pbio.2000663

Matchin, W., & Hickok, G. (2020). The cortical organization of syntax. Cerebral Cortex, 30(3), 1481–1498. https://doi.org/10.1093/cercor/bhz180

McClelland, J. L., Botvinick, M. M., Noelle, D. C., Plaut, D. C., Rogers, T. T., Seidenberg, M. S., & Smith, L. B. (2010). Letting structure emerge: connectionist and dynamical systems approaches to cognition. Trends in Cognitive Sciences, 14(8), 348–356. https://doi.org/10.1016/j.tics.2010.06.002

Meng, K., Bau, D., Andonian, A., & Belinkov, Y. (2022). Locating and editing factual associations in GPT. In S. Koyejo, S. Mohamed, A. Agarwal, D. Belgrave, K. Cho, & A. Oh (Eds.), Advances in Neural Information Processing Systems (Vol. 35) (pp. 17359–17372). Curran Associates, Inc. https://proceedings.neurips.cc/paper_files/paper/2022/file/6f1d43d5a82a37e89b0665b33bf3a182-Paper-Conference.pdf

Mesulam, M.-M., Thompson, C. K., Weintraub, S., & Rogalski, E. J. (2015). The Wernicke conundrum and the anatomy of language comprehension in primary progressive aphasia. Brain, 138(Pt 8), 2423–2437. https://doi.org/10.1093/brain/awv154

Mikolov, T., Sutskever, I., Chen, K., Corrado, G. S., & Dean, J. (2013). Distributed representations of words and phrases and their compositionality. In C. J. Burges, L. Bottou, M. Welling, Z. Ghahramani, & K. Q. Weinberger (Eds.), Advances in Neural Information Processing Systems (Vol. 26). Curran Associates, Inc. https://proceedings.neurips.cc/paper/2013/file/9aa42b31882ec039965f3c4923ce901b-Paper.pdf

Millet, J., Caucheteux, C., Orhan, P., Boubenec, Y., Gramfort, A., Dunbar, E., Pallier, C., & King, J.-R. (2022). Toward a realistic model of speech processing in the brain with self-supervised learning. arXiv. http://arxiv.org/abs/2206.01685

Mineroff, Z., Blank, I. A., Mahowald, K., & Fedorenko, E. (2018). A robust dissociation among the language, multiple demand, and default mode networks: evidence from inter-region correlations in effect size. Neuropsychologia, 119, 501–511. https://doi.org/10.1016/j.neuropsychologia.2018.09.011

Mitchell, T. M., Shinkareva, S. V., Carlson, A., Chang, K.-M., Malave, V. L., Mason, R. A., & Just, M. A. (2008). Predicting human brain activity associated with the meanings of nouns. Science, 320(5880), 1191–1195. https://doi.org/10.1126/science.1152876

Murphy, E., Woolnough, O., Rollo, P. S., Roccaforte, Z. J., Segaert, K., Hagoort, P., & Tandon, N. (2022). Minimal phrase composition revealed by intracranial recordings. Journal of Neuroscience, 42(15), 3216–3227. https://doi.org/10.1523/JNEUROSCI.1575-21.2022

Naselaris, T., Kay, K. N., Nishimoto, S., & Gallant, J. L. (2011). Encoding and decoding in fMRI. NeuroImage, 56(2), 400–410. https://doi.org/10.1016/j.neuroimage.2010.07.073

Nasr, K., Viswanathan, P., & Nieder, A. (2019). Number detectors spontaneously emerge in a deep neural network designed for visual object recognition. Science Advances, 5(5). https://doi.org/10.1126/sciadv.aav7903

Nastase, S. A., Gazzola, V., Hasson, U., & Keysers, C. (2019). Measuring shared responses across subjects using intersubject correlation. Social Cognitive and Affective Neuroscience, 14(6), 667–685. https://doi.org/10.1093/scan/nsz037

Nastase, S. A., Goldstein, A., & Hasson, U. (2020). Keep it real: rethinking the primacy of experimental control in cognitive neuroscience. NeuroImage, 222, 117254. https://doi.org/10.1016/j.neuroimage.2020.117254

Nastase, S. A., Liu, Y.-F., Hillman, H., Zadbood, A., Hasenfratz, L., Keshavarzian, N., Chen, J., Honey, C. J., Yeshurun, Y., Regev, M., Nguyen, M., Chang, C. H. C., Baldassano, C., Lositsky, O., Simony, E., Chow, M. A., Leong, Y. C., Brooks, P. P., Micciche, E., … Hasson, U. (2021). The “Narratives” fMRI dataset for evaluating models of naturalistic language comprehension. Scientific Data, 8(1), 250. https://doi.org/10.1038/s41597-021-01033-3

Nili, H., Wingfield, C., Walther, A., Su, L., Marslen-Wilson, W., & Kriegeskorte, N. (2014). A toolbox for representational similarity analysis. PLoS Computational Biology, 10(4), e1003553. https://doi.org/10.1371/journal.pcbi.1003553

Ni, W., Constable, R. T., Mencl, W. E., Pugh, K. R., Fulbright, R. K., Shaywitz, S. E., Shaywitz, B. A., Gore, J. C., & Shankweiler, D. (2000). An event-related neuroimaging study distinguishing form and content in sentence processing. Journal of Cognitive Neuroscience, 12(1), 120–133. https://doi.org/10.1162/08989290051137648

Nunez-Elizalde, A. O., Huth, A. G., & Gallant, J. L. (2019). Voxelwise encoding models with non-spherical multivariate normal priors. NeuroImage, 197, 482–492. https://doi.org/10.1016/j.neuroimage.2019.04.012

Partee, B. (1995). Lexical semantics and compositionality. An Invitation to Cognitive Science: Language, 1, 311–360.

Pavlick, E. (2022). Semantic structure in deep learning. Annual Review of Applied Linguistics, 8, 447–471. https://doi.org/10.1146/annurev-linguistics-031120-122924

Pedregosa, F., Varoquaux, G., Gramfort, A., Michel, V., Thirion, B., Grisel, O., Blondel, M., Prettenhofer, P., Weiss, R., Dubourg, V., Vanderplas, J., Passos, A., Cournapeau, D., Brucher, M., Perrot, M., & Duchesnay, É. (2011). Scikit-learn: machine learning in Python. Journal of Machine Learning Research: JMLR, 12(85), 2825–2830. http://www.jmlr.org/papers/v12/pedregosa11a.html

Pennington, J., Socher, R., & Manning, C. (2014). GloVe: Global vectors for word representation. Proceedings of the 2014 Conference on Empirical Methods in Natural Language Processing (EMNLP), 1532–1543. https://doi.org/10.3115/v1/d14-1162

Pereira, F., Lou, B., Pritchett, B., Ritter, S., Gershman, S. J., Kanwisher, N., Botvinick, M., & Fedorenko, E. (2018). Toward a universal decoder of linguistic meaning from brain activation. Nature Communications, 9(1), 963. https://doi.org/10.1038/s41467-018-03068-4

Price, C. J. (2012). A review and synthesis of the first 20 years of PET and fMRI studies of heard speech, spoken language and reading. NeuroImage, 62(2), 816–847. https://doi.org/10.1016/j.neuroimage.2012.04.062

Pylkkänen, L. (2019). The neural basis of combinatory syntax and semantics. Science, 366(6461), 62–66. https://doi.org/10.1126/science.aax0050

Radford, A., Wu, J., Child, R., Luan, D., Amodei, D., & Sutskever, I. (2019). Language models are unsupervised multitask learners. OpenAI Blog. https://www.techbooky.com/wp-content/uploads/2019/02/Better-Language-Models-and-Their-Implications.pdf

Raghu, M., Unterthiner, T., Kornblith, S., Zhang, C., & Dosovitskiy, A. (2021). Do vision transformers see like convolutional neural networks? In M. Ranzato, A. Beygelzimer, Y. Dauphin, P. S. Liang, & J. W. Vaughan (Eds.), Advances in Neural Information Processing Systems (Vol. 34, pp. 12116–12128). Curran Associates, Inc. https://proceedings.neurips.cc/paper/2021/file/652cf38361a209088302ba2b8b7f51e0-Paper.pdf

Reddy, A. J., & Wehbe, L. (2020). Can fMRI reveal the representation of syntactic structure in the brain? bioRxiv. https://doi.org/10.1101/2020.06.16.155499

Richards, B. A., Lillicrap, T. P., Beaudoin, P., Bengio, Y., Bogacz, R., Christensen, A., Clopath, C., Costa, R. P., de Berker, A., Ganguli, S., Gillon, C. J., Hafner, D., Kepecs, A., Kriegeskorte, N., Latham, P., Lindsay, G. W., Miller, K. D., Naud, R., Pack, C. C., … Kording, K. P. (2019). A deep learning framework for neuroscience. Nature Neuroscience, 22(11), 1761–1770. https://doi.org/10.1038/s41593-019-0520-2

Rogers, A., Kovaleva, O., & Rumshisky, A. (2020). A primer in BERTology: what we know about how BERT works. Transactions of the Association for Computational Linguistics, 8, 842–866. https://direct.mit.edu/tacl/article-abstract/doi/10.1162/tacl_a_00349/96482

Santoro, R., Moerel, M., De Martino, F., Goebel, R., Ugurbil, K., Yacoub, E., & Formisano, E. (2014). Encoding of natural sounds at multiple spectral and temporal resolutions in the human auditory cortex. PLoS Computational Biology, 10(1), e1003412. https://doi.org/10.1371/journal.pcbi.1003412

Saur, D., Kreher, B. W., Schnell, S., Kümmerer, D., Kellmeyer, P., Vry, M.-S., Umarova, R., Musso, M., Glauche, V., Abel, S., Huber, W., Rijntjes, M., Hennig, J., & Weiller, C. (2008). Ventral and dorsal pathways for language. Proceedings of the National Academy of Sciences of the United States of America, 105(46), 18035–18040. https://doi.org/10.1073/pnas.0805234105

Schaefer, A., Kong, R., Gordon, E. M., Laumann, T. O., Zuo, X.-N., Holmes, A. J., Eickhoff, S. B., & Yeo, B. T. T. (2018). Local-global parcellation of the human cerebral cortex from intrinsic functional connectivity MRI. Cerebral Cortex, 28(9), 3095–3114. https://doi.org/10.1093/cercor/bhx179

Schell, M., Zaccarella, E., & Friederici, A. D. (2017). Differential cortical contribution of syntax and semantics: an fMRI study on two-word phrasal processing. Cortex, 96, 105–120. https://doi.org/10.1016/j.cortex.2017.09.002

Schrimpf, M., Blank, I. A., Tuckute, G., Kauf, C., Hosseini, E. A., Kanwisher, N., Tenenbaum, J. B., & Fedorenko, E. (2021). The neural architecture of language: Integrative modeling converges on predictive processing. Proceedings of the National Academy of Sciences of the United States of America, 118(45). https://doi.org/10.1073/pnas.2105646118

Tenney, I., Das, D., & Pavlick, E. (2019). BERT rediscovers the classical NLP pipeline. Proceedings of the 57th Annual Meeting of the Association for Computational Linguistics, 4593–4601. https://doi.org/10.18653/v1/P19-1452

Toneva, M., & Wehbe, L. (2019). Interpreting and improving natural-language processing (in machines) with natural language-processing (in the brain). In H. Wallach, H. Larochelle, A. Beygelzimer, F. d\textquotesingle Alché-Buc, E. Fox, & R. Garnett (Eds.), Advances in Neural Information Processing Systems (Vol. 32, pp. 14954–14964). Curran Associates Inc. https://dl.acm.org/doi/abs/10.5555/3454287.3455626

Vaidya, A. R., Jain, S., & Huth, A. G. (2022). Self-supervised models of audio effectively explain human cortical responses to speech. arXiv. http://arxiv.org/abs/2205.14252

Vandenberghe, R., Nobre, A. C., & Price, C. J. (2002). The response of left temporal cortex to sentences. Journal of Cognitive Neuroscience, 14(4), 550–560. https://doi.org/10.1162/08989290260045800

Vaswani, A., Shazeer, N., Parmar, N., Uszkoreit, J., Jones, L., Gomez, A. N., Kaiser, Ł., & Polosukhin, I. (2017). Attention is all you need. In I. Guyon, U. V. Luxburg, S. Bengio, H. Wallach, R. Fergus, S. Vishwanathan, & R. Garnett (Eds.), Advances in Neural Information Processing Systems (Vol. 30, pp. 6000–6010). Curran Associates Inc. https://dl.acm.org/doi/10.5555/3295222.3295349

Vig, J., & Belinkov, Y. (2019). Analyzing the structure of attention in a transformer language model. Proceedings of the 2019 ACL Workshop BlackboxNLP: Analyzing and Interpreting Neural Networks for NLP, 63–76. https://doi.org/10.18653/v1/W19-4808

Vigneau, M., Beaucousin, V., Hervé, P. Y., Duffau, H., Crivello, F., Houdé, O., Mazoyer, B., & Tzourio-Mazoyer, N. (2006). Meta-analyzing left hemisphere language areas: phonology, semantics, and sentence processing. NeuroImage, 30(4), 1414–1432. https://doi.org/10.1016/j.neuroimage.2005.11.002

Wehbe, L., Murphy, B., Talukdar, P., Fyshe, A., Ramdas, A., & Mitchell, T. (2014). Simultaneously uncovering the patterns of brain regions involved in different story reading subprocesses. PloS One, 9(11), e112575. https://doi.org/10.1371/journal.pone.0112575

Willems, R. M., Nastase, S. A., & Milivojevic, B. (2020). Narratives for neuroscience. Trends in Neurosciences, 43(5), 271–273. https://doi.org/10.1016/j.tins.2020.03.003

Wolf, T., Debut, L., Sanh, V., Chaumond, J., Delangue, C., Moi, A., Cistac, P., Rault, T., Louf, R., Funtowicz, M., Davison, J., Shleifer, S., von Platen, P., Ma, C., Jernite, Y., Plu, J., Xu, C., Le Scao, T., Gugger, S., … Rush, A. (2020). Transformers: state-of-the-art natural language processing. Proceedings of the 2020 Conference on Empirical Methods in Natural Language Processing: System Demonstrations, 38–45. https://doi.org/10.18653/v1/2020.emnlp-demos.6

Yamins, D. L. K., & DiCarlo, J. J. (2016). Using goal-driven deep learning models to understand sensory cortex. Nature Neuroscience, 19(3), 356–365. https://doi.org/10.1038/nn.4244

Yang, G. R., Joglekar, M. R., Song, H. F., Newsome, W. T., & Wang, X.-J. (2019). Task representations in neural networks trained to perform many cognitive tasks. Nature Neuroscience, 22(2), 297–306. https://doi.org/10.1038/s41593-018-0310-2

